# Poly ADP-ribosylation of SET8 leads to aberrant H4K20 methylation in mammalian nuclear genome

**DOI:** 10.1101/2021.11.13.468478

**Authors:** Pierre-Olivier Estève, Sagnik Sen, US Vishnu, Cristian Ruse, Hang Gyeong Chin, Sriharsa Pradhan

**Affiliations:** New England Biolabs Inc, 240 County Road, Ipswich, MA 01938, USA

## Abstract

In mammalian cells, SET8 mediated Histone H4 Lys 20 monomethylation (H4K20me1) has been implicated in regulating mitotic condensation, DNA replication, DNA damage response, and gene expression. Here we show SET8, the only known enzyme for H4K20me1 is post-translationally poly ADP-ribosylated by PARP1 on lysine residues. PARP1 interacts with SET8 in a cell cycle-dependent manner. Poly ADP-ribosylation on SET8 renders it catalytically compromised, and degradation via ubiquitylation pathway. Knockdown of PARP1 led to an increase of SET8 protein levels, leading to aberrant H4K20me1 and H4K20me3 domains genome-wide. H4K20me1 is associated with higher gene transcription levels while the increase of H4K20me3 levels was predominant in DNA repeat elements correlating with modulation of chromatin states. Hence, SET8 mediated chromatin remodeling in mammalian cells are modulated by poly ADP-ribosylation by PARP1.

## Introduction

In the eukaryotic nucleus, double-stranded DNA is wrapped around histone octamers to make chromatin fibers. These chromatin fibers undergo various structural changes during cell division and organism development (1–3). The decompaction and re-establishment of chromatin organization immediately after mitosis are essential for genome regulation (4). Given the absence of membranes separating intranuclear substructures and during cell division, it has been postulated that other structural features of the nucleus, such as the chromatin itself, must impart regulations that would control molecular information flow and stage different physiological activities. Indeed, chromatin accessibility plays a central role in ensuring cell cycle exit and the terminal differentiation during metamorphosis (5). Furthermore, chromatin compaction threshold in cells exiting mitosis ensures genome integrity by limiting replication licensing in G1 phase. Recent studies on histone writer enzymes have shade some lights on the role of histone marks in cell biology (6). The nucleosome structure illustrates that highly basic histone amino (N)- terminal tails can protrude from the core and would be in direct contact with adjacent nucleosomes. Therefore, modification of these histone tails would affect inter-nucleosomal interactions, and thus would modulate the overall chromatin 3D-structure and compaction. Recent advances in Hi-C studies support that this is indeed the case, where transcriptionally active A and inactive B compartments are differentially positioned in the nucleus (7). Histone modifications not only regulate chromatin structure by merely being there, but they also recruit chromatin readers and remodeling enzymes that utilizes the energy derived from the hydrolysis of ATP to reposition nucleosomes (6). For example, lysine acetylation of histone tails is dynamic and regulated by the opposing action of two families of enzymes, histone acetyltransferases (HATs) for acetylation, and histone deacetylases (HDACs) to remove the acetyl groups (8). Indeed, acetylated histones on the chromatin are a cue to transcriptional gene activation (9). Therefore, the balancing act between both HATs and HDACs is essential in maintaining the dynamic equilibrium during gene expression.

The other significant chromatin mark, histone methylation, primarily occurs on the side chains of lysines and arginines (10). Histone methylation does not alter the charge of the histone protein, unlike phosphorylation. However, the epsilon amino group of lysines may be modified to mono-, di- or tri-methylated configuration, whereas arginine may be mono-, symmetrically or asymmetrically di-methylated, creating an unprecedented array of complexity for the reader proteins. These core modifications are established and propagated by SET domain containing enzymes (11). One such enzyme is SET8 (also known as KMT5A, PR-Set7 and SETD8), the only histone methyltransferase that monomethylates histone H4K20 (H4K20me1) (12, 13). Subsequent modification of H4K20me1 by Suv4-20h1/h2 leads to the transition from H4K20me1 to H4K20me2/3 (14). Although H4K20 specific demethylases are less understood, PHF8 is the only known enzyme that demethylates H4K20me1 to H4K20 (15). Similarly, Rad23 has also been shown to demethylate H4K20me1/2/3 (16). In the mouse genome, H4K20 methylation state is distributed in specific regions and is bound by specific reader proteins. Histone H4K20me1 has been implicated in regulating diverse processes ranging from the DNA damage response, mitotic condensation, DNA replication and gene regulation. Indeed, loss of the SET8 causes a more severe and complex phenotype, as this negates the catalysis of all levels of H4K20 methylation. Loss of SET8 causes lethality at the third instar larval stage in Drosophila (17), and in mice it leads to embryonic lethality via developmental arrest between the four- and eight-cell stages (12). Rescue experiments on mouse SET8-/- phenotype embryos by reintroduction of either catalytically active or inactive allele have demonstrated that the catalytic activity of the enzyme is essential for embryo development.

SET8 protein expression is tightly regulated during the cell cycle; it is highest during G2/M and early G1 and is absent during the S phase. Indeed, upon mitotic exit, chromatin relaxation is controlled by SET8-dependent methylation of histone H4K20. In the absence of either SET8 or H4K20me1, substantial genome-wide chromatin decompaction occurs allowing excessive loading of the origin recognition complex (ORC) in the daughter cells (18). During cell cycle, SET8 undergoes ubiquitination and phosphorylation. SET8 is a direct substrate of the E3 ubiquitin ligase complex CRL4Cdt2 (19–21). In addition, SET8 is also regulated by the E3 ubiquitin ligase SCF/Skp2 (22). Skp2 degrades substrates during S and G2 phase and may partially contribute to the steep decrease in SET8 levels during the S phase (23). The other post-translational modification of SET8 that may affect its activity or stability is serine phosphorylation, as discovered by proteomics studies (24–26). SET8 S29 is phosphorylated in vivo by Cyclin B/cdk1 during mitosis, and it causes SET8 to dissociate from mitotic chromosomes at anaphase and relocate to the extrachromosomal space, where it is dephosphorylated by cdc14 phosphatase and subsequently subjected to ubiquitination by APC/Cdh1 (27). Therefore, the coordination of ubiquitylation and phosphorylation processes may be necessary to maintain precise levels of H4K20me1 and SET8 in the genome.

Apart from acetylation, methylation, phosphorylation, and ubiquitylation, poly ADP-ribosylation is another post-translational modification of proteins catalyzed by PARP family of enzymes by the addition of linear or branched chains of ADP-ribose units, from NAD+ substrate (28). The central enzyme for poly ADP-ribosylation in cells during DNA damage is poly-ADP-ribose polymerase 1 (PARP1). Multiple different amino acids are shown to be acceptors of PAR, such as Lys, Arg, Glu, Asp, Cys, Ser, Thr, pSer (phospho-serine, through the phosphate group), although His and Tyr residues were also proposed by proteomic approaches (29–35). PARP family members modify many proteins by poly and/or mono ADP-ribosylation (36). PARPs catalyze short or long, branched, or linear ribosylation. The structural diversity of poly ADP-ribosylation on these proteins could be recognized by poly ADP-ribosylation readers (or binders) with their specific binding motifs for biological function.

In a proteomics study of SET8 binding proteins, we found PARP1 as a strong binder (Table S1). This led us to study the nature of SET8-PARP1 interaction and the role of poly ADP-ribosylation on the catalytic activity of enzymes. Here, we have also investigated the role of poly ADP-ribosylation in SET8 degradation, chromatin remodeling and aberrant H4K20 methylation in mammalian cells.

## Materials and Methods

### XX

#### Cell culture, transfections, and cell cycle synchronization

HeLa, HCT116 and COS-7 cells were cultured in DMEM supplemented with 10% fetal bovine serum according to ATCC’s recommendations. For PARP1 overexpression, HeLa cells were transfected with 500 ng of 3xFLAG or 3xFLAG-PARP1 plasmids using Fugene HD transfection reagent (Promega, # E2311) according to manufacturer’s recommendations. Cells were harvested after 48 hrs. For PARP1 and SET8 knockdown, HeLa cells were transfected with 10 nM of esiRNA targeting either PARP1 (Millipore-Sigma # EHU050101) or SET8 (Millipore-Sigma # EHU111111) using HiPerfect reagent according to manufacturer’s protocol (Qiagen # 301704). 10 nM of esiRNA against EGFP (Millipore-Sigma # EHUEGFP) was used as control and cells were harvested after 48 hrs.

For cell cycle studies, HeLa cells were synchronized using 2 mM thymidine for 24 hrs, washed once with media and released for 8 hours with 24 mM dCTP (New England Biolabs # N0441S). 2 mM of thymidine or 0.1 mg/ml of Nocodazole (Millipore-Sigma # SML1665) were added for 14-15 hrs to arrest the cells in S or G2/M phase respectively. Cells were synchronized in G1 phase using 20 mM of Lovastatin (Tocris # 1530) for 24 hrs. Chromatin was extracted using CSK buffer (10 mM Pipes pH 6.8, 300 mM sucrose, 100 mM NaCl, 1.5 mM MgCl2 and 0.5% Triton X-100 supplemented with protease cocktail inhibitors and PMSF) for 30 min on ice. Cells were centrifuged at 2, 000 g at 4°C for 5 min, washed once with CSK buffer and the pellet (chromatin) was resuspended in buffer containing 50 mM Tris.Cl, pH 7.5, 200 mM NaCl and 1% SDS. Chromatin was then sonicated (10 cycles of 10 pulses) and used either as total chromatin extract or SET8 immunoprecipitation.

#### Western Blot and Densitometry

Western blots were performed as previously described (37). Antibodies against SET8 were obtained from Cell Signaling Technology (# 2996), ABCAM (# ab3798) and Santa Cruz Biotechnology (# sc-515433). Anti-PARP1 (# 9532) and anti-ADPribose antibody (# 83732S) were purchased from Cell Signaling Technology. Anti-H4K20me1 (# MA5-18067), anti-histone H4K20me2 (# 9759) and anti-H4K20me3 (# 5737) were obtained from Thermo Fisher Scientific and Cell Signaling Technology respectively and used at 1/1000 dilution.

Anti-β-actin (# 4970S) and anti-GFP (# 11814460001) antibodies were obtained from Cell Signaling Technology and Millipore-Sigma respectively and used at a 1:5000 dilution. Densitometry was performed using ImageJ software (W. S. Rasband, National Institutes of Health). All densitometry values (arbitrary units) were normalized to either their respective β-actin or Ponceau S staining. ADPribose and ubiquitin quantification levels were normalized to immunoprecipitated SET8 protein levels.

#### In vivo Ubiquitination Assay

COS-7 cells were co-transfected with a mixture of GFP, GFP-SET8FL or GFP-SET8KA, 3xFLAG or 3xFLAG-PARP1, HA-ubiquitin plasmids, and Fugene HD transfection reagent (Promega, # E2311) according to manufacturer’s recommendations. After 48 hrs, transfected COS-7 cells were treated or not with 50 μM MG132 for 2 h and lysed as described previously (38). Cell lysates (100–200 μg) were then immunoprecipitated with GFP antibody (Thermo Fisher Scientific # G10362). Ubiquitination was detected as described previously (38).

#### Determination of GFP-SET8 proteins half-life

HeLa cells were transfected with 500 ng of plasmids encoding for GFP, GFP-SET8FL or GFP-SET8KA. After 24 hrs, 50 mg/ml of cycloheximide (Sigma-Aldrich # C4859) was added and cells were collected at different time points until 22 hrs. Cells were extracted using RIPA buffer. GFP protein levels were detected by Western blot. After densitometry analyses, the half-life of GFP-SET8FL and GFP-SET8KA were calculated using least square method from PRISM 9 where the known equation of Half-life calculation was fitted for GFP-SET8FL and GFP-SET8KA respectively.

#### PARP1 and SET8 cloning and mutagenesis

Sequences coding for PARP1 full-length (cDNA obtained from Origen # RC20708), SET8 full-length (39) were cloned into pEGFP-C2 or pGEX5X.1 vector and transformed into NEB 10-beta competent E.Coli (New England Biolabs # C3019H). Mutagenesis to generate PARP1, SET8 domains as well as point mutations were performed using Q5 site-directed mutagenesis kit (New England Biolabs # E0554S). Primers used for cloning and mutagenesis are available upon request.

#### PARP1 and SET8 interaction studies

For *in vitro* interaction studies, PARP1 and SET8 GST-fusion and mutant proteins were induced in Escherichia coli ER2566 cells (New England Biolabs # C2566H) using 0.4mM IPTG overnight at 16°C. Purification of recombinant fusion proteins from the bacterial lysate were performed as previously described (40). For GST pull-down assay, GST or the GST fusion proteins was bound to glutathione-Sepharose beads. The assay was performed by pre-incubating the GST or GST fusion protein beads with 100 µg/ml bovine serum albumin (BSA) and the protein to be studied in a binding buffer (1X PBS with 0.1% Triton X-100) with end over end mixing at 4°C for 1 h. Beads were washed 3 times for 5 min with a binding buffer containing 1% Triton X-100. The beads were mixed with 1× SDS–PAGE sample loading buffer (New England Biolabs # B7703S) and incubated at 98°C for 5 min. The protein mixtures were separated on a 10% polyacrylamide Tricine gel (Thermo Fisher Scientific # EC6675BOX). Recombinant PARP1 was purchased from Active Motif (# 81037), SET8 was purified from New England Biolabs (# M0428).

Co-immunoprecipitation of endogenous PARP1 and SET8 were performed with 200 µg of total extract from crosslinked HCT116 or HeLa cells with 1% formaldehyde for 10 min using anti-PARP1 antibody (Cell Signaling Technology # 9532), anti-SET8 antibody (Santa Cruz Biotechnology # sc-515433) or 5 µg of rabbit IgG as a control antibody (Santa Cruz Biotechnology # sc-2027). The immunoprecipitations (IP) were performed overnight with end-over-end mixing at 4°C in TD buffer (50 mM HEPES, pH 7.5, 250 mM NaCl and 1% Triton X-100). The antibodies were captured using 50 µl of protein G magnetic beads (S1430, New England Biolabs) incubated for 60 min with end over end mixing at 4°C. After 3 washes with TD buffer, IP reactions were blotted using anti-PARP1 antibody (Millipore-Sigma # HPA045168) and rabbit anti-SET8 antibody (Cell Signaling Technology # 2996S). For immunoprecipitation of SET8 from synchronized HeLa cells, 200 µg of chromatin cell lysate was incubated with 5 µg of anti-SET8 (sc-515433, Santa Cruz). IP reactions were performed and blotted with anti-PARP1 and anti-SET8 antibodies as described above.

#### Immunofluorescence studies

For the detection of PARP1 and SET8 colocalization, COS-7 cells were grown on coverslips and co-transfected with FLAG-PARP1 and GFP-SET8 plasmids as described above. After 48 hrs, cells were crosslinked with 4% paraformaldehyde (Electron Microscopy Sciences # 15710) for 10 min at RT and quenched with 0.125M Glycine for 5 min at RT. After 20 min permeabilization with 100 % methanol for 20 min at -20°C, cells were incubated with PBS including 0.5 % Tween20 and 5% BSA (Millipore-Sigma # A-7906) for 1 hr at RT. Epitope tagged PARP1 was detected by mouse anti-FLAG antibody (# F3165, Millipore-Sigma) and visualized with an anti-mouse IgG coupled with Alexa Fluor 594 dye (Thermo Fisher Scientific # A-11005). GFP-SET8 and FLAG-PARP1 were detected using 458, 488, 514 nm multiline Argon laser and 561 nm DPSS laser, respectively. Endogenous PCNA was visualized with anti-PCNA Alexa Fluor 647 conjugate (Cell Signaling Technology #82968S) using 633 nm HeNe laser. Slides were mounted using Prolong Gold Antifade Mountant with DAPI (Thermo Fisher Scientific # P36931). Images were captured using a confocal microscope (LSM 880, Zeiss).

DNA replication sites were detected using EDU Staining Proliferation Kit (iFluor 647, ABCAM # ab222421) on GFP-SET8 full-length transfected COS-7 cells according to manufacturer’s recommendations. Before EDU labeling, GFP-SET8 transfected COS-7 cells were pretreated with 20 µM of PARG inhibitor (Millipore-Sigma # SML1781) for 1 hr. After EDU labeling, cells were incubated overnight at 4C with anti-pan-ADPribose reagent at 1/400 dilution (Millipore-Sigma # MABE1016). ADPribose was detected using anti-rabbit IgG coupled with Alexa Fluor 594 dye (Thermo Fisher Scientific # A32740). EDU was visualized using 633 nm HeNe laser. Slides were mounted and images were captured as described above.

#### SET8 gel shift DNA-nucleosome binding assays

For SET8 DNA gel shift assay, Cy3-end-labeled EcoRI hairpin oligonucleotide (0.5 mM) (GG<u>GAATTC</u>CCAAAGG<u>GAATTC</u>CC, EcoRI sites underlined) and different concentration of recombinant His-SET8 protein (0 to 1.4 mM, New England Biolabs) were incubated for 10 min on ice in 1× GRB binding buffer [20 mM HEPES pH 7.5, 10% glycerol, 25 mM KCl, 0.1 mM EDTA, 1 mM dithiothreitol (DTT), 2 mM MgCl2, 0.2% NP40] in a 20 µl reaction volume. Protein–DNA complexes were resolved on a 6% TBE DNA retardation gel (Thermo Fisher Scientific # EC6365BOX) at 4°C in 0.5× TBE (45 mM Tris–HCl, 45 mM H3BO3, 10 mM EDTA pH 8.3) at 140 V. Complex were visualized using Typhoon scanner. Kd was calculated using GraphPad Prism 8.0.

For GST-SET8 full-length as well as mutant DNA or nucleosome binding, 5 µg of GST-beads were incubated for 15 min on ice in GRB buffer in a 25 ml reaction volume with 1 µg of 100 bp DNA ladder (New England Biolabs # N3231S) or mononucleosome (Active motif # 81070). After 1, 000 rpm spin for 5 min at 4°C, supernatant representing the unbound DNA was mixed with 6x purple dye (New England Biolabs # B7024S) and loaded on a 6% TBE DNA retardation gel (Thermo Fisher Scientific # EC6365BOX) at 4°C in 0.5× TBE at 140 V. DNA was visualized under UV Transilluminator. In some cases, GST or GST-SET8 full-length fusion proteins were ADP-ribosylated using PARP1 recombinant enzyme (Active Motif # 81037) during 15 min at RT. The GST beads were then washed twice with PBS+1% Triton X-100 and 1M NaCl to remove PARP1 bound to the beads. After 2 other washes with 1X PBS containing 0.1% Triton X-100, beads were resuspended in GRB buffer and incubated with 1 µg of 100 bp DNA ladder (New England Biolabs # N3231S) or mononucleosome (Active motif # 81070) for 10 min on ice. After 1, 000 rpm spin for 5 min at 4°C, supernatant representing the unbound DNA was loaded on a 6% TBE DNA retardation gel and detected under UV Transilluminator.

#### ADP-ribosylation and SET8 methyltransferase assays

For ADP-ribosylation assay, PARP1 recombinant enzyme (from 20 to 50 nM per reaction) was incubated with different substrates including either His-SET8 full-length recombinant protein, SET8 peptides or GST, GST-SET8 full-length or mutants and activated using EcoRI hairpin oligo as described above. The reaction was performed in 1x buffer (50 mM Tris.Cl, pH.8, 4 mM MgCl2 and 250 µM DTT) in the presence or absence of cofactor ß-Nicotinamide adenine dinucleotide (0.5 mM NAD+ per reaction, New England Biolabs # B9007S) at RT during 15 min in 20 ml total reaction volume. The reaction was then loaded on 10% Tricine gels and subjected to Western blotting. ADP-ribosylation was detected by mono/poly-ADP-ribose (Cell Signaling Technology # 83732S) or polyADP-ribose (monoclonal 10H, Santa Cruz Biotechnology # sc-56198) antibodies.

For mass spectrometry analyses, 500 nM of PARP1 recombinant enzyme was incubated with 160 µM SET8 peptide (158-170 aa: KKPIKGKQAPRKK and 86-98 aa: KPLAGIYRKREEK) from 1 hr to overnight at RT.

Histone methyltransferase assays were carried out as described previously [41]. SET8 recombinant enzyme (New England Biolabs # M0428S) was incubated with recombinant active PARP1 (Trevigen # 4668-100-01) with or without EcoRI hairpin oligo or NAD+ in histone methyltransferase buffer and 6 µM of radiolabeled [3H] AdoMet (Perkin Elmer Life Science # NET155V001MC). Recombinant human histone H4 (New England Biolabs # M2504S) was used as a substrate. Filter disc method was used to process the samples and the [^3^H]CH3 incorporated into the H4 protein was determined using a liquid scintillation counter.

### LC-MS analysis of poly ADP-ribosylated peptides

Peptide solutions were analyzed with ProxeonII nLC – LTQ Orbitrap XL by direct injection on Reprosil-Pur C18-AQ 3µm 25cm column and eluted at 300nL/min. Full scan MS was acquired FT Resolution 60k in orbitrap MS. The most abundant three ions were selected for data-dependent CID MS/MS fragmentation with normalized collision energy 26. Using the same nanoLC conditions, HCD MS/MS fragmentation was acquired using quadrupole-orbitrap with NCE 27. Isotope peak intensity areas were determined with SIEVE 2.2.58. (Thermo Fisher Scientific) following chromatographic alignment. Chromatographic extracted ions were plotted with the manufacturer’s software XCalibur. Potential location of ADPr unit was screened using a match score modeling of fragment ions in CID MS/MS spectra. The matching score was calculated with Peptide Sequence Fragmentation Modeler, Molecular Weight calculator (https://omics.pnl.gov/software/molecular-weight-calculator). The algorithm is based on Sequest Sp preliminary score but differs in treatment of immonium ions (42). The HCD spectra were mapped manually to the sequence of ADP-ribosylated peptides.

#### Chromatin immunoprecipitation sequencing (ChIP-seq)

HeLa cells overexpressing PARP1 or knockdown PARP1 with esiRNA as mentioned above were subjected to ChIP-seq. Briefly, chromatin was extracted as described above and crosslinked with 1% formaldehyde for 10 min and quenched with 0.125 M glycine. After sonication, H4K20me1 IP was performed overnight in TD buffer as described above using 5 µg of H4K20me1 antibody (Thermo Fisher Scientific # MA5-18067). After antibody capture with protein G magnetic beads and 3 washes with TD buffer, beads were incubated at 65°C overnight in buffer including 50 mM Tris-HCl, pH 7.5, 200 mM NaCl, 1% SDS and 1.6 U/µl of proteinase K (New England Biolabs # P8107S) to reverse crosslinks. DNA from supernatants was extracted using phenol/chloroform procedure. Between 1 and 10 ng of DNA were used to generate DNA libraries for subsequent sequencing analyses. DNA libraries were made using NEBNext Ultra II DNA Library Prep Kit for Illumina (New England Biolabs # E7645S) according to the manufacturer’s recommendations.

#### RNA-sequencing (RNA-seq)

RNA from HeLa cells overexpressing PARP1 or knockdown PARP1 with esiRNA as mentioned above was extracted using Quick-RNA Miniprep Kit (Zymo Research # R1054). 1 µg of RNA was used to isolate Poly(A) mRNA using NEBNext Poly(A) mRNA Magnetic Isolation Module (New England Biolabs # E7490S). NEBNext Ultra II directional RNA Library Prep Kit for Illumina (New England Biolabs # E7760S) was used according to the manufacturer’s recommendations to generate cDNA and DNA libraries for subsequent DNA sequencing analyses.

### Data analysis

#### ChiP-seq data processing

The raw fastq sequences were trimmed using Trim Galore (http://www.bioinformatics.babraham.ac.uk/projects/trim_galore/) to remove the adapters and low-quality sequences. Trimmed reads were mapped to the human reference assembly hg38 using Bowtie2 (45). Aligned reads in bam format were filtered for duplicates and low-quality alignments using Picard (http://broadinstitute.github.io/picard/) and samtools (46). The aligned bam files of technical replicates were merged using Sambamba (47). H4K20me1 enriched regions were identified by calling broad peaks (target over input) using MACS2 (48) where the parameter broad-cutoff was set to 0.025 for more robustness . Signal tracks were generated using deeptools bamCoverage (49) with the parameters, -normalizeUsing RPKM -of bigwig -e. Spearman correalation analysis was performed using deeptools plotCorrelation (49) function. H4K20me1 peaks were annotated using HOMER (50) annotatePeaks.pl. Repeats elements were annotated using HOMER (50). H4K20me1 profile in the gene regions for PARP1 from Hela cells for bothe the conditions was computed with the deeptools computeMatrix and plotProfile (49) functions. Peak length was calculated for all the conditions to estimate the gain and loss of H4K20me1 after PARP1 knock down. Transcription factor binding motifs enrichment near H4K20me1 peaks was searched using the HOMER tool findMotifsGenome.pl (50) with default parameters. The enrichment scores (-10log(P-value) of individual TF binding motifs were calculated for PARP1 overexpressed and knockdown HeLa cells and their respective controls. Then the enrichment scores were summarized into a data matrix in R (51) and a heatmap was then created using heatmap.2 function to represent condition-specific enrichment of TF binding motifs near the H4K20me1 peaks. Genomic regions were visualized using Integrative genomic viewer (IGV) (53).

#### ChromHMM

ChromHMM (54) was used to create epigenomic segmentations for PARP1 overexpressed and knockdown HeLa cells and their respective controls using bam files for ChIP-seq of H4K20me1, H4K20me3 from siPARP1 and siGFP. A 15-state model was trained using 200 bp bins (LearnModel -b 200) and genome version hg38.

#### RNA-seq

The raw fastq files were trimmed and quality check was performed using Trim Galore and FASTQC (https://www.bioinformatics.babraham.ac.uk/projects/fastqc/) respectively. All the samples were in good quality. Trimmed reads were mapped to the human reference assembly hg38 using STAR (55). Transcript abundance was estimated from these high-quality mapped reads using htseq count module (56). DESeq2 (57) was used to normalize the count matrix and identification of differentially expressed genes. Genes that were showing logFC≥0.5 at FDR < 0.05 and logFC≤0.5 at FDR < 0.05 were considered up-regulated genes and down-regulated respectively. In this experiment, two sets of RNA seq data were utilized from siGFP and siPARP1 conditions respectively where siGFP was considered as control data. Initially, the positively expressed genes from siGFP and siPARP1 conditions were compared with genes of peak annotated files from intragenic regions of H4K20me1 and H4K20me3 Chip seq data. Subsequently, the transcript abundance, estimated from mapped reads of Chip seq were compared with respected RNAseq expression. More elaborately, the up regulated genes, identified from Chip-Seq data from H4K20me1 and, H4K20me3 (where siGFP_H4K20me1 and siGFP_H4K20me3 were used as control) were compared with the corresponding expression values from RNA seq Data and vice versa. To compare, the data distribution, 2D density plotting from ggplot2 from R had been considered where kernel distribution estimator (KDE) was used to plot the random variables in terms of gene expression from the two different data sources under a polygonal space. These helped to understand the similarity between expression from Chip-Seq and RNA-Seq.

#### Model Building

In this experiment, the monomeric sequence of SET8 was modeled using I-TASSER (77)and considered as Wild Type SETD8 (WT SET8). In this case, the main focus was on lysine at residue 86, 158, 159, 162, 164, 168, 169, 170, 174. For the mutated structure, all these positions had been replaced with alanine. I-TASSER produced top five energy minimized models in each case using *ab initio* method. The top models from WT and mutated SET8 were selected for further studies.

#### Studying the folding pattern and Predicting the Structural Disorder

To observe the nature of the DNA binding region, we have utilized to web based tools namely, PONDR-VLXT (78), and FoldIndex (79). PONDR-VLXT is a intrinsically disorder region prediction tool where predctive disorder score ≥ 0.5 of amino acid residues have been considered as intrinsically disordered residues. Rate of disorderedness can be defined as value of predictive disordered score. Similarly, FoldIndex can calculate the index values for each amino acids ranging between 1 to -1 where any residues having index value lower than zero is considered as unfolded residues. Rate of unfoldedness is highly dependent on the index value.

#### Structure Network Analysis of Wild Type and Mutant SET8

Analysis and prediction of dynamics associated with complex protein systems can be explained and represented using network architecture. In general, a complex system is composed of elements interacting with one another bound together by links like interactions (80). Weighted edges characterize the strength of interaction. Overlapping modules can, in turn, be dissected from the network (i.e., communities, groups), formed the modules.

Protein network structure is measured using topology of complex 3D architecture commonly known as Gaussian Network Model (GNM). In this approach, a weighted graph G was constructed that represented a 3D structure, *(V, E) ∈ G, where V (V = V1, V2 … Vn)* represented residues as nodes and *E (E = E1, E2, … En)* weighted edges represented pairwise interaction. The internal motions and intrinsic dynamics of proteins dictate the global protein structure and, hence, the function and activity. Normal Mode analysis (NMA) was utilized for predicting the functional motions in SETD8. Elastic Network Model using C-alpha force field was designed through NMA. Subsequently, the cross-correlation study was performed to identify protein segments with similar oscillation, and a matrix was generated using correlation coefficient score. The correlation-based matrix was further used as a full residue adjacency matrix. These networks were split into a highly correlated coarse-grained community cluster network using the Girvan– Newman clustering method (which was highky dependent on edge betweenness) (81). Betweenness centrality characterized the regions of a protein that show modifications in coupled oscillations derived from mutant as well as the WT protein. Residues having a significant contribution to the intrinsic dynamics of the protein show high centrality value.

## Results

### PARP1 ribosylates SET8

In a proteomic analysis of SET8 pull-down in HEK293T cells, we discovered that PARP1 is a strong binder (Table S1). To reconfirm this observation, we performed reciprocal immunoprecipitation either with anti-SET8 or anti-PARP1 antibody, performed western blots, and probed with respective antibodies. Indeed, PARP1 pulled down SET8 and vice-versa, compared to IgG control (Fig. 1A). This led us to investigate if this interaction is cell cycle-dependent, since SET8 expression is regulated by cell cycle (55). For this experiment, we transfected COS-7 cells with FLAG-PARP1 and GFP-SET8 and studied their association using confocal microscopy. The same cells were also probed with an anti-PCNA antibody to examine the chromatin replication foci. Indeed, PCNA remained punctate in the nucleus, so as GFP-SET8, and both were colocalized. However, as expected PARP1 remained throughout the nucleus and appeared as prominent punctate foci with both PCNA and SET8 as observed by bright yellow spots (Fig. 1B). These observations suggest that PARP1 has multiple targets throughout the nucleus, but also has strong colocalization with SET8 at the chromatin replication foci. To narrow down the exact binding motifs between PARP1 and SET8, we performed a reciprocal GST-pulldown assay. Immobilized GST fusion of overlapping SET8 fragments covering the entire protein was challenged with full-length PARP1. After stringent washes, the bound proteins were denatured in SDS gel loading buffer and separated on SDS-PAGE followed by blotting onto a membrane for western blot with anti-PARP1 antibody to determine the binding domain. Two fragments covering both N-terminal 1-98 amino acids and disorder domain 80-230 amino acids residues showed binding. Although, a stronger binding was observed with the disorder domain (DD) (Fig. 1C, left panel). A similar reciprocal GST pulldown experiment using immobilized GST-PARP1 fusion fragments covering the entire protein was challenged with full-length SET8. The bound proteins were western blotted and probed for SET8 to reveal GST-PARP1 DNA binding domain (GST-PARP1DB) as the strongest binder (Fig. 1C, right panel). Based on the results of both the colocalization and GST pulldown experiments, we concluded that both SET8 and PARP1 are indeed binding partners.

**Figure 1:**
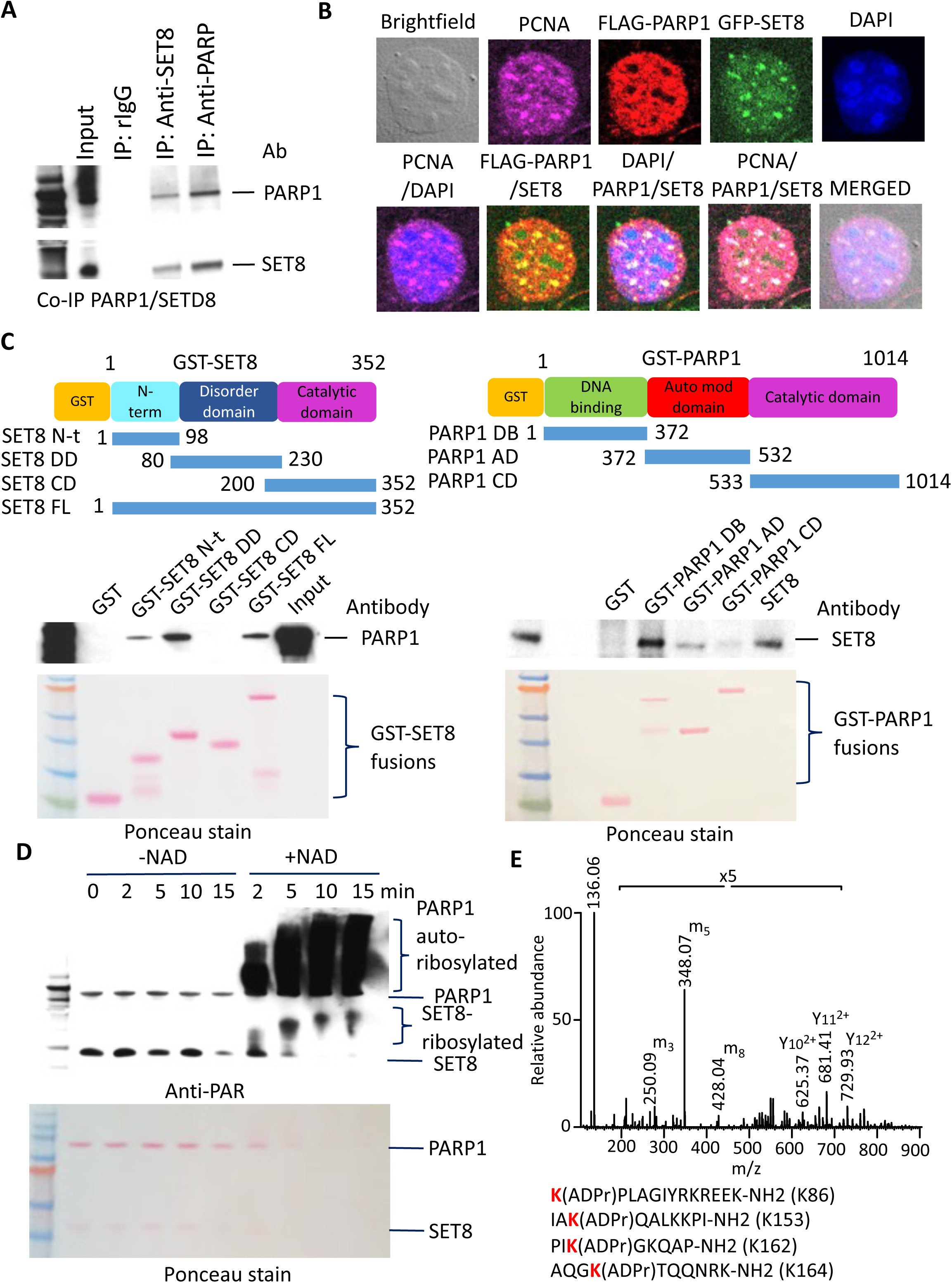
Colocalization, binding and poly ADP-ribosylation of SET8 by PARP1. (A) Co-immunoprecipitation of endogenous SET8 and PARP1 in HCT116 cells. Rabbit IgG (left lane) was used as a negative control. (B) Colocalization between FLAG-PARP1 (red), GFP-SET8 (green) and endogenous PCNA (pink) in COS-7 cells. DAPI (blue) represents the nuclear DNA content. (C) Mapping of domain interactions using GST-pulldown assays between SET8 domains (top, left) and full-length recombinant PARP1 protein and GST-pulldown assays between PARP1 domains and full-length recombinant SET8 protein (top, right). The PARP1 or SET8 binding were detected by western blotting using PARP1 antibody (middle, left) or SET8 antibody (middle, right) respectively. Ponceau stain gels (bottom) represent the amount of GST beads constructs used for the GST-pulldown assays. (D) In vitro detection of full-length recombinant SET8 ADP-ribosylation by full-length recombinant PARP1 by western blotting using anti-ADP ribose antibody (top). Ponceau stain gels (bottom) represent the amount of PARP1 and SET8 recombinant enzyme used for ADP-ribosylation assay (bottom). (E) Detection of SET8 lysines ADP-ribosylation using mass spectrometry analysis of SET8 ADP-ribosylated peptides by full-length recombinant PARP1 protein in vitro.

We next investigated the functional consequence of this interaction. Both purified SET8 and PARP1 were incubated together in the absence or presence of cofactor NAD+ for various lengths of time, spanning between 2-15 min. The reactions were stopped, and the proteins were separated on SDS PAGE, western blotted and probed with anti-PAR (anti-poly ADP ribose) antibody. If the proteins are ribosylated, an anti-PAR antibody will show higher molecular weight shift for the proteins. We observed auto-poly-ADP-ribosylation of PARP1 as expected, only in the presence of NAD+ (Fig. 1D, upper panel; Fig. S1A). We also observed a strong reaction time-dependent SET8 poly ADP-ribosylation. Indeed, all SET8 molecules were poly ADP-ribosylated by PARP1 within 10 min of reaction as observed by high molecular weight migrating smear (Fig. 1D). We further investigated the poly ADP-ribosylated regions of SET8 by performing *in vitro* poly ADP-ribosylation assay of overlapping peptides followed by western blotting and probing the blot with anti-PAR antibody. We found that GST-SET8 DD is indeed the substrate of PARP1 (Fig. S1B). We narrowed down the amino acid residues 81-98 and 157-180 as the putative acceptor of ADP-ribose for poly ADP-ribosylation (Fig. S1C, D).

### LC-MS analysis of PARP1 activity on SET8 peptides

To determine the nature of poly-ADP-ribosylation and the acceptor amino acid, we made synthetic peptides covering 86-98 and 158- 170 amino acids and performed *in vitro* poly ADP-ribosylation, and analyzed the reaction product using LC-MS. First, we monitored the PARP1 activity with SET8 peptide KPLAGIYRKREEK-NH2 (86-98 aa). Peptides with an amidated C-terminus require specific conditions for detection to support MS/MS sequencing identification (56). We detected and quantified signals for the three main isotopes starting with 0 min (negative control) and incubation with PARP1 for 1hr, 4hrs and overnight (Fig. S2A). We observed a decrease in the signal of the KPLAGIYRKREEK-NH2 peptide after the initial 1hr incubation. Concomitantly, we assessed the peak areas for two charged states (+3, +4) of the (ADPr) KPLAGIYRKREEK-NH2 in order to detect the activity of PARP1 (Fig. S2B, C). Three main isotopes for each charge state showed the signal of (ADPr)KPLAGIYRKREEK-NH2 present at 1hr followed by a decrease in intensity at 4hrs and overnight reactions. This suggested the formation of additional species beyond the initial addition of one ADP-ribosylation unit. Next, we investigated the peptide signals at two faster PARP1 reaction times, 5 min and 15 min.

Chromatographic ion signals for (1ADPr)KPLAGIYRKREEK-NH2 were detected for both charge states +3 and +4 at retention time 8.35 min (Fig. S2D). The minimal chromatographic signal was detected for (2ADPr)KPLAGIYRKREEK-NH2 with an increased elution time of 10.33 min. At 15 min reaction time, the signal for di-ADP-ribosylated peptide near 10.3 min increased with a wider base (Fig. S2E). A second peak with intermediate hydrophobicity relative to the mono ADP-ribosylated peptide showed the most intense di-ADP-ribosylation signal at RT 9.77 min (Fig. S2E). Therefore, we detected the formation of possible diastereomers of increased hydrophobicity upon addition of 1ADPr unit to mono ADP-ribosylated peptides during a 10 min reaction time interval.

Characterization of MS/MS HCD fragmentation pattern of (ADPr) KPLAGIYRKREEK-NH2 peptide is presented in Fig. S3. For charge state +3, we observed the formation of fragment ions m3, m5, and m8 that are independent of the peptide sequence and indicative of the ADP-ribose group (Fig. S3A, top panel). The nomenclature for ADP ribosylation fragment ions is according to Hengel et al. (57). Similarly, but with lower intensity, charge state +4 showed confirmed the presence of these fragment ions m3, m5 and m8 (Fig. S3B, bottom panel). Intact peptide backbone ions containing partial ADP ribose group and peptide y sequencing ions can be visualized at m/z 500 – 900 (Fig. S3A, B, bottom panels; Fig. S3B is the same spectrum from Fig 1E). In addition to HCD spectra, CID spectra can be probed for ADP-ribosylation information of amidated C-terminus peptides. We screened seven amino acids for the position of the ADPr unit (Fig. S4). For both charges +3 and +4, the associated MS/MS spectra mapped with a higher score the N-terminal lysine of KPLAGIYRKREEK-NH2.

Analysis of SET8 peptide KKPIKGKQAPRKK-NH2 (158-170 aa) showed that activity of PARP1 resulted in maximum signal for 1ADPr unit addition at charge +4 for overnight reaction (Fig. S5A). Detection of lower intensity peak areas for all three main isotopes of charge +4 together with moderate signal decrease of unmodified KKPIKGKQAPRKK-NH2 (Fig. S5B) suggested a slower PARP1 kinetic towards this SET8 peptide. For comparison, CID MS/MS spectrum of this charge state with unit resolution linear ion trap showed the presence of ADP-ribose signature ions (Fig. S6). Taken together, we found multiple Lys residues were poly ADP-ribosylated, including but not limited to K86, K153, K162, and K164 (Fig. 1E).

### Mutation of lys residues in SET8 alters the Disorder Domain

SET8 has a prominent disordered domain with PONDR-VLXT score above 0.5 and fold index lower than 0 (Fig. S7A, B). The disorder domain binds strongly to PARP1 (Fig. 1C). It is composed of charged amino acids residues such as lysines that are also poly ADP ribosylation accepter (Fig. 1E). To determine the role of the amino acids and poly ADP-ribosylation in PARP1 binding, GST-SET8 fusion was mutated at K86A, K158/159A, K162/164A, R168A, K169/170A, K174A (GST-SET8 M). The disorder domain of the SET8m displayed ordered structure with PONDR score below 0.5 suggesting the mutant lysine residues facilitating this event (Fig. S7A, C vs. B, D). Furthermore, a brief network algorithm guided structure space analysis was performed to determine the effect of the structural modification due to amino acids substitution. Structures of the monomeric SET8 and SET8m were modeled using I-TASSER prediction tool (Fig. S8). The modelled structure displayed a prominant binding cleft in the wild type compared to the mutant (Fig. S8A, B). Overall, the distance among the original Lys/Arg or substituted Ala residues at position 86, 158, 159, 162, 164, 168, 169, 170 and 174 (within the domain of wt SET8) was lower for SET8m (Fig. S8C-F; Table S2). Subsequently, normal mode based GNM utilized the residual oscillation scores for studying the structural modification. These scores were also used to determine the weight between two amino acids residues (given as nodes) which can further be considered as co-oscillation possibility. The edge betweenness based clustering model considered these weights for shortest path calculation. Therefore, the centrality score assigned for each residue could be used as structural dependency on those residues. In Fig. S9, significant modifications of betweenness centrality score were observed (Fig. S9A, B right panels). As per the distributions of the scores throughout the structures, localize residue specific dependencies were higher in SET8m comparing to wild type which could be elaborated more through module detection. The number of modules were 10 and 12 for wt and SET8m respectively (Fig. S10). The incremental number of modules can be considered as the indication of minor orderedness where the member of each clusters helped to analyze it further. Cluster 5 from wild type enzyme had all the lysine enriched residues from disordered regions. However, these residues are distributed in three different clusters, 5, 8, and 9 in SET8m. Thus, it is plusible that SET8m has higher propensity of localized orderedness than wild type enzyme that may affect protein-protein interaction and other macromolecule binding. To test our hypothesis, we performed GST-pulldown assay by incubating purified PARP1 with the full-length GST fusion SET8 (GST-SET8 FL) or the mutant (GST-SET8 M). After several washes the bound proteins were separated on SDS-PAGE, westen blotted and probed with anti-PARP1 antibody. We observed significant (∼70%) loss of PARP1 binding, confirming SET8 DD lysine residues are indeed essential for this intraction (Fig. S11).

### PARP1 Poly ADP-ribosylates SET8 impacting DNA and Nucleosome Binding

SET8 is well known for its interaction with nuclear proteins, PCNA, a processivity factor involved in DNA replication and required for S-phase progression (55, 58). A crystallography study demonstrated that SET8 employs its i-SET and c-SET domains to engage nucleosomal DNA 1 to 1.5 turns from the nucleosomal dyad (59). To evaluate the DNA binding activity of SET8, we incubated a 100 bp DNA ladder with various GST-fusion fragments of SET8, GST-SET8 FL and GST-SET8 M. After the incubation time, we resolved fusion proteins -DNA or nucleosome complexes on the gel. If a defined size of DNA is bound with protein, that DNA band will be shifted and will not be represented on the gel compared to the control ladder. As predicted by crystallography, GST-SET8 FL protein bound predominately to double-stranded DNA ranging between 100-300 bp in the gel-shift assay. This binding was partially dependent on the 157-352 amino acids of SET8, and a significant loss of binding was observed in a deletion mutant comprising 175-352 amino acids, suggesting 157-175 amino acids play a functional role in DNA binding (Fig. 2A). We also performed gel shift assay of purified SET8 protein with a fluorescent-labeled double stranded DNA oligo and measured the Kd values of 1.6+/-0.6 µM (Fig. 2B). After observing the robust DNA binding activity of SET8, we also validated its binding activity with recombinant mono-nucleosomes using similar GST-SET8 fusions that were used for DNA binding activity studies shown in figure 2A. Once again, 157-175 amino acids played a functional role in nucleosome binding (Fig. 2C). Taken together, data from previous crystallography studies and our gel-shift assays, we conclusively demonstrated that SET8 has both double-stranded DNA and nucleosome binding activity.

**Figure 2:**
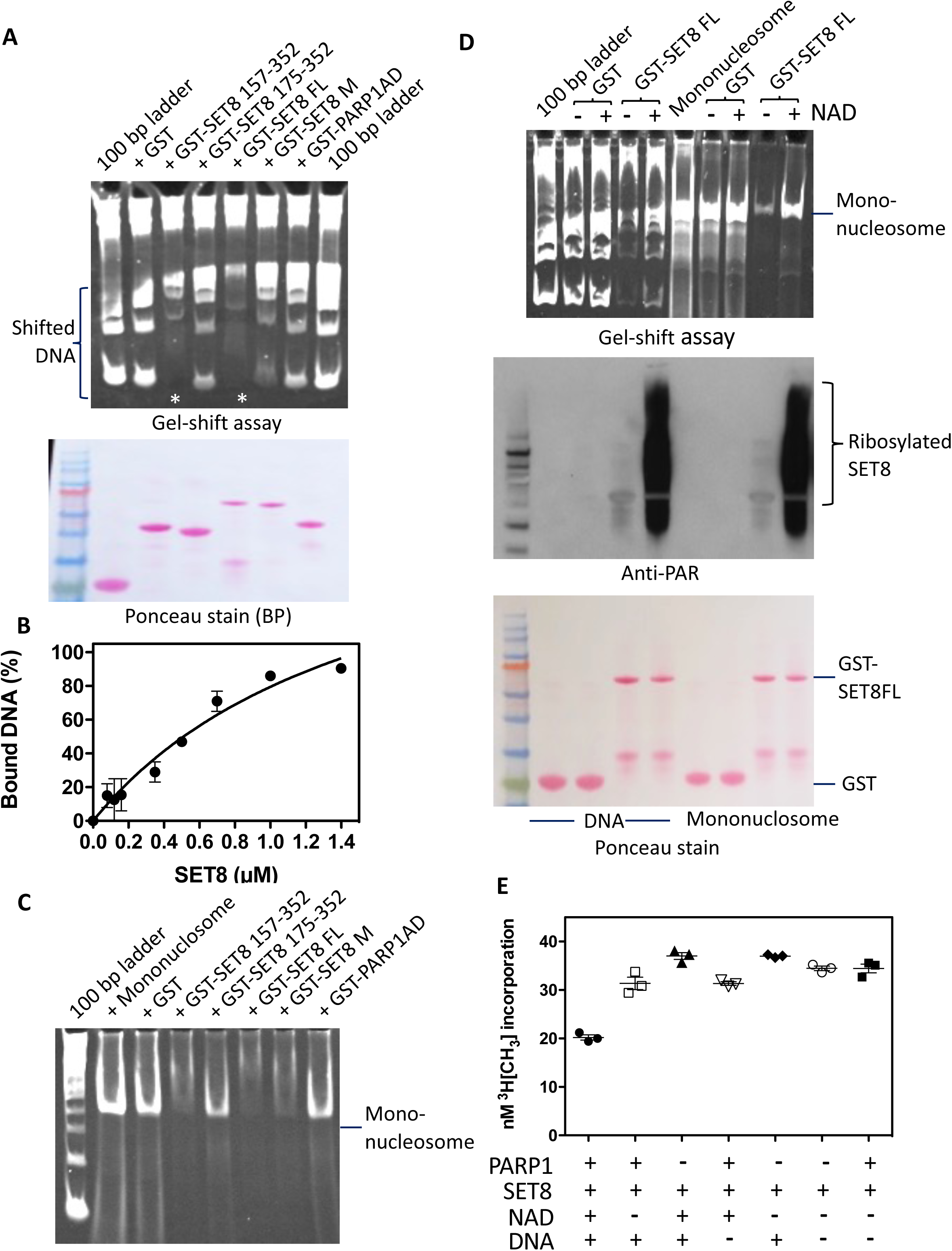
Poly ADP-ribosylation impairs DNA, nucleosome binding and catalytic activity of SET8 protein. (A) Detection of unbound 100 bp DNA ladder by TBE ethidium bromide-stained gel in supernatants (left lane) on GST-SET8 domains or mutant (M) using GST-pulldown assays (top, left side). Asterisks are representing shift of DNA on GST-SET8 157-352 amino acid protein or GST-SET8 full-length protein (FL) beads. Ponceau stain represents the amount of GST beads constructs used for the GST-pulldown (bottom, left side). (B) Different concentrations of recombinant full-length SET8 protein binding to DNA using EMSA to determine the equilibrium dissociation constant (Kd). (C) Detection of unbound mononucleosome by TBE ethidium bromide-stained gel in supernatants on GST beads versus GST-SET8 domains or mutant (M) beads using GST-pulldown assays. (D) Detection of unbound DNA (left side) or unbound mononucleosome (right side) by TBE ethidium bromide-stained gel in supernatants (top) on GST beads versus GST-SET8 FL beads using GST-pulldown assays. GST or GST-SET8 FL beads were poly ADP-ribosylated or not (with or without NAD) using full-length recombinant PARP1 as demonstrated by western blot using anti- ADP ribose antibody (middle). Ponceau stain gel (bottom) represents the amount of GST bead constructs used for the GST-pulldown and the ADP-ribosylation western blot analysis. (E) SET8 histone methyltransferase assay on full-length recombinant histone H4 using full-length recombinant SET8 in presence or absence of activated full-length recombinant PARP1.

Since SET8 is a substrate for PARP1, we investigated the role of DNA in SET8-PARP1 binary complex formation, and whether poly ADP-ribosylation can affect either DNA or nucleosome binding activity of SET8. DNA is a catalytic activator of PARP1, we incubated GST-SET8, DNA and PARP1 in the presence and absence of DNase I and performed GST-pull down assay followed by western blotting and probing the bound proteins with respective antibodies. Indeed, both GST-SET8 and PARP1 formed binary complexes in the presence of double-stranded DNA. However, in the presence of DNase I, SET8-PARP1 binding was reduced confirming DNA as a facilitator of binary complex formation (Fig. S12A, B).

### PARP1 poly ADP-ribosylates SET8 impacting catalytic activity

Since SET8m displayed ∼70% loss of PARP1 binding compared to the wild type enzyme and the mutations were in poly ADP ribosylation acceptor amino acids, we hypothesized that ADP ribosylation residues would impact its substrate binding, thus impacting catalytic activity. To examine our hypothesis, we incubated SET8 with either a 100 bp ladder or with a recombinant mononucleosome in the presence of PARP1 and NAD+ for poly ADP-ribosylation, and the control remained without NAD+ cofactor. Indeed, poly ADP-ribosylated SET8 lost the DNA as well as nucleosome binding activity as observed by prominent DNA or nucleosome bands (Fig. 2D, top panel). To confirm that this observation is due to poly ADP-ribosylation, a portion of the same reaction was western blotted and probed with anti-PAR antibody. Indeed, the SET8 poly ADP- ribosylation was evident in those lanes that had poor binding of DNA or nucleosome with GST-SET8 (Fig. 2D middle panel).

Next we evaluated if SET8 methyltransferase activity on histone H4 is modulated by PARP1 mediated poly ADP-ribosylation of the enzyme. We performed *in vitro* histone methyltransferase assays mimicking either poly ADP-ribosylated SET8 or its unmodified form. To initiate poly ADP-ribosylation of SET8, we pre-incubated SET8, PARP1, DNA and NAD+ for 15 min at room temperature, followed by the addition of methyl donor, tritiated AdoMet, and purified recombinant histone H4 to initiate H4K20 monomethylation. After the reaction, we performed the filter binding assay for tritium incorporation on substrate histone H4. Indeed, poly ADP-ribosylation SET8 lost ∼40-50% activity compared to the control that lacked any one component for successful poly ADP-ribosylation (Fig. 2E). These results suggest that PARP1 not only inhibits SET8 binding to DNA and nucleosome, but it can also impact histone H4K20 monomethylation.

### Poly ADP-ribosylation promotes SET8 Degradation

Poly ADP-ribosylation was recently identified as a signal for triggering protein degradation through the ubiquitin-proteasome system. Indeed, poly ADP-ribosylation-mediated degradation of ARTD1 (ADP-ribosyltransferase diphtheria toxin-like 1) is documented (60). In another example, PARP1 binds and poly ADP-ribosylates bromodomain-containing protein 7 (BRD7), which enhances its ubiquitination and degradation through the PAR-binding E3 ubiquitin ligase RNF146 (61). This led us to investigate if PARP1 has any modulating effect on SET8 levels in the cell. First, we transfected cells with either FLAG or FLAG-PARP1 construct and measured the transfection efficiency by western blotting and probing with anti-FLAG antibody, followed by probing the blot with anti H4K20me1, 2 and 3 along with SET8. Indeed, PARP1 overexpression had reduced SET8 and its reaction products, H4K20me1 to almost half of the control (Fig. S13A, B). We then systematically transfected HA-ubiquitin with GFP-SET8 or FLAG-PARP1 alone or co-transfected all three constructs into the mammalian cells and treated the cells with proteasome inhibitor MG132. We performed western blots of cell extracts to evaluate transfection efficiency and expression of constructs. Probing the blot with anti-GFP antibody demonstrated equivalent expression GFP-SET8 fusion in all transfected samples. Similarly, anti-FLAG antibody probing of the blot showed robust expression of FLAG-PARP1 fusion protein (Fig. 3A, upper panel). Next, we immunoprecipitated GFP-SET8 fusion with anti-GFP antibody and probed the western blot with anti-HA to detect HA-ubiquitinated SET8 fusion or anti-PAR for poly ADP-ribosylated SET8 fusions. Indeed, GFP-SET8 showed high molecular weight HA-ubiquitinated smear when expressed alone with HA-ubiquitin (Fig. 3A, lower left panel, lane 1). GFP-SET8 co-expression with FLAG-PARP1 and HA-ubiquitin resulted in accumulation of high molecular weight HA-ubiquitinated smear, suggesting that the expression of additional PARP1 enhances ubiquitination of GFP-SET8 (Fig. 3A, lower left panel, lane 2, 1.6x).

**Figure 3:**
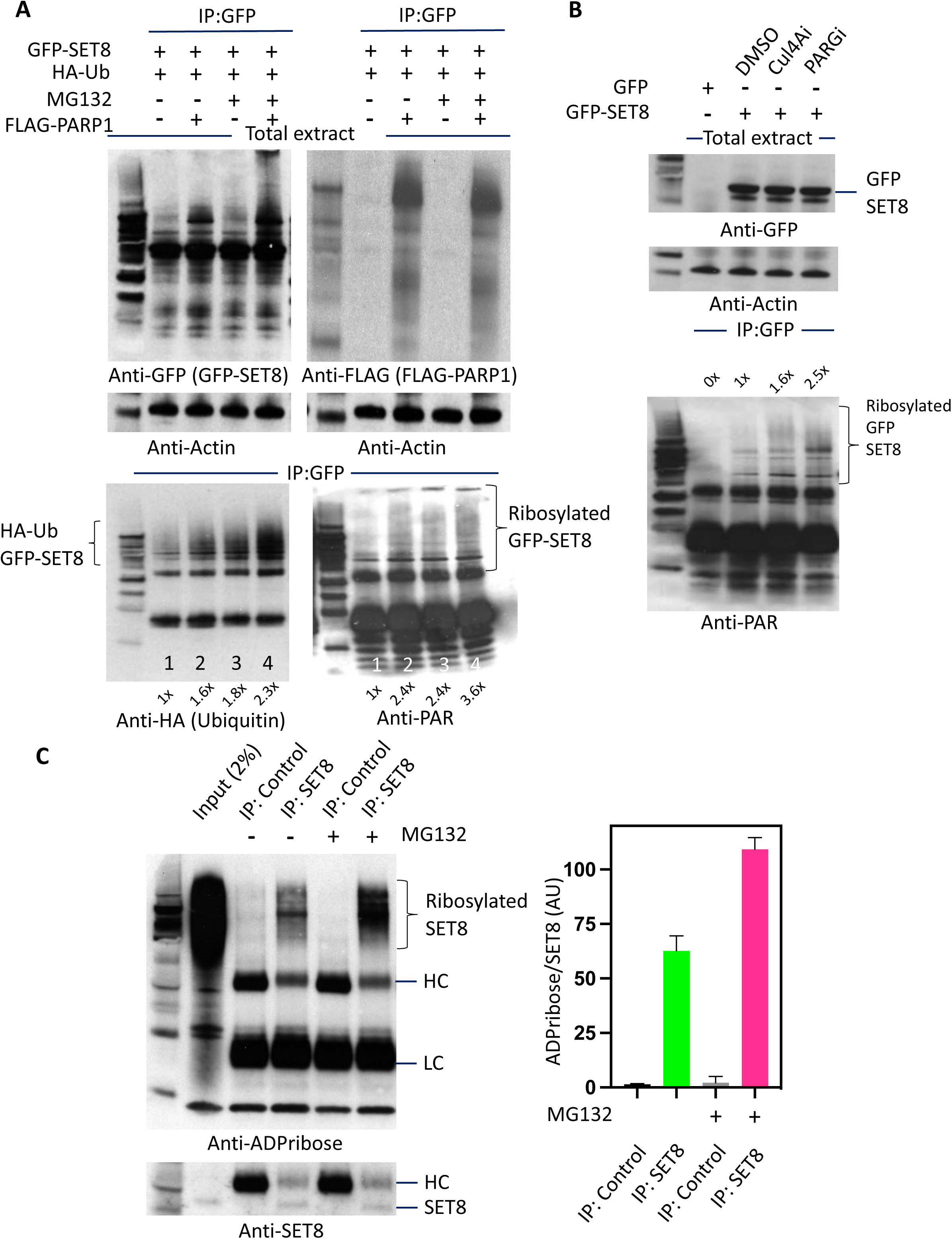
PARP1 promotes SET8 protein degradation. (A) GFP-SET8 immunoprecipitation in overexpressed GFP-SET8 COS-7 cells with or without FLAG-PARP1 overexpression in presence or not of the proteasome inhibitor MG132. Western blots detecting the amount of GFP-SET8 protein overexpressed in total extract (top, left) as well as the amount of FLAG-PARP1 protein overexpressed (top, right) using anti-GFP and anti-FLAG antibody respectively. Western blots detecting the amount of ubiquitin (Ub) (bottom, left) or ADP-ribosylation (bottom, right) of immunoprecipitated GFP-SET8 protein using anti-HA and anti-ADP ribose antibody respectively. Anti-actin was used as a loading control (middle). The fold increase was calculated by densitometry and indicated at the bottom of the western blot. (B) GFP-SET8 immunoprecipitation in overexpressed GFP-SET8 COS-7 cells in presence or not (DMSO) of Cullin inhibitor (Cul4Ai) or PARG inhibitor (PARGi). Western blot detection of total GFP-SET8 protein levels overexpressed in COS-7 cells (top). Anti-actin antibody was used as control (middle). Western blot detecting the amount of poly-ADP ribosylation (bottom) of immunoprecipitated GFP-SET8 protein using an anti-ADP ribose antibody. (C) SET8 immunoprecipitation in HeLa cells in presence or not of the proteasome inhibitor MG132. Western blot (left side) detecting the amount of ADP-ribosylation in GFP (control IP), SET8 immunoprecipitates (top panel) as well as the amount of SET8 protein (bottom panel). Respective densitometry analyses of SET8 ADP-ribosylation abundance representative of at least 2 biological experiments are shown (right side).

In addition, reducing the degradation of ubiquitin-conjugatedof GFP-SET8 by MG132 resulted in higher amounts of high molecular weight HA-ubiquitinated smear as expected. When we co-expressed GST-SET8, FLAG-PARP1 and HA-ubiquitin and reduced protein degradation using MG132, we observed ∼ 5 folds accumulation of HA-ubiquitinated GST-SET8 protein (Fig. 3A, lower left panel, lane 4, 2.3x). Similarly, ribosylated GFP-SET8 was more prominent in GFP-SET8 and FLAG-PARP1 overexpressed cells (Fig. 3A, lower right panel, lane 2 and 4, 2.4x and 3.6x respectively). All these experiments conclusively prove that PARP1 is an effector protein that aids in poly ADP-ribosylation of SET8 leading to ubiquitin-mediated degradation.

The SET8 is ubiquitinated on chromatin by CRL4(Cdt2) complexes during S phase and following DNA damage in a PCNA-dependent manner. In a transgenic mouse model for lung cancer, the level of SET8 was reduced in the preneoplastic and adenocarcinomous lesions following over-expression of Cul4A (62). Therefore, it is expected that down regulation of Cul4A by siRNA would lead to accumulation of SET8. Similarly, downregulation of PARG, that removes ADP ribose from poly ADP-ribosylated protein by PARP1 and acts as an antagonist of PARP1, would also lead to the accumulation of SET8. To validate our hypothesis, we transfected GFP-SET8 fusion constructs to mammalian cells and treated the cells with DMSO, Cul4A inhibitor (MLN4924), or PARG inhibitor (PDD00017273) and monitored the SET8 pattern of expression by immunoprecipitation and western blotting. Indeed, GFP-SET8 fusion protein was expressed in a similar level in the absence or presence of either Cul4A or PARG inhibitor (Fig.3B, top panel). Immunoprecipitation of GFP-SET8 showed more prominent high molecular weight poly ADP-ribosylated GFP-SET8 fusion in PARGi or Cul4Ai -treated cells (Fig. 3B, bottom panel).

We next investigated if poly ADP-ribosylation indeed occus naturally in endogenous SET8 enzyme. We treated HeLa cells with MG132 to reduce protein degradation and immunoprecipitated SET8 with anti-SET8 antibody. The control antibody was anti-GFP. The immune precipitates were separated on SDS-PAGE, western blotted and probed with anti-ADPribose and anti-SET8. Immunoprecipitated samples displayed a strong high molecular weight smear of SET8, although MG132 treated sample had higher intensity confirming poly-ADP ribosylation is crucial for SET8 degradation (Fig. 3C, left panel). Quantititive measurements of relative band intensities showed ∼40% more signal intensity in the presence of MG 132 for SET8 (Fig. 3C, right panel). Taken together with results from figure 3A-C, we correlate that high molecular weight SET8 or its GFP fusion are both ubiquitinylated and poly ADP-ribosylated, suggesting a cross-communication between both post translational modifications to maintain a steady-state level of SET8 in the cells.

### Poly ADP-ribosylation and ubiquitinylation regulate SET8 levels and H4K20 methylation

Since there was a correlation between poly ADP-ribosylation and ubiquitinylation of SET8 molecules, we transfected both GFP-SET8 FL or GFP-SET8 M in to HeLa cells in the presence of HA-ubiquitin plasmid and monitored levels of fusion SET8, ADP-ribosylated fusion SET8 and ubiquitinylated-SET8. Indeed, wild type fusion displayed prominent high molecular weight ladder pattern indicating poly ADP-ribosylated fusion SET8, correlating with ubiquitinylated-SET8 and the levels were higher ∼3.6x compared to the mutant enzyme, since the acceptor Lys residues were mutated (Fig. 4A). This led us to hypothesize there is a synergistic effect of poly ADP-ribosylation and poly ubiquitinylation in SET8 stability. We performed half-life measurement for both GFP-SET8 FL and GFP-SET8 M in the presence of cycloheximide. Indeed wild type SET8 fusion degraded much faster compared to the mutant enzyme, although the loading control, actin remain constant throughout the time course. The wild type SET8 had a half-life of 3.8 hrs compared to 12.3 hrs for the mutant (Fig. 4B). Therefore, it is plausible that the lysine residues in the disordered domain are crucial in maintaining SET8 enzyme levels in cell.

**Figure 4:**
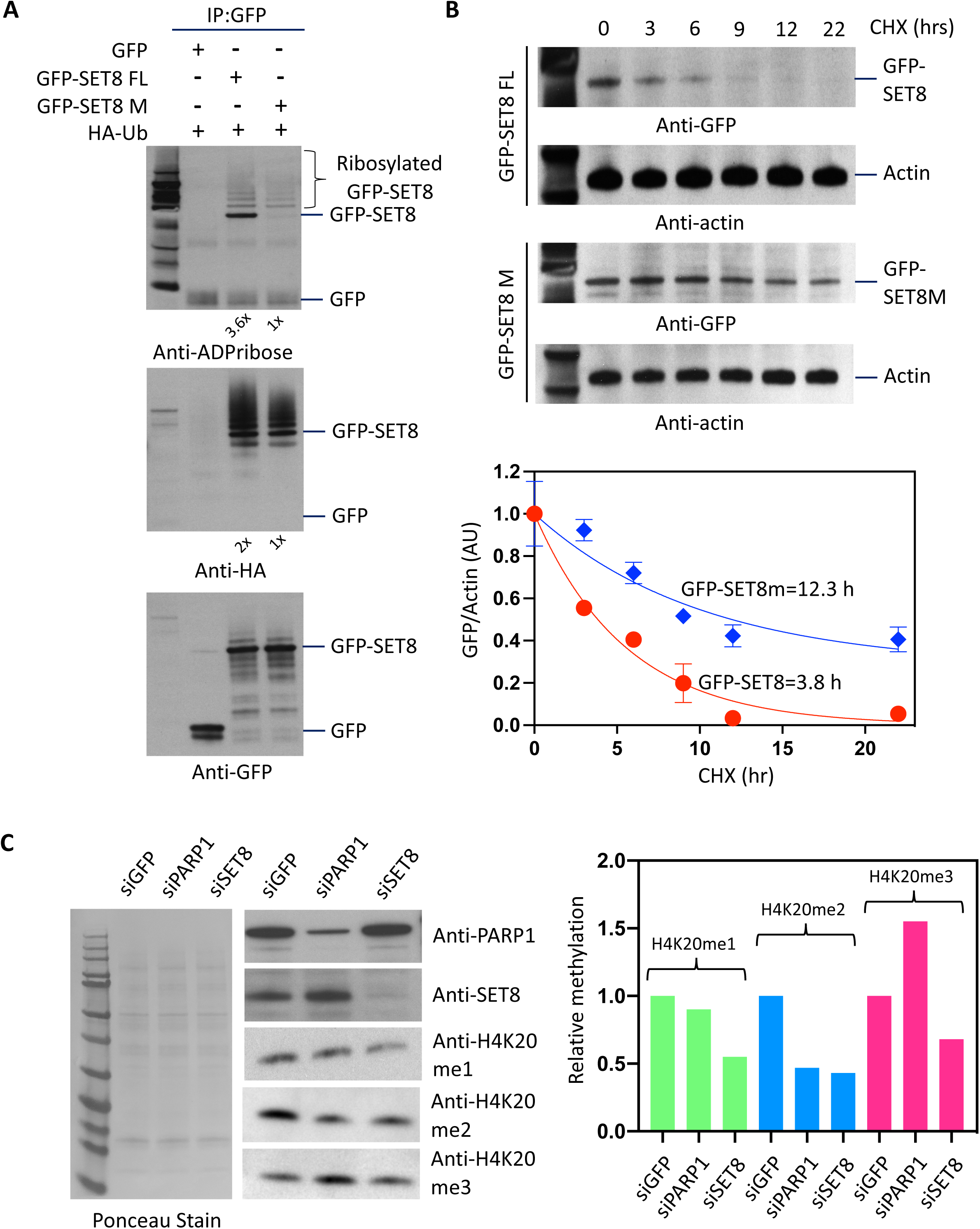
PARP1 regulates SET8 protein stability. (A) GFP-SET8 immunoprecipitation in overexpressed GFP-SET8 or GFP-SET8m with HA-Ubiquitin in COS-7 cells. Western blots detecting the amount of ADP ribosylation (top panel), Ubiquitination (middle panel) and GFP fusion protein levels (bottom panel) in GFP immunoprecipitates. The fold increase was calculated by densitometry and indicated at the bottom of the western blot. (B) Cycloheximide chase analysis of SET8 stability in GFP-SET8 or GFP-SET8m in HeLa cells. Western blots detecting the amount of GFP-SET8 and GFP-SET8m protein levels with their respective actin levels (control) during cycloheximide time course (top panel). Respective densitometry analyses of GFP/Actin ratio representative of at least 2 biological experiments (bottom panel). (C) Western blots (left side) detecting the amount of PARP1 (top), SET8 protein (middle) as well as the amount of H4K20me1, H4K20me2 and H4K20me3 levels (bottom) in total protein extract of knockdown HeLa cells treated with esiRNA GFP (control), esiRNA PARP1 and esiRNA SET8, respectively. Respective densitometry analyses of protein abundance representative of at least 2 biological experiments are shown (right side). Ponceau stain was used as control (left side).

We next knocked down PARP1 using siRNA to define its role in SET8 stability conclusively. HeLa cells were treated with control siGFP, siPARP1 and siSET8 and the extracts were western blotted and probed with respective antibodies. As expected, PARP1 siRNA was able to knock down 70% PARP1 resulting in 1.5-fold increase in SET8 protein level but not SET8 mRNA level confirming that SET8 increase was due to post translational modification (Table S3). Surprisingly, SET8 level increase did not translate into an increase in global H4K20me1 level (Fig. 4C). This led us to investigate if global H4K20me2/3 heterochromatic marks are affected in the knockdown cells using western blot. Surprisingly, H4K20me2 level decreased along with the concurrent gain of global H4K20me3 suggesting knockdown of PARP1 facilitates the rapid formation of H4K20me3 that may result in aberrant heterochromatic mark establishment following cell cycle.

### Cell cycle dependent interaction of SET8 and PARP1

Since siPARP1 led to an increase in H4K20me3, and precursor H4K20me1 and SET8 enzyme are crucial to cell cycle progression genome stability, DNA replication, mitosis, and transcription, we hypothesized that SET8 and PARP1 would be colocalized in cells in a cell cycle-dependent manner for SET8 dynamics on chromatin. Lovastatin and thymidine-nocodazole treated synchronized HeLa cell nuclei were isolated and extracts were made corresponding to G1, S and G2/M phase. Equal amounts of extracts were evaluated for the relative abundance of SET8, PARP1, and H4K20me1 using western blot with respective antibodies. CDT1 (G1 phase marker) was used as a positive control for cell cycle synchronization. As expected, SET8 level was lowest during S phase and highest at G2/M phase, mirroring H4K20me1 levels. PARP1 level remained unchanged throughout the cell cycle (Fig. 5A). Since SET8 and PARP1 directly interact, we investigated SET8 levels on the chromatin. To validate this hypothesis, we used G1, S, G2/M nuclear extracts for SET8 immunoprecipitation (IP). The captured proteins were resolved on SDS-PAGE, western blotted and probed with SET8 and PARP1 antibodies (Fig. 5B). We observed the highest amounts of SET8 being captured in G2/M compared to S or G1 phase. SET8 IP also revealed that PARP1 is co-immunoprecipitated in G2/M compared to G1 or S phase. However, all three phases with PARP1 co-immunoprecipitated with SET8, abate a higher amount during S phase. We, therefore, quantified relative co-immunoprecipitation between PARP1-SET8 during all three cell cycle phases by comparing the ratios between PARP1 and SET8 in 3 independent experiments. Indeed, PARP1/SET8 ratio was highest during S phase compared to the lowest in G2/M (Fig. 5B, bottom panel). Concurrently, in the same IP samples we also measured the ribosylated SET8 enzymes, the substrate of PARP1 (Fig. 5C). Indeed, the ratio of PARP1 and SET8 mirrored the ratio between ADP ribosylated SET8 and SET8 confirming PARP1 indeed modulates SET8 ribosylation which impacts the stability of the enzyme on the chromatin in a cell cycle-dependent manner (Fig. 5B vs. C). This would reflect SET8 enzyme level to global H4K20me1 during cell cycle.

**Figure 5:**
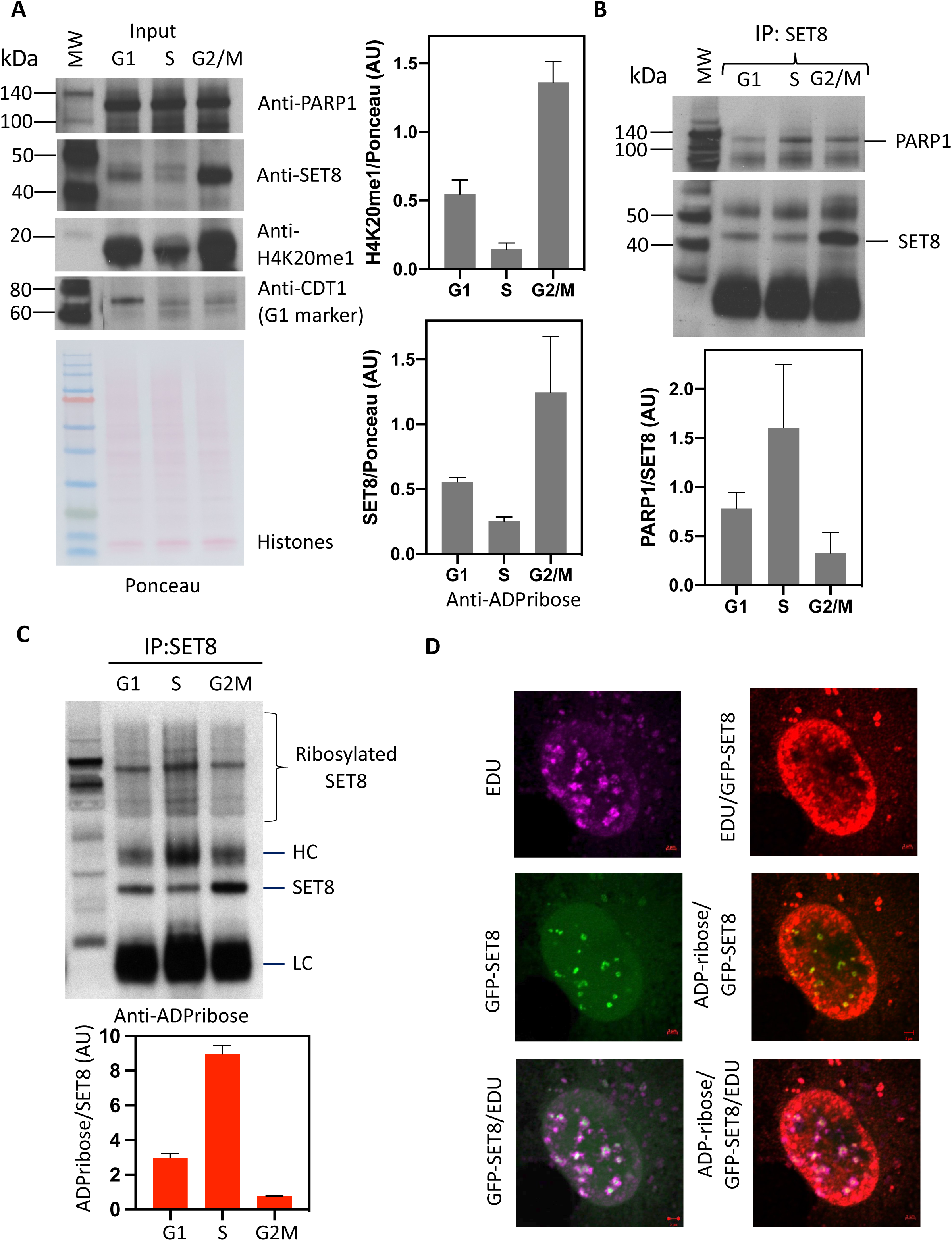
Cell cycle dependent interaction of PARP1 and SET8 correlates with global H4K20me1. (A) Western blot indicating PARP1 (top, left), SET8 (middle, left) and H4K20me1 (middle, left) levels in total protein extracts from HeLa cells synchronized in G1, S and G2/M phases, respectively. Western blot of CDT1 protein levels is shown as a cell cycle synchronization control as well as Ponceau stain for loading control and densitometry analyses (bottom, left). Respective densitometry analyses of H4K20me1 (top, right) and SET8 (bottom, right) relative protein abundances are shown (right) and representative of at least 2 biological experiments. (B) SET8 immunoprecipitation from total protein extract in HeLa cells synchronized in G1, S and G2/M phases. Western blots detection of PARP1 (top, left) as well as SET8 immunoprecipitated protein levels (bottom, left) are revealed. Densitometry analyses of PARP1/SET8 ratio during G1, S and G2/M cell cycle phases are shown (right) and are representative of at least 2 biological experiments. (C) SET8 immunoprecipitation from total protein extract in HeLa cells synchronized in G1, S and G2/M phases. Western blots detection of ADP-ribosylation as well as SET8 protein levels in SET8 immunoprecipitates (top panel). Respective densitometry analyses of SET8 ADP-ribosylation abundance representative of at least 2 biological experiments are shown (bottom panel). (D) Pulsed chased cells with 5-ethynyl-2′-deoxyuridine (EdU) to label DNA (magenta) is transfected with GFP-SET8 (green). Endogenous ADP-ribose (red) is revealed by anti-ADP-ribose conjugated with Texas Red. Merged images demonstrates the co-localization of EDU, SET8 and ADP-ribose.

To observe ADP ribosylation during cell cycle, we pulse changed COS-7 cells with 5-ethynyl-2’- deoxyuridine (EdU) to selectively labeled the newly synthesized DNA in S phase, for rapid visualization and its ability not to interfere with subsequent antibody staining. The same cells were also transfected with GFP-SET8 and further probed with anti-PAR antibody. EdU formed the punctate pink nuclear staining pattern, typical of S phase. And the same regions were also visualized as yellow indicating GFP-SET8 colocalization. Poly ADP-ribosylation was visualized using anti-PAR antibody as red throughout the nucleus, as expected with clear colocalization at the replication forks and SET8 (Fig. 5D). These results indicate poly ADP-ribosylation co-exists with SET8 on DNA replication forks and correlate with high level of SET8 ADP-ribosylation detected by IP (Fig. 5C). Thus, we conclude that PARP1 plays a central role during cell cycle to dynamically regulate H4K20 methylation.

### PARP1 level regulates H4K20me1 and H4K20me3 Chromatin Domains

Since PARP1 predominantly interacts with SET8 during S phase leading to a concurrent decrease in SET8 and H4K20me1 level, we next investigated if there is a functional implication of PARP1 level in the cell and H4K20me1/me3 distribution on the chromatin. For this investigation, we knocked down PARP1 in HeLa cells and performed H4K20me1 and H4K20me3 ChIP-seq with respective siGFP controls. Prior to the ChIP-seq, we western blotted the samples and observed similar levels of SET8 and H4K20me1, me2 and me3 as observed for PARP1 knockdown (Fig. 4C). The Spearman correlation between the ChIP-seq fragments between control siGFP vs. knockdown siPARP was 1.95 and 0.95 for H4K20me1 and H4K20me3 respectively, demonstrating high degree of similarity. As expected, irrespective of siGFP or siPARP knockdown the correlation values between H4K20me1 and H4K20me3 remained below 0.37 suggesting both marks are mutually exclusive with a small percentage overlap. However, there was no significant similarity between H4K20me1 and me3 (Fig. 6A). We also mapped the H4K20me1 and me3 peaks and observed that PARP1 knockdown has higher read densities compared to their respective controls suggesting H4K20me1 hypermethylation in the peak regions and vicinity (-/+ 5 kbp), a similar pattern was observed for H4K20me3 hypermethylation (Fig. 6B-C). We performed peak annotations to determine the genomic elements associated with both post-translational marks. Indeed, LINE, SINE, intron elements displayed appreciable amounts of hypermethylation for both H4K20me1 and me3. However, the satellites displayed loss of H4K20me1, and gain of H4K20me3 regional density in response to PARP1 knock down, suggesting PARP1 levels have a different mechanism in maintaining the methylation status (Fig. S14). The Spearman’s correlation between the ChIP-seq peaks between H4K20me1 and H4K20me3 for the above sets of genomic features remained below r=0.5 for LINE, SINE and intergenic regions. Surprisingly, the satellite regions had high degree of correlation indicating both H4K20me1 and me3 are enriched there as reported previously (Fig. S15; 63, 64).

**Figure 6:**
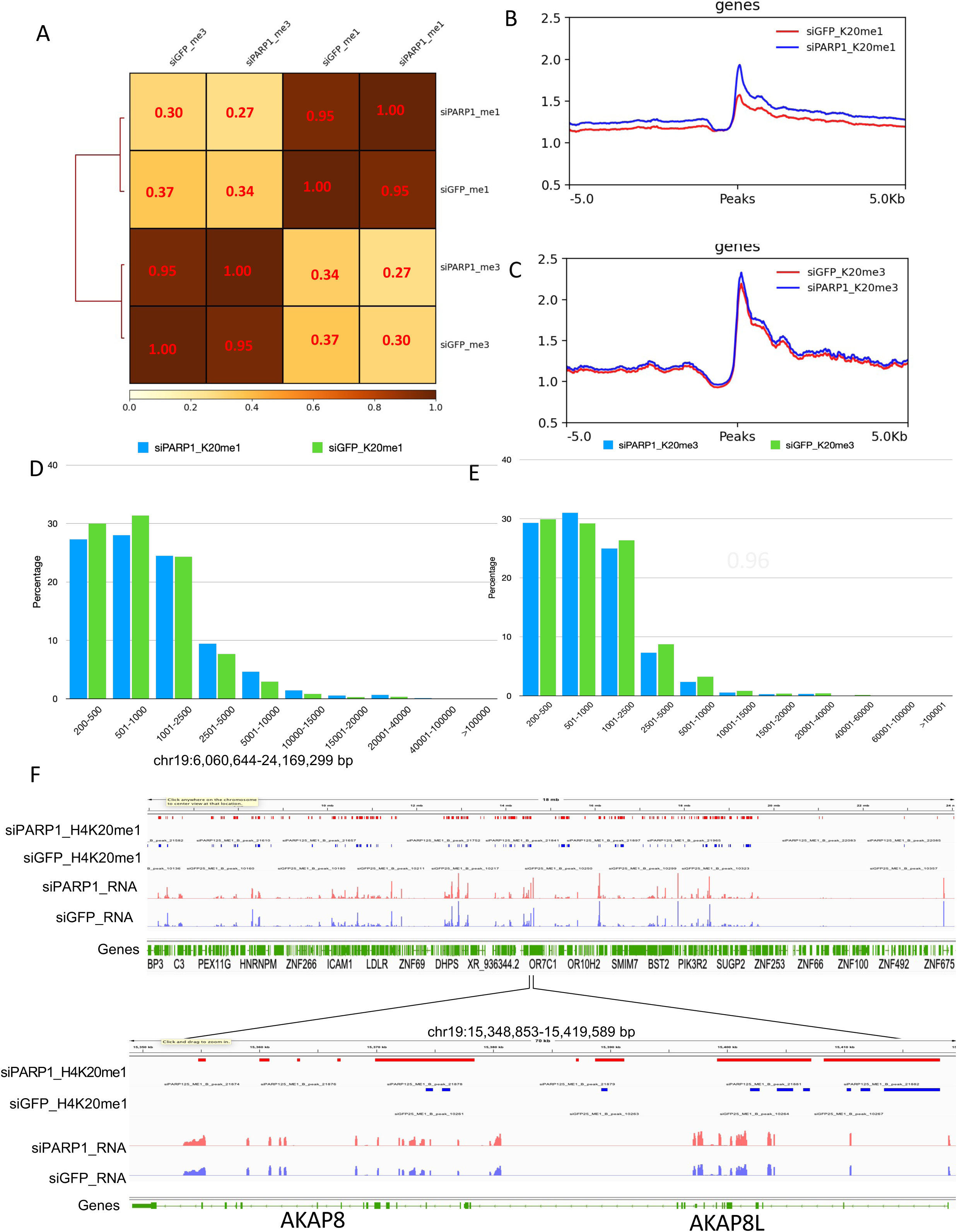
PARP1 regulates H4K20me1 and H4K20me3 distribution in the genome. (A) Spearman correlation of H4K20me1 and H4K20me3 regions in PARP1 knockdown HeLa cells and its control. (B) Genome-wide metagene plot showing H4K20me1 profile (ChIP-seq) in control and PARP1 knockdown HeLa cells. (C) Genome-wide metagene plot showing H4K20me3 profile (ChIP-seq) in control and PARP1 knockdown HeLa cells. (D) Peak width profile of H4K20me1 by binning the peaks into different lengths in PARP1 knockdown cells and its control. (E) Peak width profile of H4K20me3 by binning the peaks into different lengths in PARP1 knockdown cells and its control (F) Representative IGV genomic tracks showing H4K20me1 and RNAseq profile in PARP1 knockdown cells and its control.

Further we binned various peak width of H4K20me1 and H4K20me3 and observed that H4K20me1 distribution reduced below 1 kb peak width with concurrent gain up to 10 kb in response to PARP1 knock down (Fig. 6D). Similarly, H4K20me3 distributions was more prominent on 500-1000 bp (Fig. 6E). This suggested that the knock down of PARP1 may shift the dynamic equilibrium between both marks and thus changing the chromatin domains. Indeed, we observed more dramatic changes are in H4K20me1 domains. IGV browser displayed changes in H4K20me1 boundaries throughout the genome upon PARP1 knockdown compared to control knockdown correlating with gene expression (Fig. 6F). To study the global gene expression corresponding to H4K20me1, ChIP-seq derived sequences annotated genes identified. The percentage overlap of the genes is shown based on the intersecting genes taken from respective RNA-seq and the intragenic regions of H4K20me1 annotated files. The intersecting genes in various conditions, siPARP1_K20me1, siGFP_K20me1, were 9.59%, 4.89% respectively confirming H4K20me1 has 2- folds more association with transcriptionally active genes (Fig. S16A). Indeed, the logarithmic distribution of the positively expressed genes from ChIP-seq of H4K20me1 show more compatibility with global gene expression in siPARP1 cells as displayed in the density plots compared to control siGFP (Fig. S16B).

### Enrichment of consensus TF-binding motifs near H4K20me1/me3 affecting chromatin state

Further analysis of the sequence tags obtained from H4K20me1 and H4K20me3 ChIP-seq and the consensus transcription factor binding sites analysis shows that PARP1 knockdown in cells leads to enrichment or decrease of some transcription factor (TF) binding sites in its vicinity. Enrichment or decrease of those TF binding sites are associated with the Wnt/beta-catenin and TGF-beta pathway (Fig. S17A, B; Table S4A, S4B). For example, TCF7, a transcriptional repressor implicated in the Wnt pathway by directly binding to beta-catenin was observed to be PARP1 regulated via H4K20me1/3. Similarly, SMAD4 and STAT5, two central TGF-beta transcription factors were PARP1 dependent. The PARP1 dependent TF enrichment sequences with H4K20me1/me3 chromatin signature fell into three general categories. (1) Overall PARP1 dependent TF binding site enrichment, described as enriched in control cells, but not when PARP1 depleted. (2) PARP1 independent TF binding site enrichment, described as no enrichment in cells when PARP1 is depleted. (3) PARP1 inhibited TF binding site enrichment, described as enriched in cells when PARP1 is depleted (Fig. S12A, B). However, significant numbers of TF enrichment were dependent on H4K20me1 or H4K20me3 marks. Therefore, it is plausible that PARP1 mediated mis-regulation of H4K20me1 and H4K20me3 domains are linked to gene regulation via transcription factor recruitment (Table S4A and S4B).

In chromatin state analysis studies PARP1 depletion positively affected H4K20me1 levels leading to various transcription states, particularly for transcription at 5’ and 3’, enhancers, strong and weak transcription. Similarly, H4K20me3 enriched states were more prominant at genic enhancers, enhancers, ZNF genes and repeats (Fig. S18). Therefore, these observations suggest that optimal PARP1 level is essential for TF motif enrichment and chromatin state maintenance that could affect gene expression and cellular development.

## Discussion

The principal enzyme of poly ADP-ribosylation is PARP1. In this study, poly ADP-ribosylation not only impaired SET8 catalytic activity, it also ribosylated and facilitated SET8 degradation by the ubiquitin degradation pathway. As expected, knockdown of PARP1 resulted in accumulation of SET8 in cells, although it did not increase global H4K20me1, the enriched peaks displayed small hypermethylation at proximal and distal regions of the peaks. However, there was a profound decrease of global H4K20me2 and corresponding increase in global H4K20me3. This is not unexpected since H4K20me2 is the precursor of heterochromatin mark H4K20me3. In gene annotation analysis, the satellite DNA that comprises 10-15% of the human genome showed a decrease in H3K20me1 and corresponding increase of H4K20me3. This was further supported by the FLAG-PARP1 overexpression studies. Indeed, overexpression of PARP1 not only decreased the level of SET8, it also correspondingly decreased the level of H4K20me1, H4K20me2 and H4K20me3. A reasonable explanation would point that the knockdown cells were mostly in resting phase with saturated amounts of H4K20me1 on the genome. And secondly, the decrease in H4K20me2 and concurrent increase in H4K20me3 may be as a result of another methylase that is catalytically regulared by PARP1, which would need additional studies. Taken together, these results suggest an alteration in the level of SET8 in cells can profoundly impact heterochromatic mark H4K20me3.

An increase in H4K20me3 may have resulted in heterochromatic domain rearrangement in overexpressing cells. While many studies have reported that PARP1 promotes gene transcription, it also promotes gene expression at the post-transcriptional level by modulating the RNA-binding protein HuR (65, 66). Indeed, PARP-1 is required for a series of molecular outcomes at the promoters of PARP-1 regulated genes, leading to a permissive chromatin environment for RNA Pol II machinery loading. Poly ADP-ribosylation has an important role in the maintenance of H3K4me3, as the enzyme for demethylation, KDM5B, is impaired by poly ADP-ribosylation. Consistently, an increased level of KDM5B at TSS of active genes is associated with decreased H3K4me3 after inhibition of poly ADP-ribosylation *in vivo*. (67). There is also *in vitro* evidence of a direct involvement of poly ADP-ribosylation in the crosstalk between H3 and H1 methylation.

Indeed, poly ADP-ribosylation of H3 impairs its methylation by the H3K4 mono-methyltransferase SET7/9, thus shifting its catalytic activity towards other lysine residues of H1 (68). Another role of poly ADP-ribosylation can contribute to transcriptional repression by H3K9me2 accumulation at retinoic acid (RA)-dependent genes. In this mechanism, demethylase KDM4D is covalently poly ADP-ribosylation at the N-term domain, impairing its recruitment onto RA-responsive promoters, leading to repression by H3K9me2 accumulation (69). Apart from the transcriptional role of PARP, inhibition/depletion of the enzymes also causes loss of epigenetic marker on heterochromatin, H3K9me3 (70), H4K20me3 (71), and 5mC (72) at the centromeric regions. These all studies suggest that PARP enzymes can modulate chromatin structure and regulate gene expression.

However, previous studies have reported poly ADP-ribosylated PARP1 might conflict with CG methylation by non-covalent interaction with DNMT1 preventing its access on DNA and catalysis (72). PARP1 also affects DNA methylation by forming a complex with the transcription factor CTCF. In another report, PARP1 was shown to control the UHRF1-mediated ubiquitination of DNMT1 to timely regulate its abundance during S and G2 phase, thus impacting CG methylation. Therefore, the above observations, and our current studies, demonstrate other epigenetic marks particularly DNA methylation and SET8 levels in the cell, may play a role in H4K20me1 and H4K20me3 chromatin domains. Indeed, SET8 is known to regulate chromatin compaction during G1 phase (18). Therefore, it would make sense that during this phase there is little interaction between PARP1 and SET8. Indeed, upon mitotic exit, chromatin relaxation is controlled by SET8- dependent methylation of histone H4K20. In the absence of either SET8 or H4K20me1 mark, substantial genome-wide chromatin decompaction occurs allowing excessive loading of the origin recognition complex (ORC) in the daughter cells (18). Based on these results, it is plausible that PARP1 overexpression in cancer would catalytically compromise SET8 leading to slight reduction in H4K20me1 mark. This would have a larger ramification in an increase of H4K20me3 level on heterochromatin. Since H4K20me3 represses transcription when present at promoters and silences repetitive DNA and transposons, its aberrant deposition on chromatin may alter the transcriptional network during oncogenesis. Comparison of H4K20me1 ChIP seq with RNAseq also reflects the impact of the H4K20me1 on active transcription (Fig S16A) which is proportionate with the amount of H4K20me1. This was also demonstrated on housekeeping genes, chromatin openness, etc. (73). Furthermore, SET8 mediated H4K20me1 regulates Pol II promoter-proximal pausing by regulating H4K16ac and H4K20me3 levels and ultimately transcriptional output (74). Therefore, PARP1 mediated poly ADP-ribosylation can impact epigenome inheritance, chromatin structure, and transcription factor occupancy.

In summary, our study reveals a novel mechanism of SET8 mediated epigenome regulation. Poly ADP-ribosylation not only can catalytically impare the enzyme activity, it also trigger a series of events that allows ubiquitinylation of SET8 molecules in the cells leading to its final degradation. It remain to be seen if poly ADP-ribosylation acts as an allosteric activator for ubiquitin ligases of SET8. Since down regulation of SET8 inhibit progression of hepatocellular carcinoma, these insite may aid in designing SET8 inhibitors that may be useful for cancer treatment (76). Furthermore, data presented herein may provide insight into studies of PAR-dependent epigenome regulation.

## Data availability

ChIP-seq data performed in this study are available in NCBI Gene Expression Omnibus (GEO) under the accession GSE188744

## Acknowledgments

We thank C. Carlow for critical reading of the manuscript, T. Evans, D. Comb, Sir R.J. Roberts and J. Ellard for encouragement.

## Author contribution

POE, CR, HGC performed experiments. VUS and SS performed data analysis. SP conceptualized the project and wrote the manuscript in collaboration with POE, CR, HGC and VUS.

## Supplementary figures

**Figure S1:**
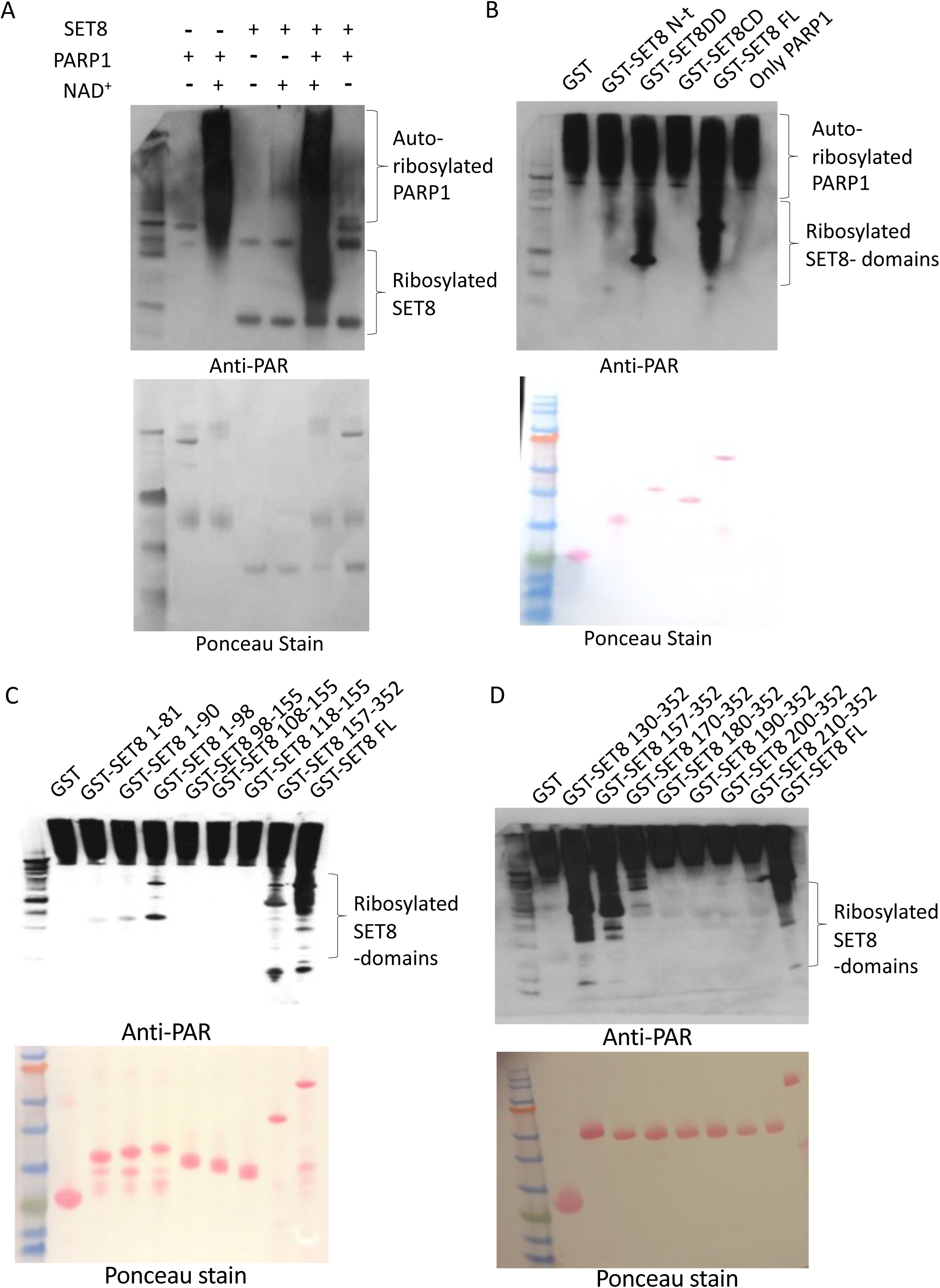
Detail poly ADP-ribosylation mapping of SET8. (A) Full length SET8 is poly ADP ribosylated by PARP1. The enzymatic in vitro reaction was separated using SDS-PAGE, western blotted and probed with anti-PAR antibody. The shifted SET8 band shows the ADP ribosylated SET8 (top). Loading of the exact blot shown using ponceau staining (bottom). (B) ADP-ribosylation of various SET8 domain in the presence of PARP1. The ribosylated SET8 GST-SET8 N-t (1-98), GST-SET8 DD (157-352) domains as well as GST-SET8 FL are shown by western blotting using anti-PAR antibody (top). Loading of the exact blot shown using ponceau staining (bottom). (C) ADP-ribosylation of various GST-SET8 regions in the presence of PARP1. GST-SET8 N-t (1-98), GST-SET8 DD (157-352) and GST-SET8 FL show ribosylation (top). Loading of the exact blot shown using ponceau staining (bottom). (D) Detailed ADP-ribosylation analysis of various truncated SET8 regions between amino acids 130-352 in the presence of PARP1. GST-SET8 (130-152), GST-SET8 (157-352), GST-SET8 (170-352) as well as GST-SET8 FL shows ADP-ribosylation (top). Loading of the exact blot shown using ponceau staining (bottom).

**Figure S2:**
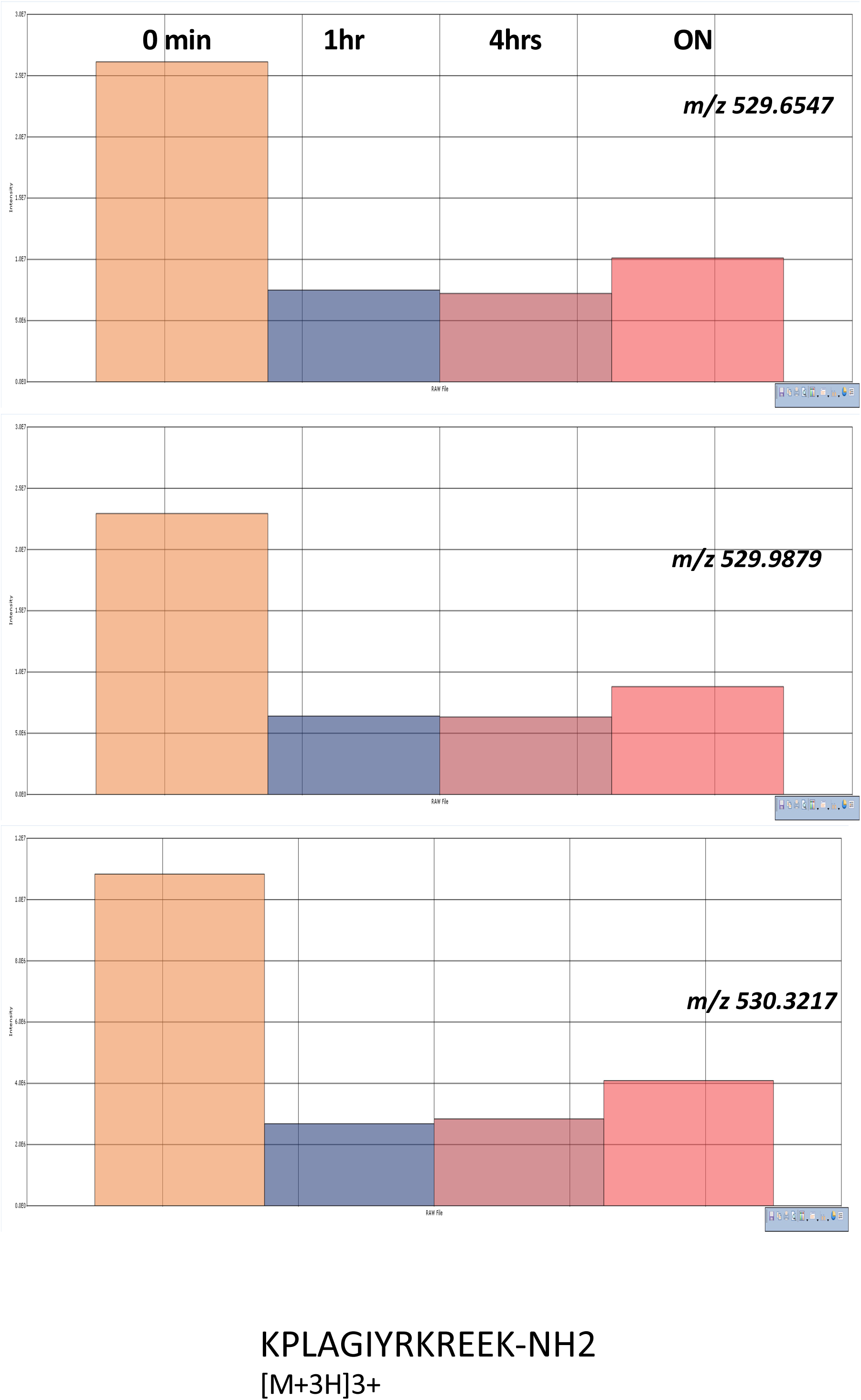
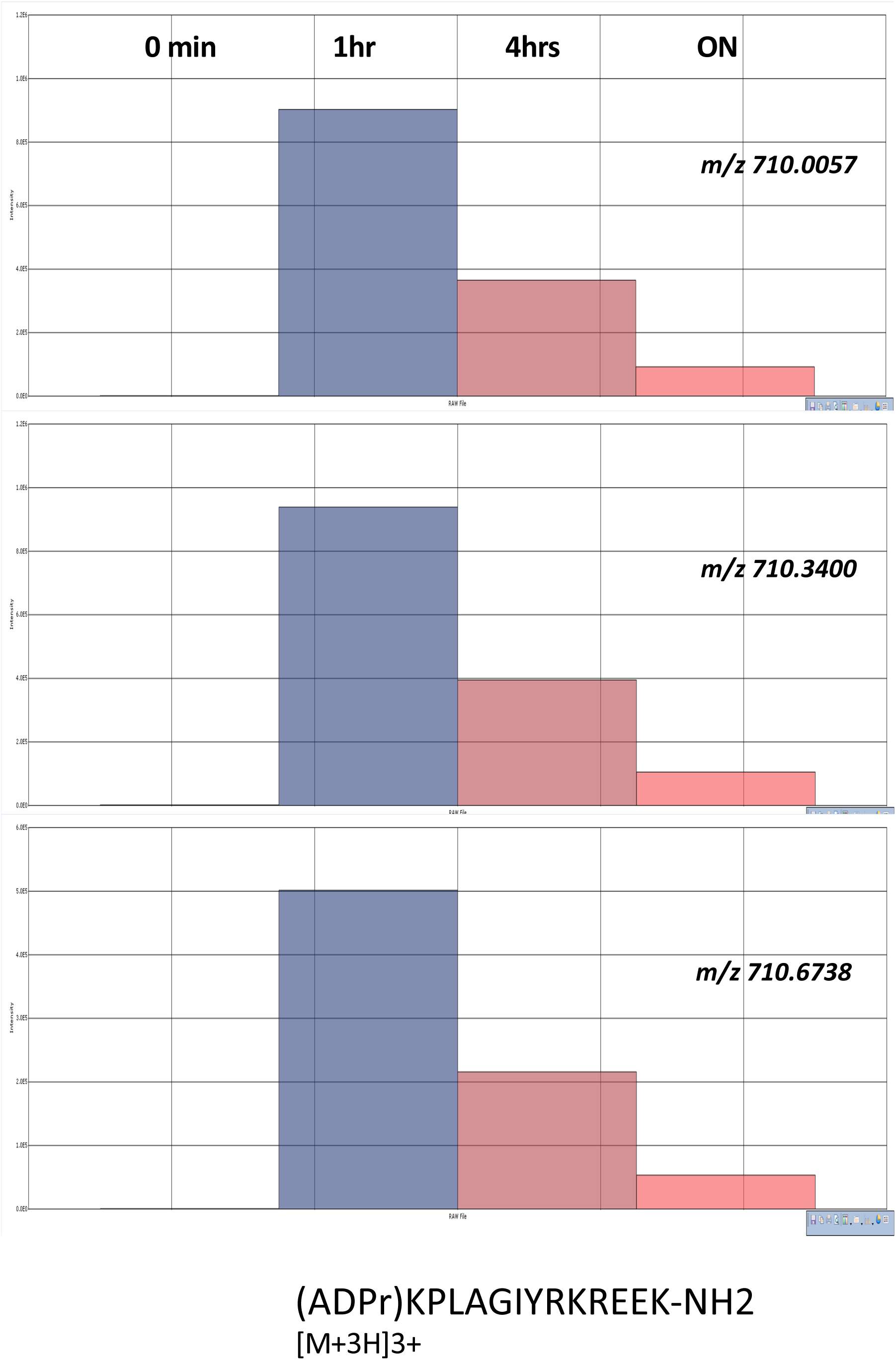
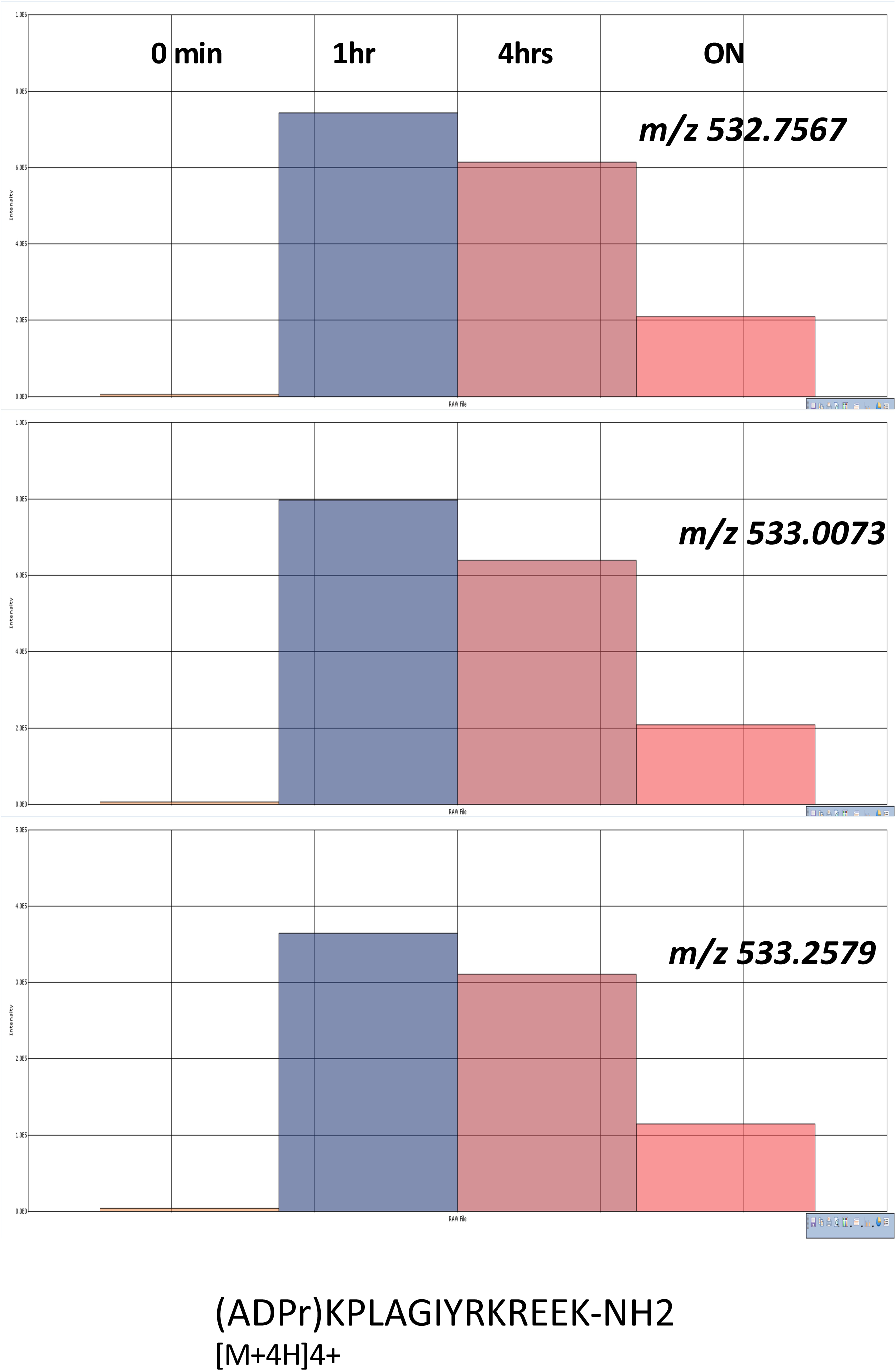
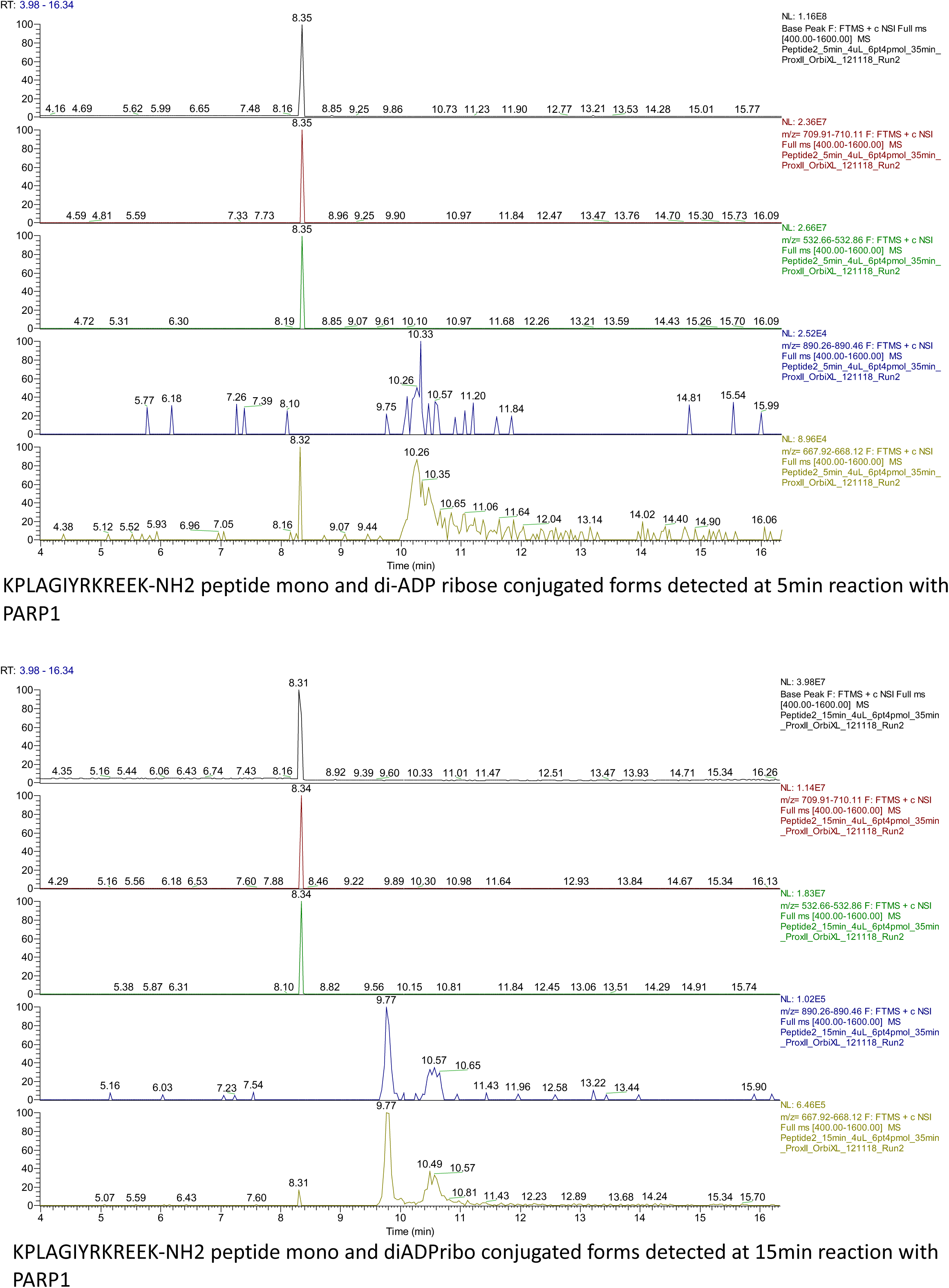
Characterization of ADP-ribosylation of SET8 synthetic peptide KPLAGIYRKREEK-NH2 using high resolution mass spectrometry and isotope quantification in nanoLC. (A) Quantification of peptide KPLAGIYRKREEK-NH2 (charge +3) at four time points: 0 min (negative control), 1hr, 4rhs and overnight (ON) reaction with PARP1. Peak areas were determined with SIEVE software package for the three main isotopes m/z 529.65449, m/z 529.98877 and m/z 530.32300, theoretical values determined with Protein Prospector MS Isotope. The figure displays the measured isotopes 0, 1 and 2 with Orbitrap MS. (B) Quantification of peptide (ADPr)KPLAGIYRKREEK-NH2 (charge +3) at four time points: 0 min (negative control), 1hr, 4rhs and overnight (ON) reaction with PARP1. Peak areas were determined with SIEVE software package for the three main isotopes m/z 710.00819, m/z 710.34247 and m/z 710.67670, theoretical values determined with Protein Prospector MS Isotope. The figure displays the measured isotopes 0, 1 and 2 with Orbitrap MS. (C) Quantification of peptide (ADPr)KPLAGIYRKREEK-NH2 (charge +4) at four time points: 0 min (negative control), 1hr, 4rhs and overnight (ON) reaction with PARP1. Peak areas were determined with SIEVE software package for the three main isotopes m/z 532.75796, m/z 533.00867 and m/z 533.25934, theoretical values determined with Protein Prospector MS Isotope. The figure displays the measured isotopes 0, 1 and 2 with Orbitrap MS. (D) Chromatographic profile of the main isotopes of (1ADPr) KPLAGIYRKREEK-NH2 and (2ADPr) KPLAGIYRKREEK-NH2 detected at 5 min reaction with PARP1. (E) Chromatographic profile of the main isotopes of (1ADPr) KPLAGIYRKREEK-NH2 and (2ADPr) KPLAGIYRKREEK-NH2 detected at 15 min reaction with PARP1.

**Figure S3.**
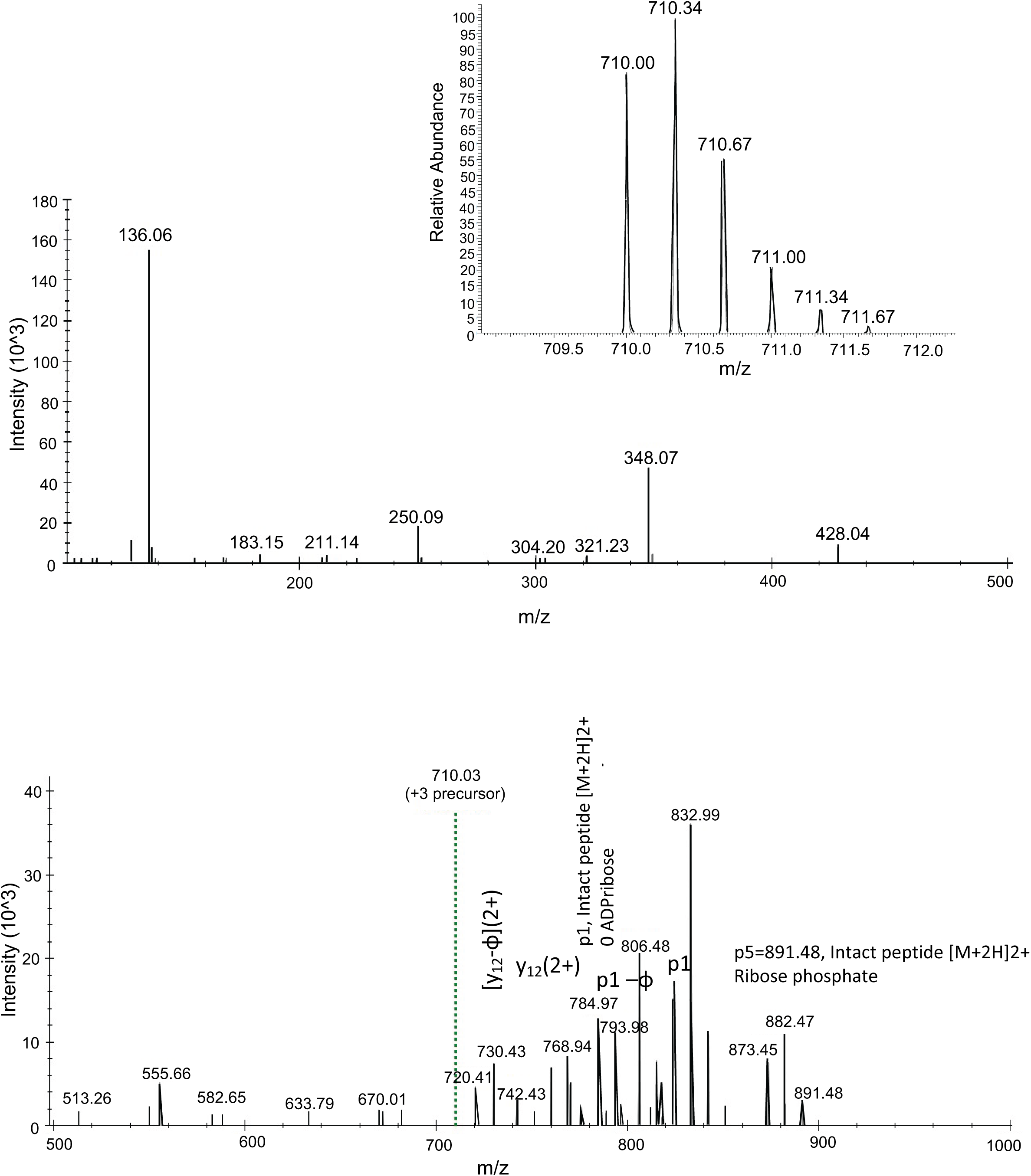
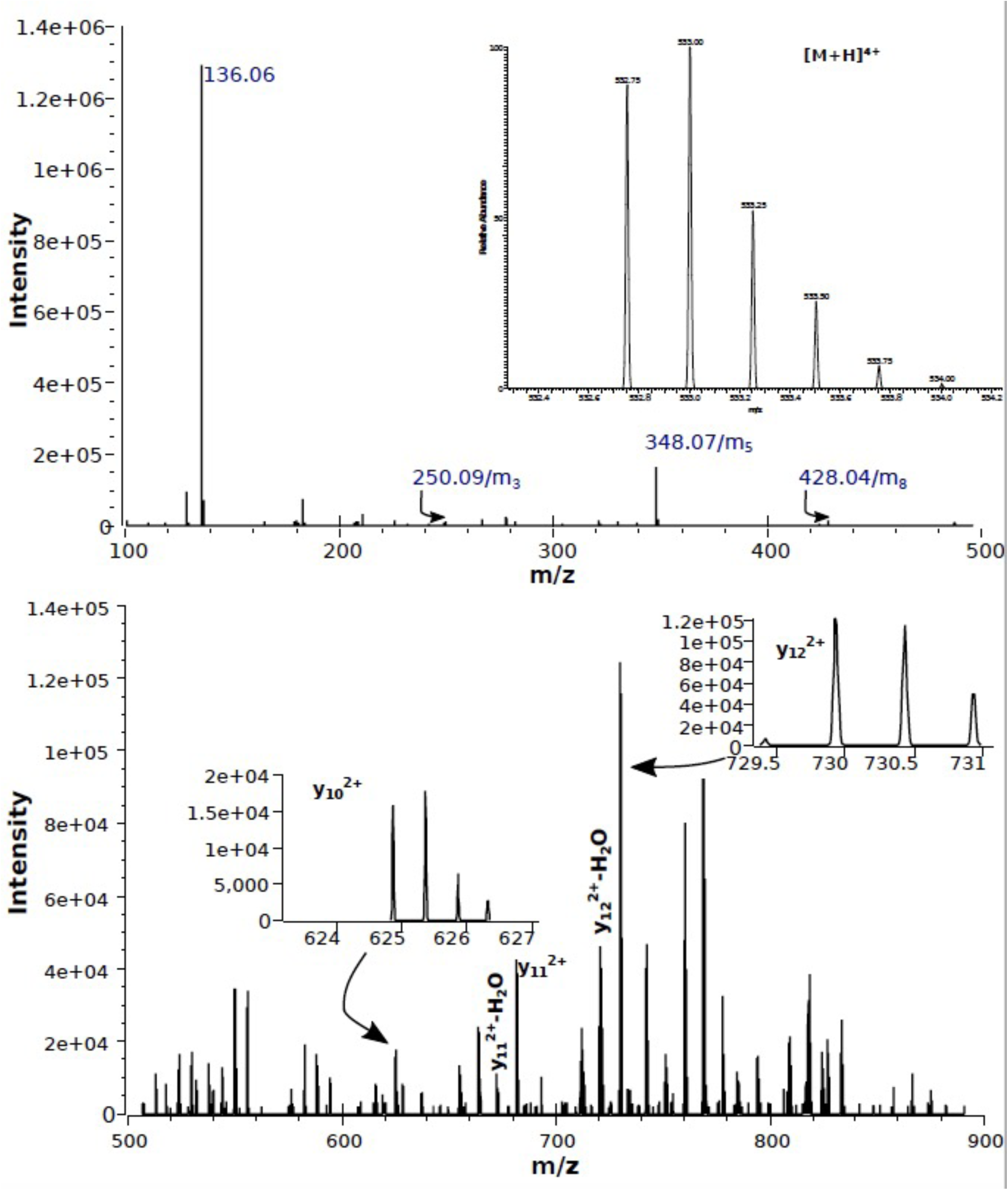
Characterization of ADP-ribosylation of SET8 synthetic peptide KPLAGIYRKREEK-NH2 using high resolution/high accuracy MS/MS HCD fragmentation, [M+3H]3+ precursor m/z 710.00 and [M+4H]4+ precursor m/z 532.75. (A) Characteristics ADP-ribose modification fragmentation ions (m-ions peptide free modification fragment ions):m1= 136.1, m3=250, m5=348, m8=428 (top panel). Ion m/z 136.1 could be sourced from either tyrosine immonium ion (m/z 136.0762) or [adenine+H]+ (m/z 136.0623). Fragment ions with intact peptide p1 and intact peptide with ribose phosphate p5 are shown on the bottom panel, m/z 500 - 900. The nomenclature for ADP-ribosylation fragment ions is according to (Hengel et al, 2009). Inset shows the [M+3H]3+ precursor full scan MS spectrum. (B) Similar to charge state +3, fragmented precursor charge state +4 showed characteristics ADP-ribose modification fragment ions. Increased intensity sequencing ions are present in the bottom panel, m/z 500 -900. Inset shows the [M+4H]4+ precursor full scan MS spectrum.

**Figure S4.**
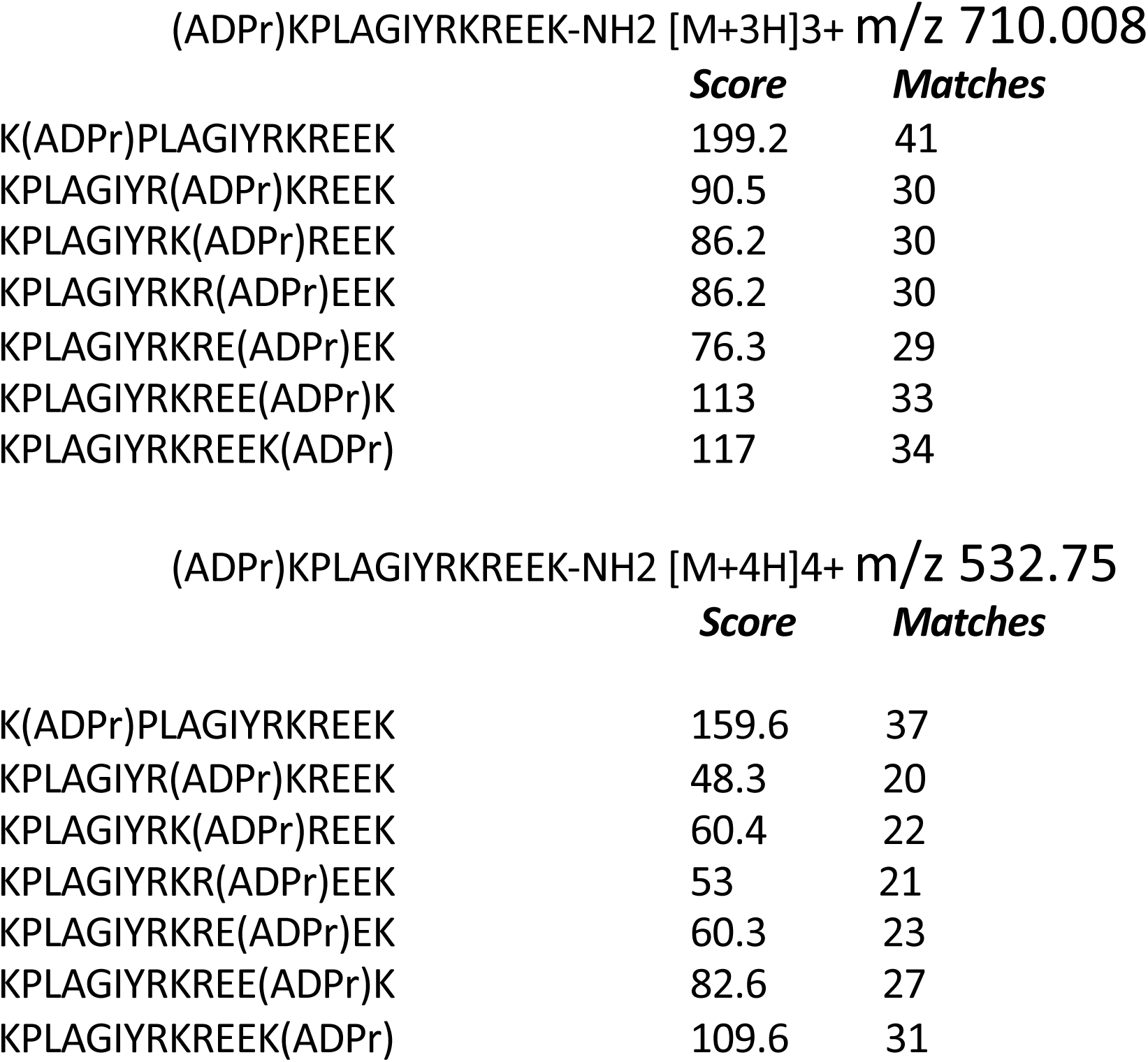
Characterization of ADP-ribosylation of SET8 synthetic peptide KPLAGIYRKREEK-NH2 using MS/MS CID fragmentation. [M+3H]3+ precursor m/z 710.008 and [M+4H]4+ precursor m/z 532.75 are shown. Scoring of ion propensity matches of the ADPr unit relative to seven possible amino acid locations in CID MS/MS spectra of (ADPr)KPLAGIYRKREEK-NH2.

**Figure S5.**
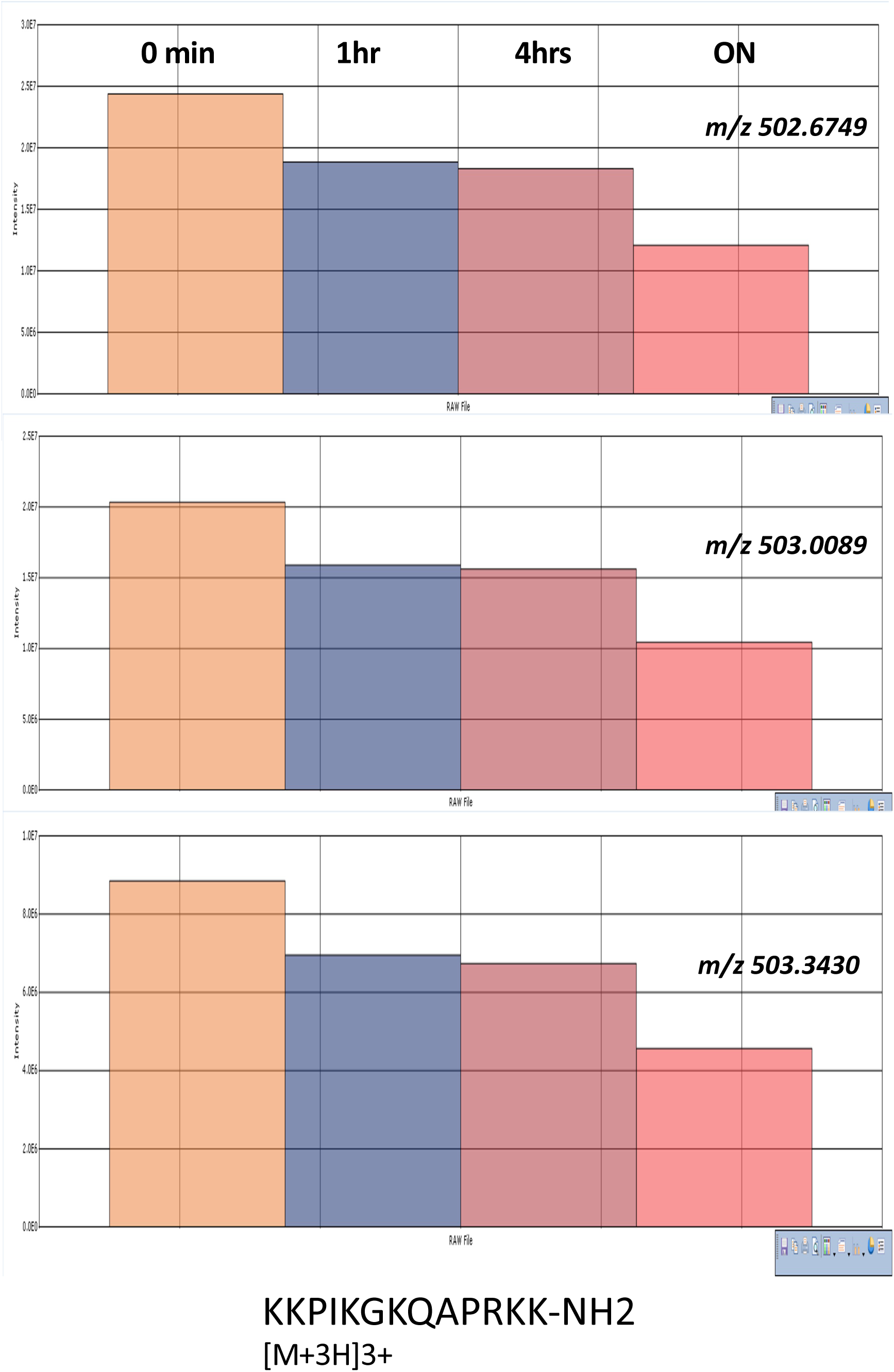
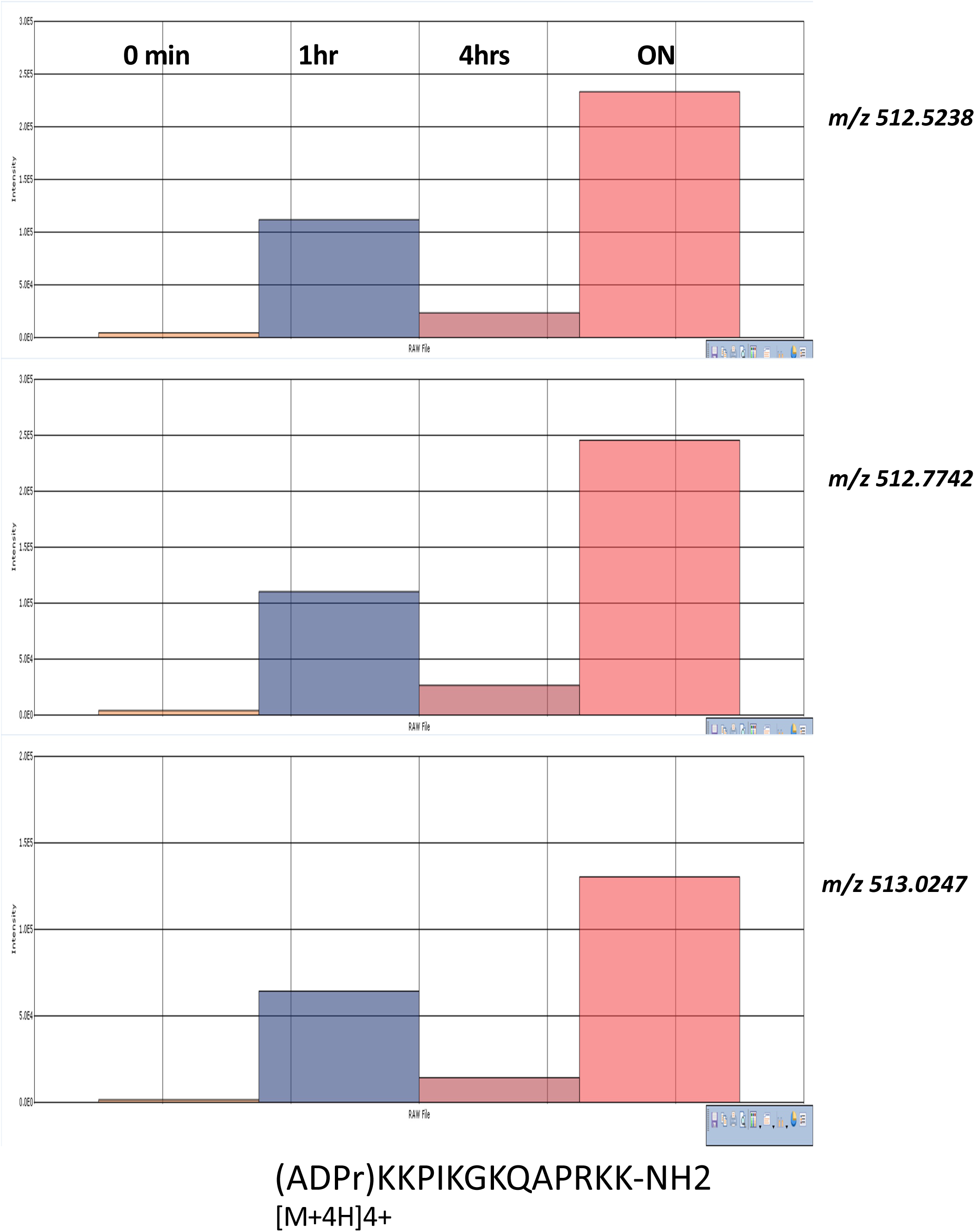
Characterization of ADP-ribosylation of SET8 synthetic peptide KKPIKGKQAPRKK-NH2 using high resolution mass spectrometry and isotope quantification in nanoLC. (A) Quantification of peptide KKPIKGKQAPRKK-NH2 (charge +3) at four time points: 0 min (negative control), 1hr, 4 hrs and overnight (ON) reaction with PARP1. Peak areas were determined with SIEVE software package for the three main isotopes m/z 502.67534, m/z 503.00960 and m/z 503.34383, theoretical values determined with Protein Prospector MS Isotope. The figure displays the measured isotopes 0, 1 and 2 with Orbitrap MS. (B) Quantification of peptide (ADPr)KKPIKGKQAPRKK-NH2 (charge +4) at four time points: 0 min (negative control), 1hr, 4 hrs and overnight (ON) reaction with PARP1. Peak areas were determined with SIEVE software package for the three main isotopes m/z 512.52360, m/z 512.77430 and m/z 513.02497, theoretical values determined with Protein Prospector MS Isotope. The figure displays the measured isotopes 0, 1 and 2 with Orbitrap MS.

**Figure S6.**
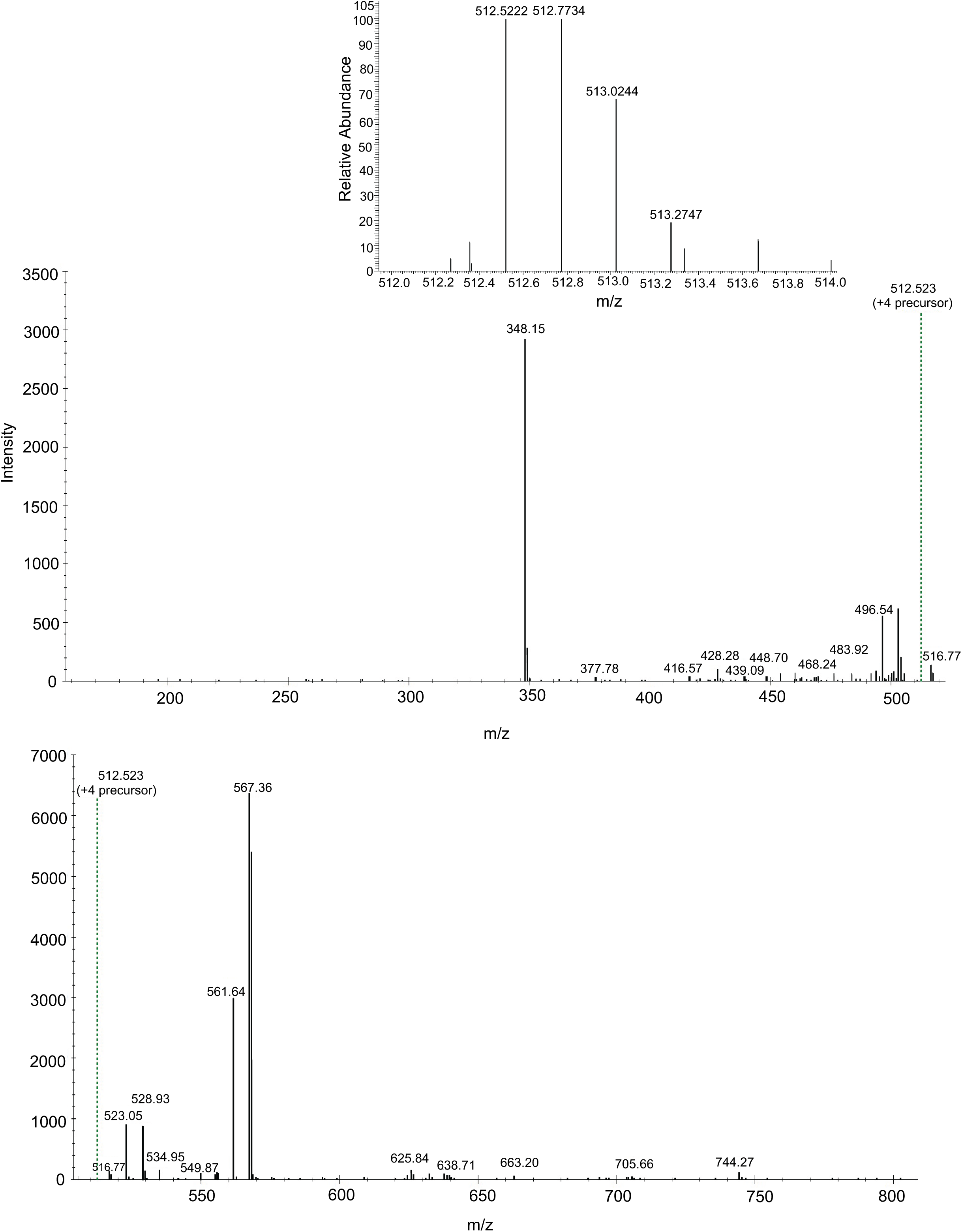
Characterization of ADP-ribosylation of SET8 synthetic peptide KKPIKGKQAPRKK-NH2 using MS/MS CID fragmentation, [M+4H]4+ precursor m/z 512.52. Characteristics ADP-ribose modification fragmentation ions (m-ions peptide free modification fragment ions): m5=348 and m8=428 (top panel). Fragment peptide ions are present most probably due to neutral loss of ADP-ribose fragments, within the unit resolution on linear ion trap MS/MS spectra (bottom panel). Inset shows the [M+4H]4+ precursor full scan MS spectrum.

**Figure S7.**
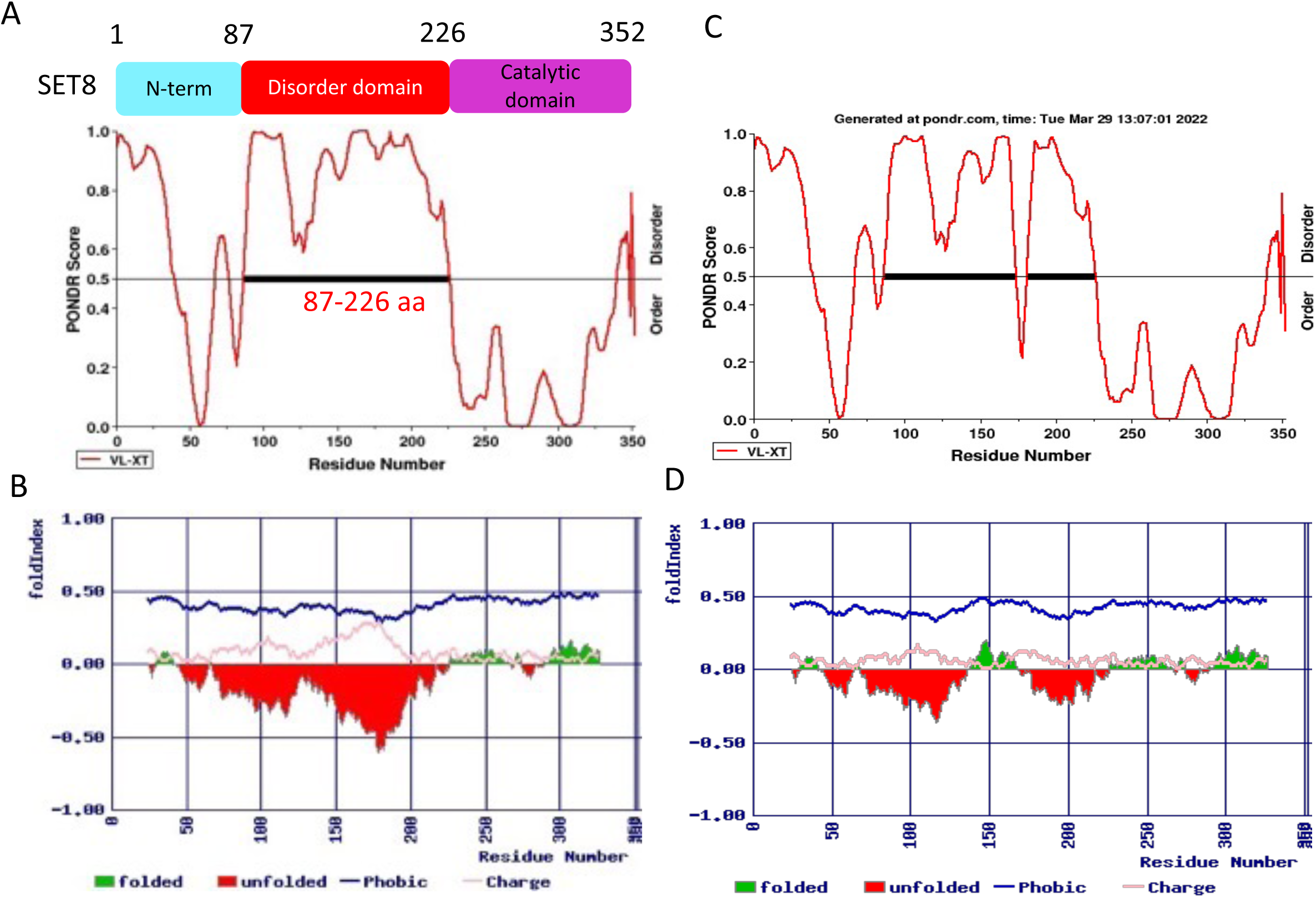
Folding Profile of Wild Type SET8 and SET8 mutation. (A) & (C) Structural disordered prediction for SET8 (87-226 aa disordered region) & SET8 mutation using PONDR-VLXT. (B) & (D) Protein folding profiling using FoldIndex tool for SET8 & SET8 mutation (greens are folded and reds are unfolded).

**Figure S8.**
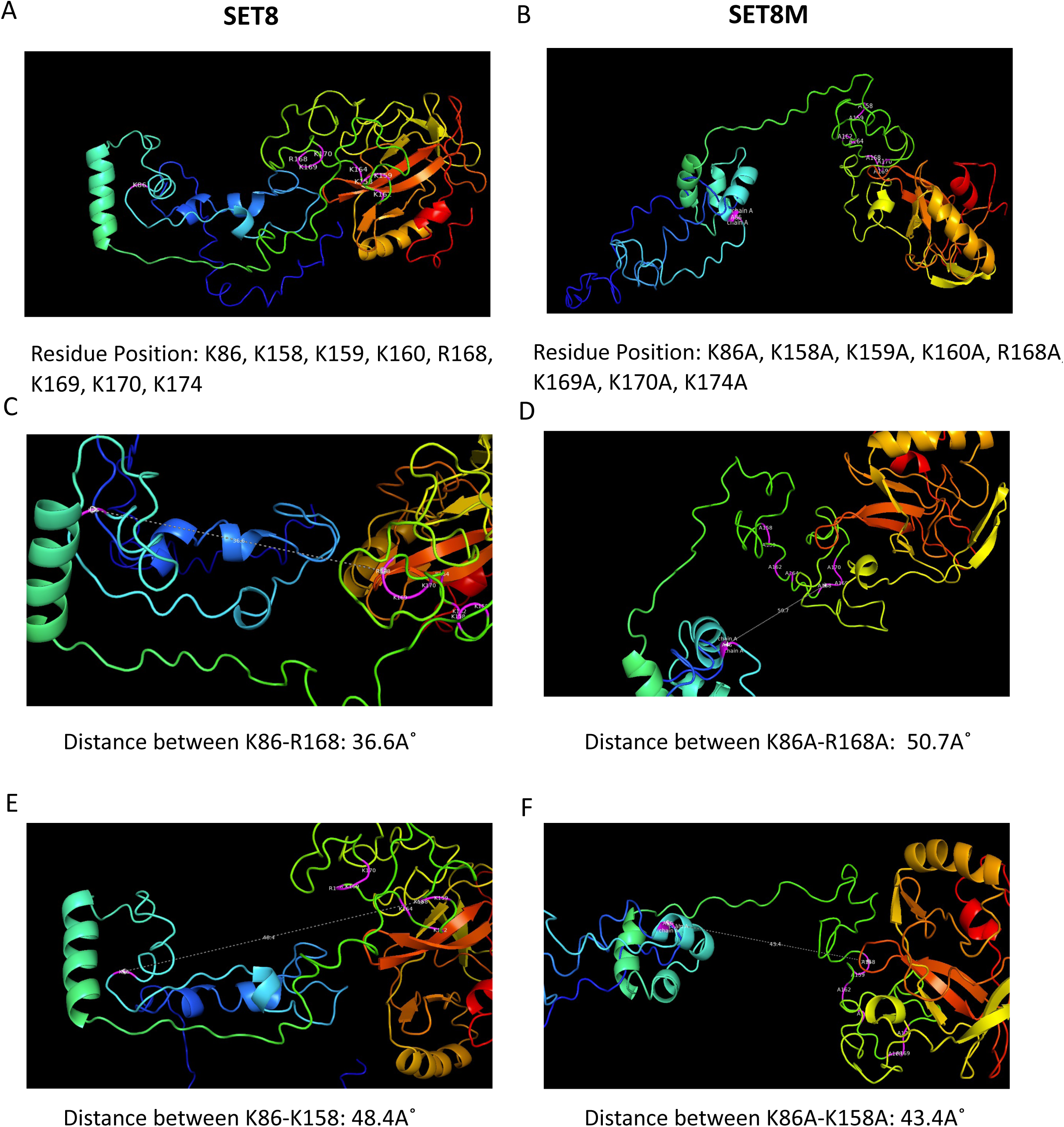
PDB structures of SET8 and mutated SET8 generated from I-TASSER. (A) PDB structure of Wild Type SET8 protein where lysine enriched domains are marked in pink (K86, K158, K159, K160, K162, R168, K169, K170, K174). (B) PDB structure of Mutated SET8 protein where mutated alanine enriched domains are colored in pink. (C) Distance from K86-R168 = 36.6Å of wild type SET8. (D) Distance from K86A-R168A = 50.7Å of mutated SET8. (E) Distance from K86-K158 = 48.4Å of wild type SET8. (F) Distance from K86A-K158A = 43.4Å of mutated SET8.

**Figure S9.**
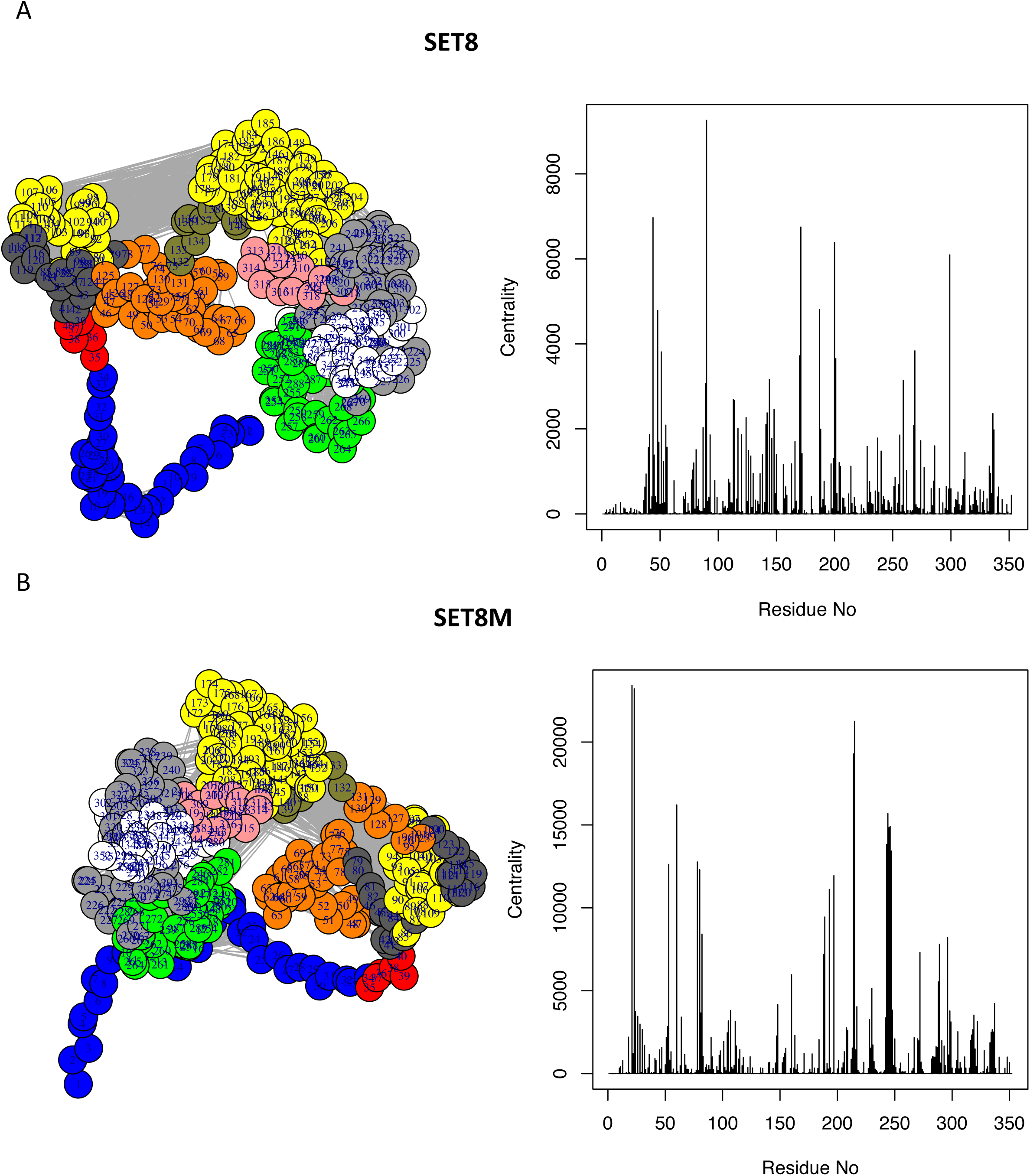
Structure Network of the Monomeric Structure of the wildtype and mutated SET8. (A) Structture Network Model(top, right) and Betweenness Centrality Plotting (top, left) throughout residue space for wildtype SET8. (B) Structture Network Model (bottom, right) and Betweenness Centrality Plotting (bottom, left) throughout residue space for mutated SET8.

**Figure S10.**
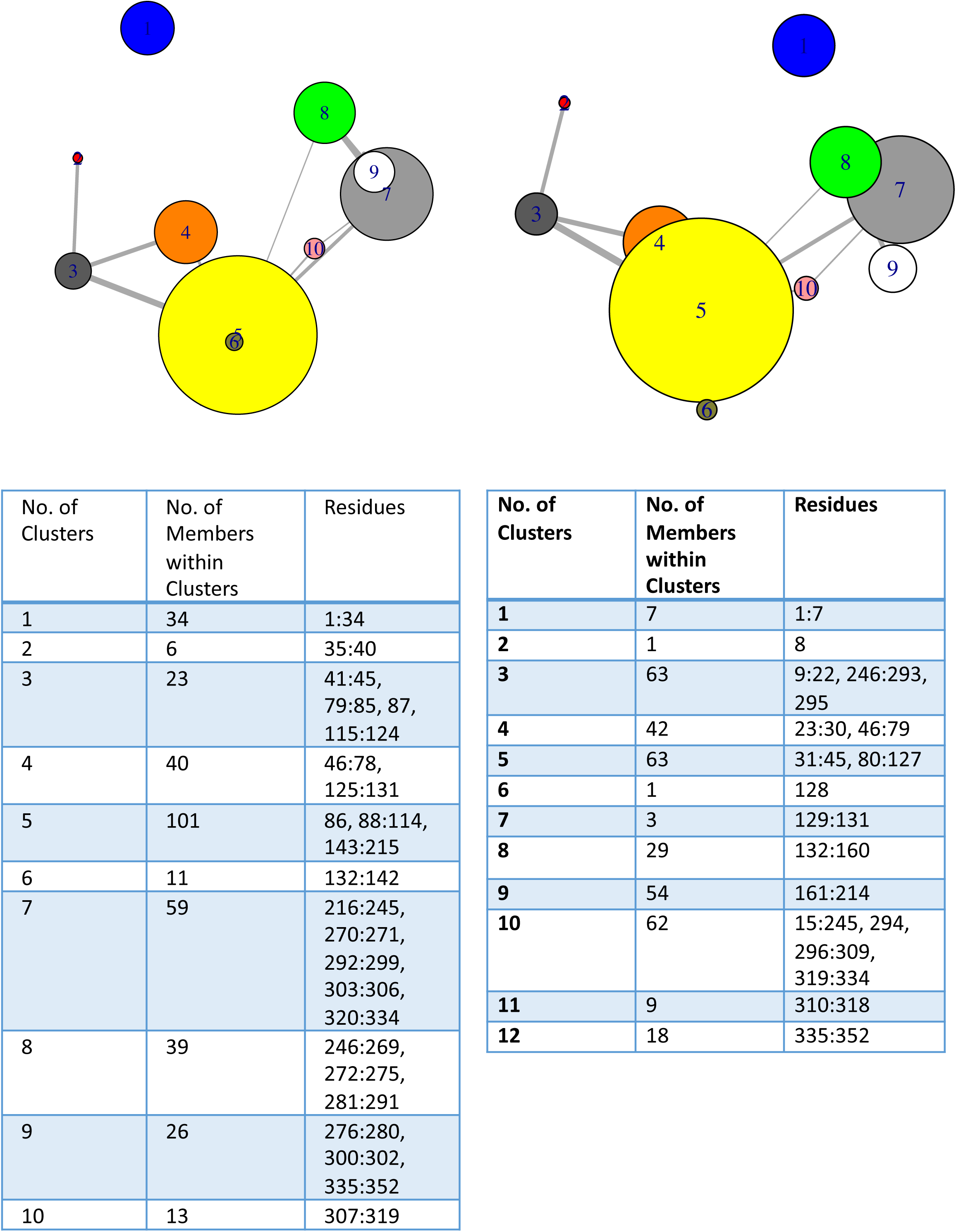
Cluster Network Models based on Betweenness Centrality Scoring where wild type SET8 has 10 modules (top, right), mutated SET8 has 12 modules (top, left). The residual members of each clusters are given in Tabular form for wild type SET8 (bottom, right) and mutated SET8 (bottom, left).

**Figure S11.**
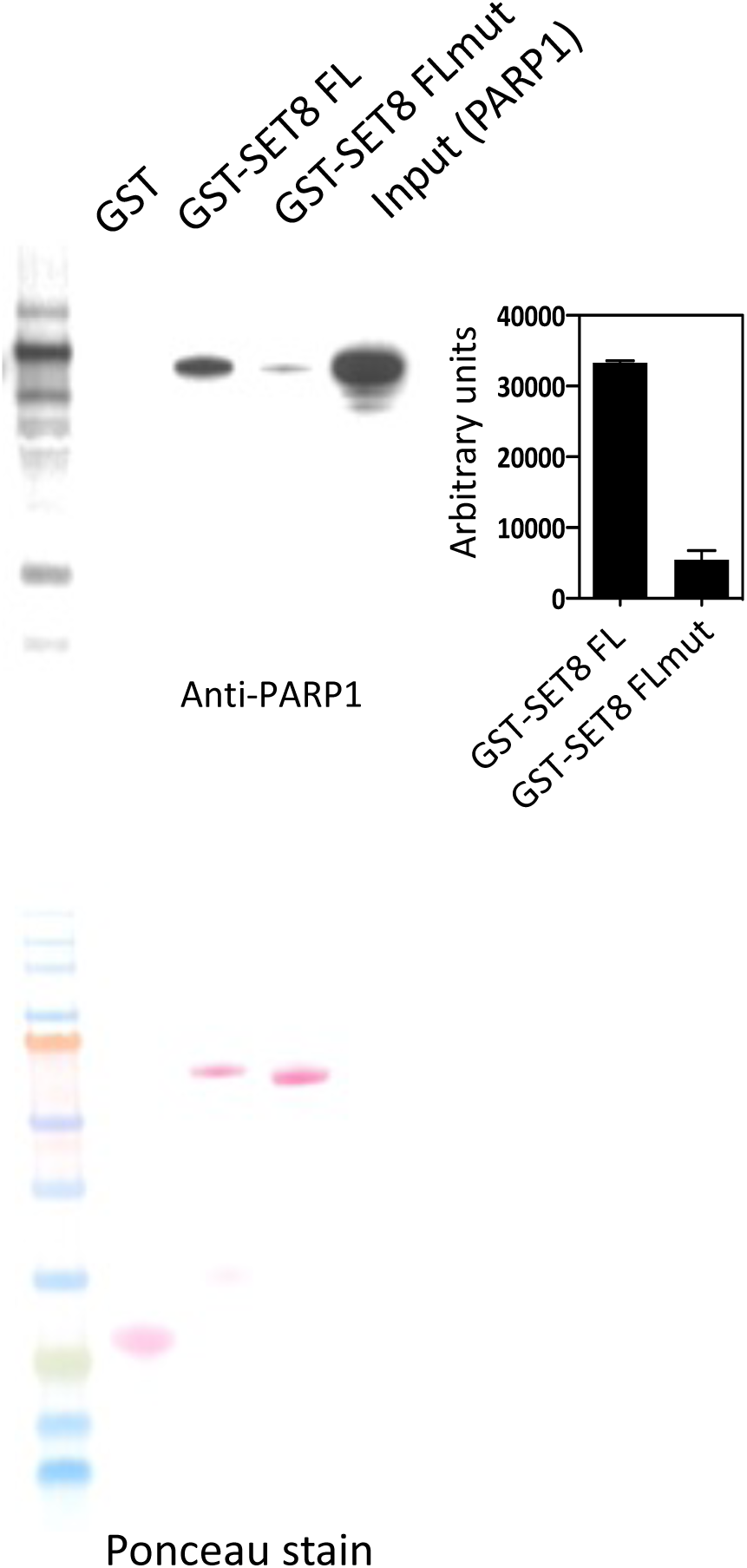
SET8 Lysine mutation restrict PARP1 binding. GST pull down of GST-SET8FL or GST-SET8 M (K86A, K158/159A, K162/164A, R168A, K169/170A, K174A) of PARP1 full-length enzyme. Western blot probed with anti-PARP1 antibody (top, left), and the relative pulldown of PARP1 quantitated and shown in arbitrary unit (top, right). Loading of the exact blot shown using ponceau staining (bottom).

**Figure S12:**
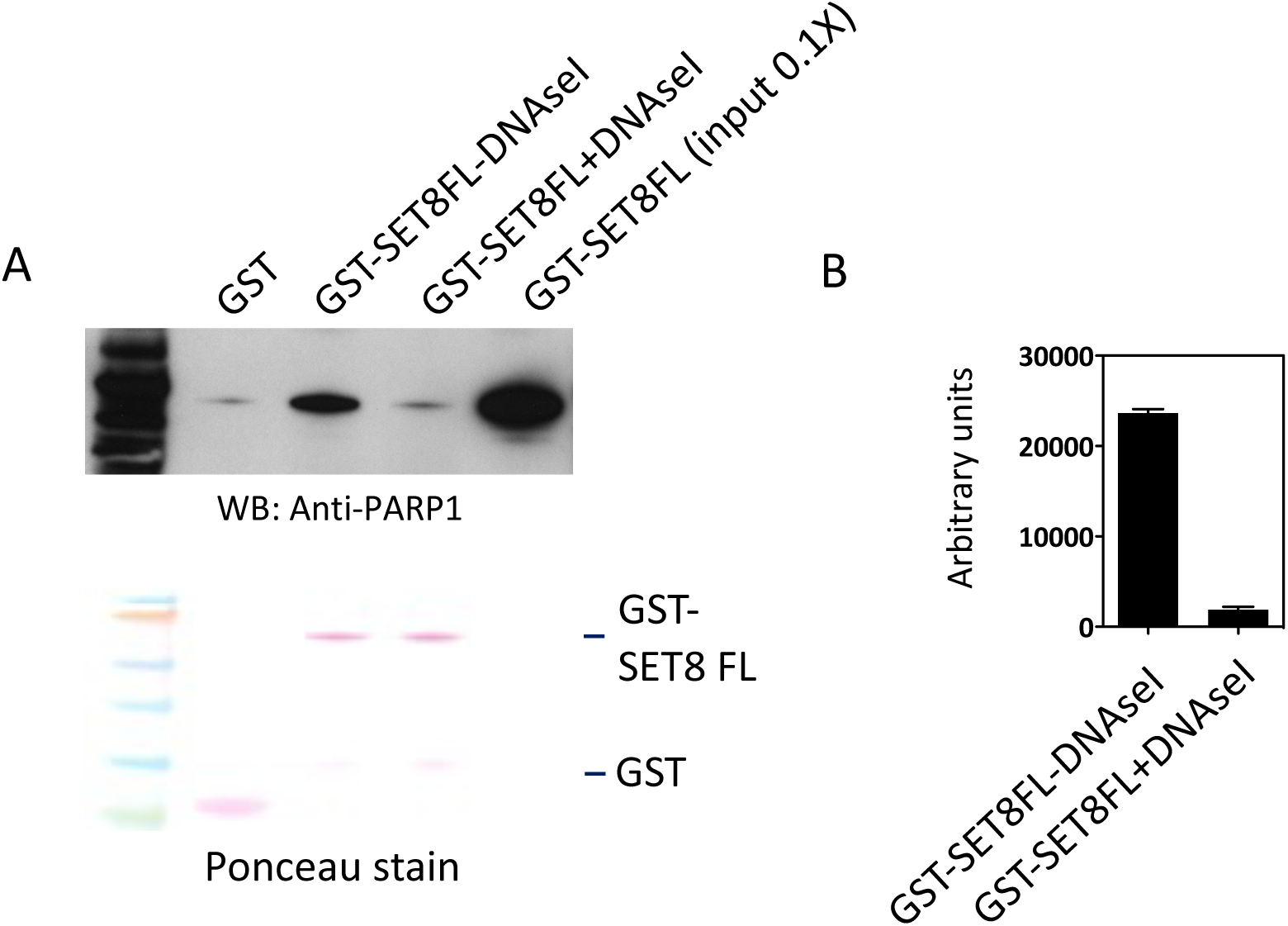
DNA enhances SET8 and PARP1 binding. (A) GST pull down assay of GST-SET8 FL with PARP1 full length in the presence or absence of DNase I. Western blot using anti-PARP1 antibody revels the presence of PARP1 (top, left side). (B) The relative pulldown of PARP1 quantitated and shown in arbitrary unit (right side). Loading of the exact blot shown using ponceau staining (bottom, left side).

**Figure S13:**
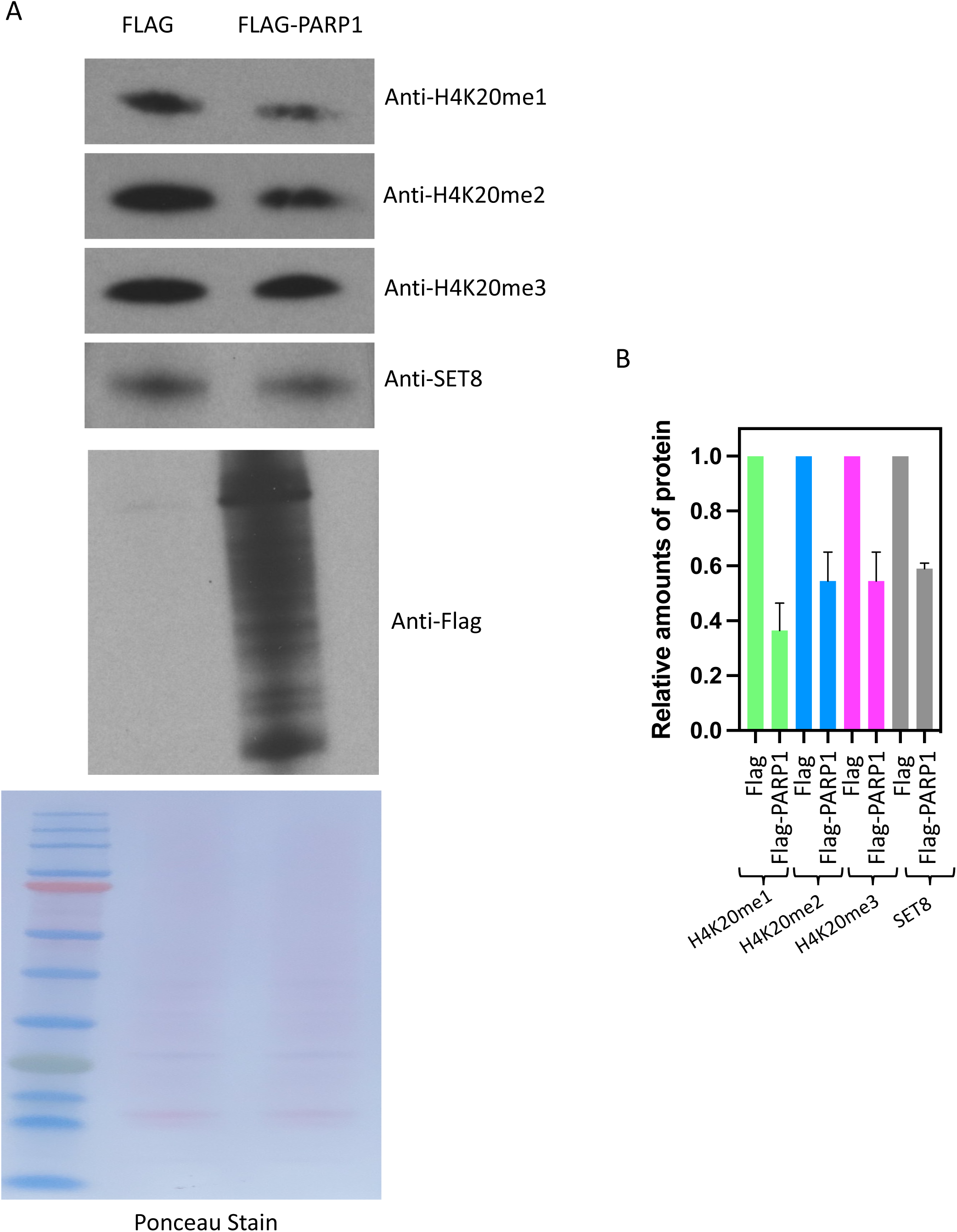
PARP1 overexpression leads to change the cellular levels of H4K20 methylation. (A) Equal amounts of cell extract from cells transfected with FLAG or FLAG-PARP1, blotted and probed with antibodies as indicated. Equal loading is shown by ponceau staining. (B) Western blot showing levels of mono, di, tri methylation of H4K20 and SET8. Relative amounts of protein are measured by scanning three independent blots

**Figure S14:**
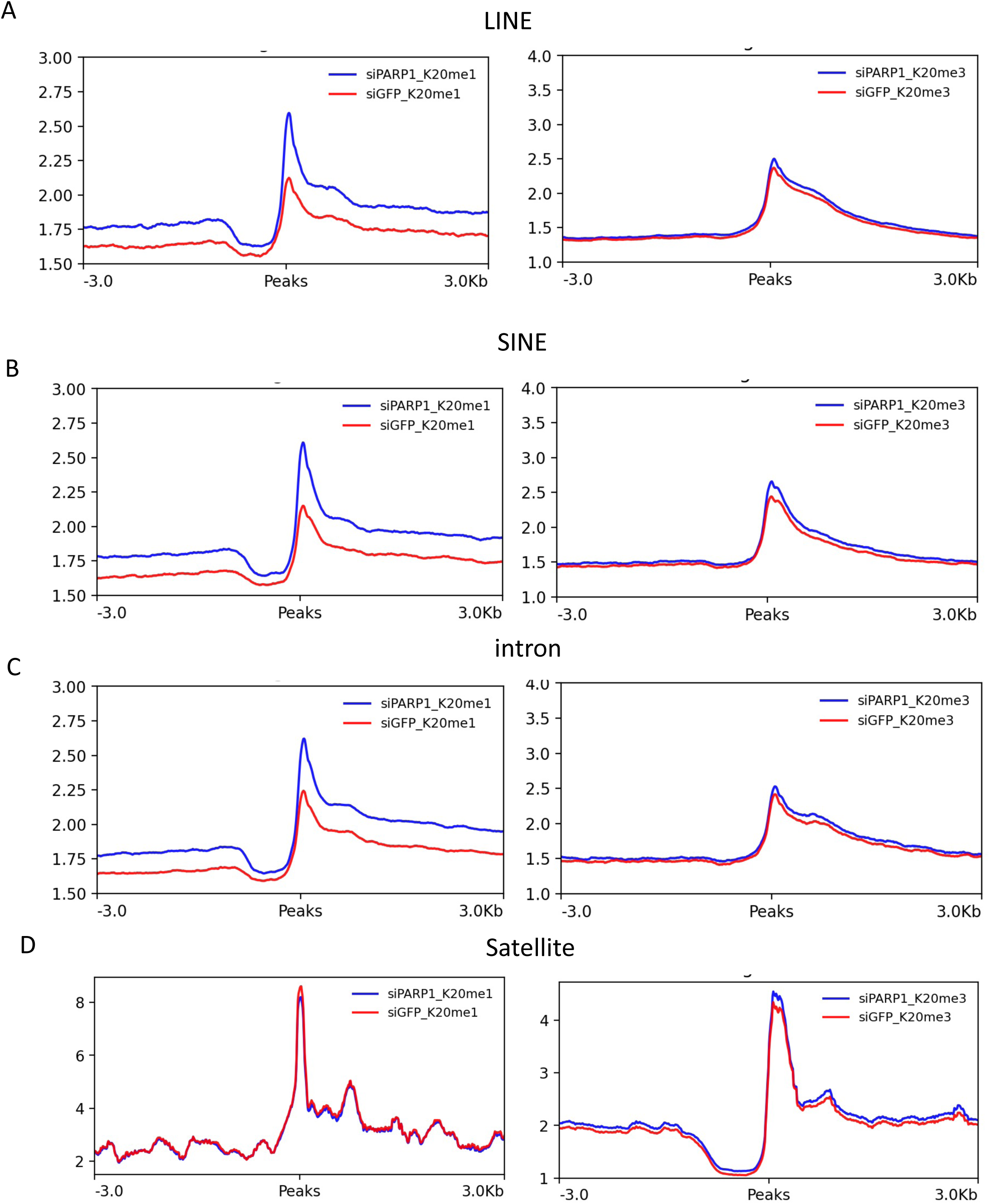
Enrichment Profiling for H4K20me1 (right) and H4K20me3 (left) for PARP1 Knockdown and Control Conditions for different Peak Annotating genomic Regions. (A) LINE, (B) SINE, (C) Intron, and (D) Satellite.

**Figure S15:**
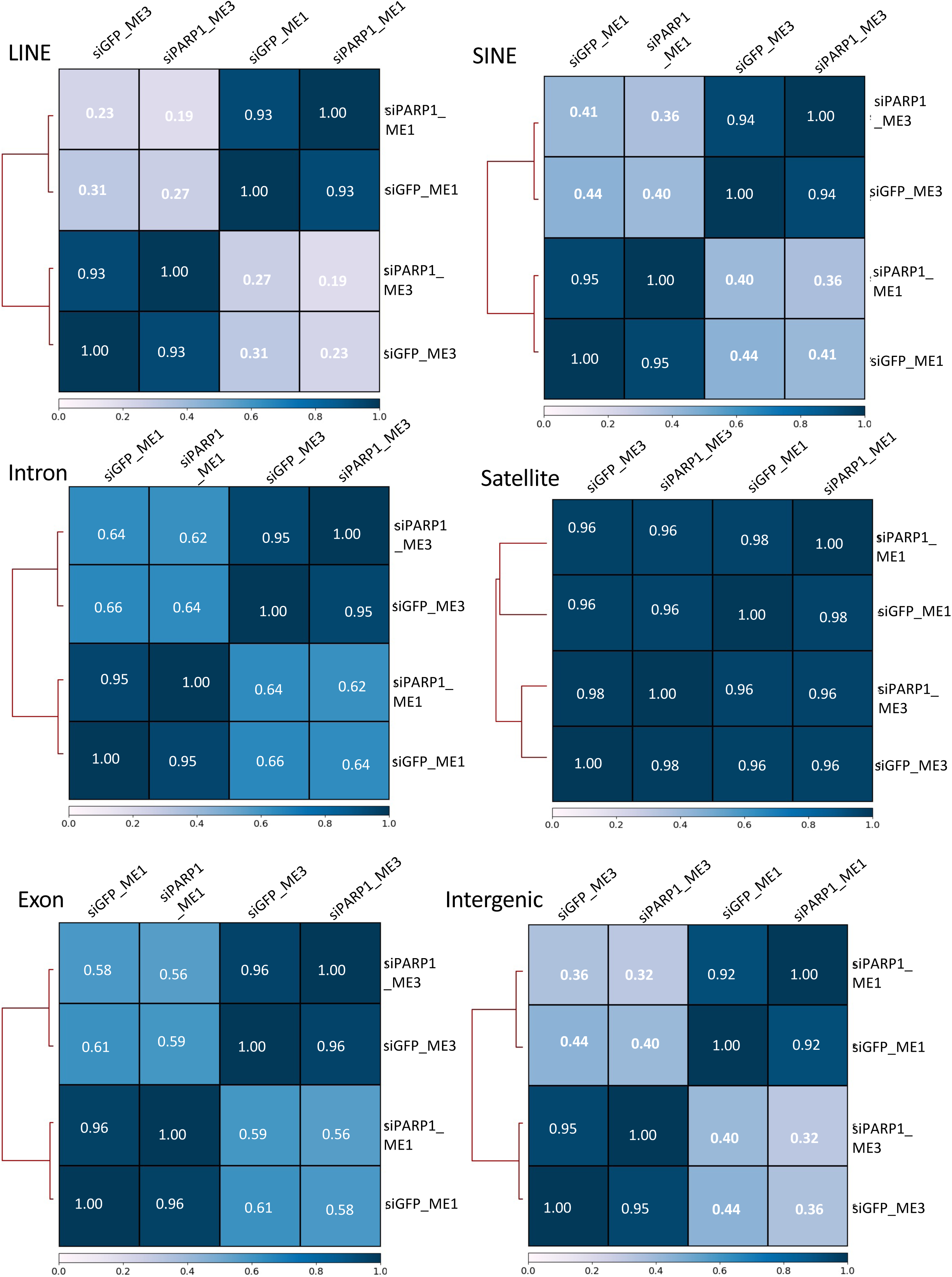
Spearman Correlation Heatmaps of PARP1 Knockdown and Control Conditions for different Peak Annotating Regions of H4K20me1 and H4K20me3 i.e., LINE, SINE, Intron, Satellite, Exon, and Intergenic.

**Figure S16:**
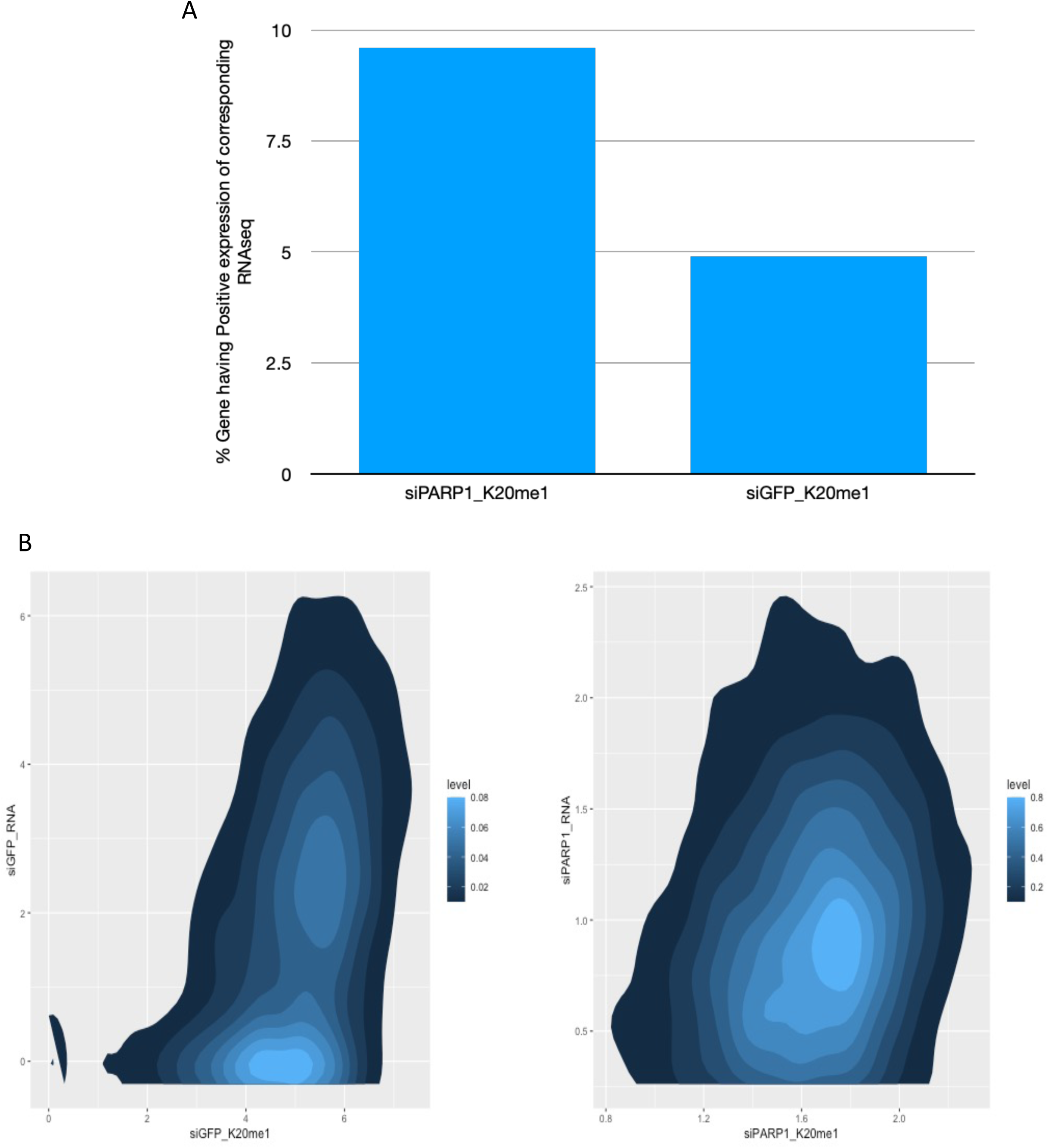
Comparative study between RNA seq and CHIP seq from PARP1 Knockdown and Control Conditions. (A) Overlaping genes between intragenic regions from different conditions and corresponding Positive Expression from RNA seq. (B) Density plot Profiling to show the distribution of the expression between siGFP_H4K20me1 positive gene expression and corresponding RNA seq genomic expression profiling (top, left) and, between siPARP1_H4K20me1 positive gene expression and corresponding RNA seq genomic expression profiling (top, right). (C) Density plot profiling to show the distribution of the expression between siGFP_H4K20me3 positive gene expression and corresponding RNA seq genomic expression profiling (bottom, left) and, between siPARP1_H4K20me3 positive gene expression and corresponding RNA seq genomic expression profiling (bottom, right).

**Figure S17:**
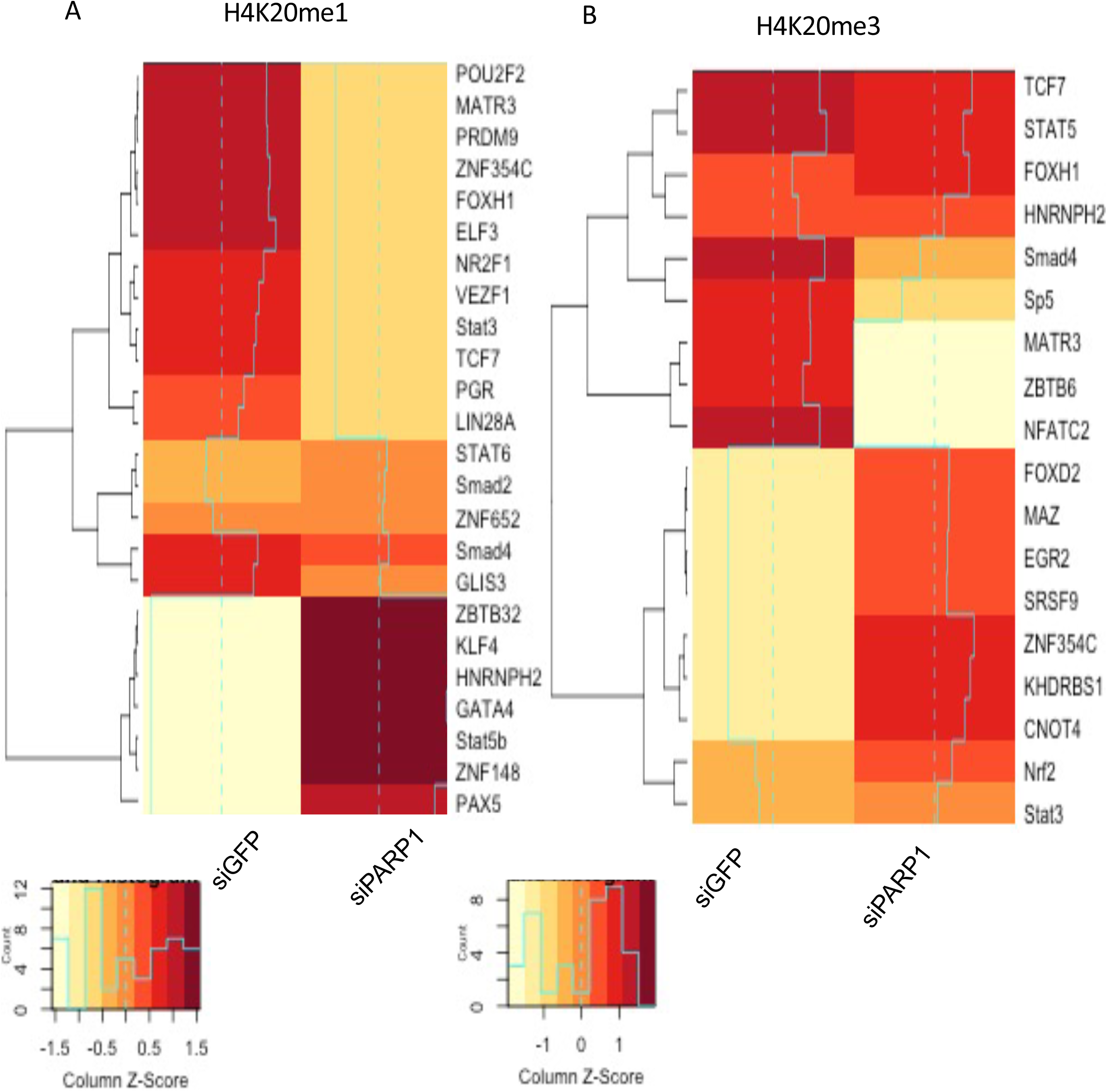
Transcription factor binding site analysis. (A) Heatmap representing the enrichment of consensus TF-binding motifs identified from H4K20me1 ChIP-seq in PARP1 knockdown cells and its control. (B) Heatmap representing the enrichment of consensus TF-binding motifs identified from H4K20me3 ChIP-seq in PARP1 knockdown cells and its control.

**Figure S18:**
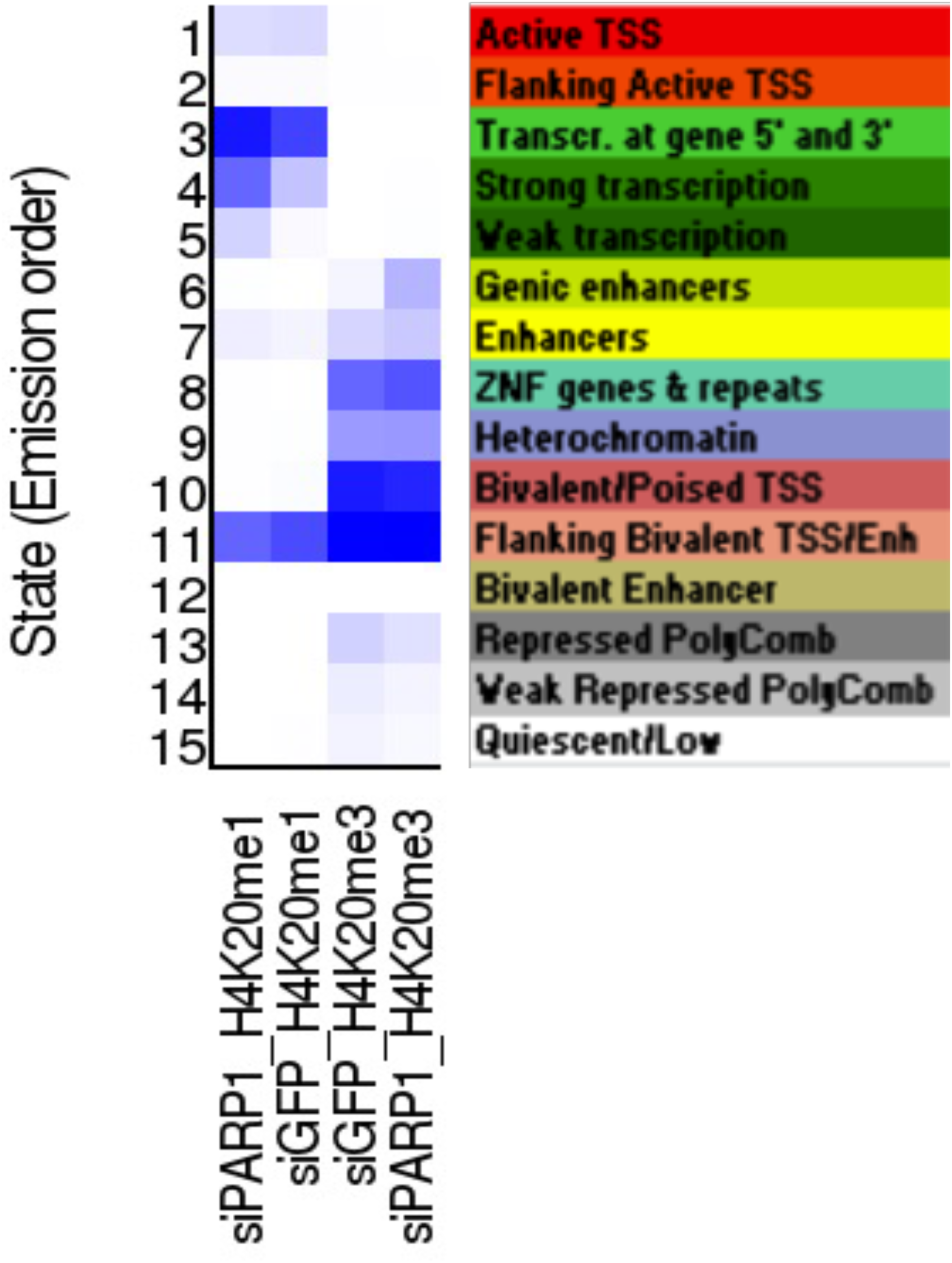
Study of Chromatin States for H4K20me1 and H4K20me3 at PARP1 Knockdown and Control Conditions using CHROMHMM.

Table S1: Proteomic analysis of SET8 pull-down in HEK293T cells using LC-MS.

Table S2: Residual distance Between residues at Wild Type and Mutated state for SET8

Table S3: RNA seq of genes post PARP1 knock down

Table S4A: Pathways associated TFs enriched due to control condition.

Table S4B: Pathways associated TFs enriched due to PARP1 knock down

## References

1. Ma,Y., Kanakousaki,K. and Buttitta,L. (2015) How the cell cycle impacts chromatin architecture and influences cell fate. Front. Genet., 10.3389/fgene.2015.00019.

2. Ho,L. and Crabtree,G.R. (2010) Chromatin remodelling during development. Nature, 10.1038/nature08911.

3. Perino,M. and Veenstra,G.J.C. (2016) Chromatin Control of Developmental Dynamics and Plasticity. Dev. Cell, 10.1016/j.devcel.2016.08.004.

4. Beagrie,R.A. and Pombo,A.N.A. (2017) Continuous chromatin changes. Nature.

5. Ma,Y. and Buttitta,L. (2017) Chromatin organization changes during the establishment and maintenance of the postmitotic state. Epigenetics and Chromatin, 10.1186/s13072-017-0159-8.

6. Bannister,A.J. and Kouzarides,T. (2011) Regulation of chromatin by histone modifications. Cell Res., 10.1038/cr.2011.22.

7. Lieberman-Aiden,E., Van Berkum,N.L., Williams,L., Imakaev,M., Ragoczy,T., Telling,A., Amit,I., Lajoie,B.R., Sabo,P.J., Dorschner,M.O., et al. (2009) Comprehensive mapping of long-range interactions reveals folding principles of the human genome. Science (80-.)., 10.1126/science.1181369.

8. Kuo,M.H. and Allis,C.D. (1998) Roles of histone acetyltransferases and deacetylases in gene regulation. BioEssays, 10.1002/(SICI)1521-1878(199808)20:8<615::AID-BIES4>3.0.CO;2-H.

9. Grunstein,M. (1997) Histone acetylation in chromatin structure and transcription. Nature, 10.1038/38664.

10. Smith,B.C. and Denu,J.M. (2009) Chemical mechanisms of histone lysine and arginine modifications. Biochim. Biophys. Acta - Gene Regul. Mech., 10.1016/j.bbagrm.2008.06.005.

11. Herz,H.M., Garruss,A. and Shilatifard,A. (2013) SET for life: Biochemical activities and biological functions of SET domain-containing proteins. Trends Biochem. Sci., 10.1016/j.tibs.2013.09.004.

12. Oda,H., Okamoto,I., Murphy,N., Chu,J., Price,S.M., Shen,M.M., Torres-Padilla,M.E., Heard,E. and Reinberg,D. (2009) Monomethylation of Histone H4-Lysine 20 Is Involved in Chromosome Structure and Stability and Is Essential for Mouse Development. Mol. Cell. Biol., 10.1128/mcb.01768-08.

13. Couture,J.F., Collazo,E., Brunzelle,J.S. and Trievel,R.C. (2005) Structural and functional analysis of SET8, a histone H4 Lys-20 methyltransferase. Genes Dev., 10.1101/gad.1318405.

14. Schotta,G., Sengupta,R., Kubicek,S., Malin,S., Kauer,M., Callén,E., Celeste,A., Pagani,M., Opravil,S., De La Rosa-Velazquez,I.A., et al. (2008) A chromatin-wide transition to H4K20 monomethylation impairs genome integrity and programmed DNA rearrangements in the mouse. Genes Dev., 10.1101/gad.476008.

15. Kleine-Kohlbrecher,D., Christensen,J., Vandamme,J., Abarrategui,I., Bak,M., Tommerup,N., Shi,X., Gozani,O., Rappsilber,J., Salcini,A.E., et al. (2010) A Functional Link between the Histone Demethylase PHF8 and the Transcription Factor ZNF711 in X-Linked Mental Retardation. Mol. Cell, 10.1016/j.molcel.2010.03.002.

16. Cao,X., Chen,Y., Wu,B., Wang,X., Xue,H., Yu,L., Li,J., Wang,Y., Wang,W., Xu,Q., et al. (2020) Histone H4K20 Demethylation by Two hHR23 Proteins. Cell Rep., 10.1016/j.celrep.2020.03.001.

17. Karachentsev,D., Sarma,K., Reinberg,D. and Steward,R. (2005) PR-Set7-dependent methylation of histone H4 Lys 20 functions in repression of gene expression and is essential for mitosis. Genes Dev., 10.1101/gad.1263005.

18. Shoaib,M., Walter,D., Gillespie,P.J., Izard,F., Fahrenkrog,B., Lleres,D., Lerdrup,M., Johansen,J.V., Hansen,K., Julien,E., et al. (2018) Histone H4K20 methylation mediated chromatin compaction threshold ensures genome integrity by limiting DNA replication licensing. Nat. Commun., 10.1038/s41467-018-06066-8.

19. Abbas,T., Shibata,E., Park,J., Jha,S., Karnani,N. and Dutta,A. (2010) CRL4Cdt2 regulates cell proliferation and histone gene expression by targeting PR-Set7/Set8 for degradation. Mol. Cell, 10.1016/j.molcel.2010.09.014.

20. Centore,R.C., Havens,C.G., Manning,A.L., Li,J.M., Flynn,R.L., Tse,A., Jin,J., Dyson,N.J., Walter,J.C. and Zou,L. (2010) CRL4Cdt2-mediated destruction of the histone methyltransferase Set8 prevents premature chromatin compaction in S phase. Mol. Cell, 10.1016/j.molcel.2010.09.015.

21. Oda,H., Hübner,M.R., Beck,D.B., Vermeulen,M., Hurwitz,J., Spector,D.L. and Reinberg,D. (2010) Regulation of the Histone H4 Monomethylase PR-Set7 by CRL4Cdt2-Mediated PCNA-Dependent Degradation during DNA Damage. Mol. Cell, 10.1016/j.molcel.2010.10.011.

22. Yin,L., Yu,V.C., Zhu,G. and Chang,D.C. (2008) SET8 plays a role in controlling G1/S transition by blocking lysine acetylation in histone through binding to H4 N-terminal tail. Cell Cycle, 10.4161/cc.7.10.5867.

23. Frescas,D. and Pagano,M. (2008) Deregulated proteolysis by the F-box proteins SKP2 and β-TrCP: Tipping the scales of cancer. Nat. Rev. Cancer, 10.1038/nrc2396.

24. Dephoure,N., Zhou,C., Villén,J., Beausoleil,S.A., Bakalarski,C.E., Elledge,S.J. and Gygi,S.P. (2008) A quantitative atlas of mitotic phosphorylation. Proc. Natl. Acad. Sci. U. S. A., 10.1073/pnas.0805139105.

25. Huttlin,E.L., Jedrychowski,M.P., Elias,J.E., Goswami,T., Rad,R., Beausoleil,S.A., Villén,J., Haas,W., Sowa,M.E. and Gygi,S.P. (2010) A tissue-specific atlas of mouse protein phosphorylation and expression. Cell, 10.1016/j.cell.2010.12.001.

26. Olsen,J. V, Vermeulen,M., Santamaria,A., Kumar,C., Miller,M.L., Jensen,L.J., Gnad,F., Cox,J., Jensen,T.S., Nigg,E.A., et al. (2010) Quantitative Phosphoproteomics Reveals Widespread Full Phosphorylation Site Occupancy During Mitosis -- Olsen et al. 3 (104): ra3 -- Science Signaling (Supplemental). Sci. Signal.

27. Wu,S., Wang,W., Kong,X., Congdon,L.M., Yokomori,K., Kirschner,M.W. and Rice,J.C. (2010) Dynamic regulation of the PR-Set7 histone methyltransferase is required for normal cell cycle progression. Genes Dev., 10.1101/gad.1984210.

28. Alemasova,E.E. and Lavrik,O.I. (2019) Poly(ADP-ribosyl)ation by PARP1: Reaction mechanism and regulatory proteins. Nucleic Acids Res., 10.1093/nar/gkz120.

29. Altmeyer,M., Messner,S., Hassa,P.O., Fey,M. and Hottiger,M.O. (2009) Molecular mechanism of poly(ADP-ribosyl)ation by PARP1 and identification of lysine residues as ADP-ribose acceptor sites. Nucleic Acids Res., 10.1093/nar/gkp229.

30. Teloni,F. and Altmeyer,M. (2016) Survey and summary readers of poly(ADP-ribose): Designed to be fit for purpose. Nucleic Acids Res., 10.1093/nar/gkv1383.

31. Daniels,C.M., Ong,S.E. and Leung,A.K.L. (2015) The Promise of Proteomics for the Study of ADP-Ribosylation. Mol. Cell, 10.1016/j.molcel.2015.06.012.

32. Hottiger,M.O. (2015) Nuclear ADP-ribosylation and its role in chromatin plasticity, cell differentiation, and epigenetics. Annu. Rev. Biochem., 10.1146/annurev-biochem-060614-034506.

33. Leslie Pedrioli,D.M., Leutert,M., Bilan,V., Nowak,K., Gunasekera,K., Ferrari,E., Imhof,R., Malmström,L. and Hottiger,M.O. (2018) Comprehensive ADP-ribosylome analysis identifies tyrosine as an ADP-ribose acceptor site. EMBO Rep., 10.15252/embr.201745310.

34. Larsen,S.C., Hendriks,I.A., Lyon,D., Jensen,L.J. and Nielsen,M.L. (2018) Systems-wide Analysis of Serine ADP-Ribosylation Reveals Widespread Occurrence and Site-Specific Overlap with Phosphorylation. Cell Rep., 10.1016/j.celrep.2018.07.083.

35. Zhang,Y., Wang,J., Ding,M. and Yu,Y. (2013) Site-specific characterization of the Asp-and Glu-ADP-ribosylated proteome. Nat. Methods, 10.1038/nmeth.2603.

36. Vivelo,C.A. and Leung,A.K.L. (2015) Proteomics approaches to identify mono-(ADP-ribosyl)ated and poly(ADP-ribosyl)ated proteins. Proteomics, 10.1002/pmic.201400217.

37. Estève,P.O., Hang,G.C., Smallwood,A., Feehery,G.R., Gangisetty,O., Karpf,A.R., Carey,M.F. and Pradhan,S. (2006) Direct interaction between DNMT1 and G9a coordinates DNA and histone methylation during replication. Genes Dev., 10.1101/gad.1463706.

38. Estève,P.O., Chin,H.G., Benner,J., Feehery,G.R., Samaranayake,M., Horwitz,G.A., Jacobsen,S.E. and Pradhan,S. (2009) Regulation of DNMT1 stability through SET7- mediated lysine methylation in mammalian cells. Proc. Natl. Acad. Sci. U. S. A., 10.1073/pnas.0810362106.

39. Chin,H.G., Esteve,P.O., Ruse,C., Lee,J., Schaus,S.E., Pradhan,S., Hansen,U. and de la Cruz,E.M. (2020) The microtubule-associated histone methyltransferase SET8, facilitated by transcription factor LSF, methylates α-tubulin. J. Biol. Chem., 10.1074/jbc.RA119.010951.

40. Kim,G. Do, Ni,J., Kelesoglu,N., Roberts,R.J. and Pradhan,S. (2002) Co-operation and communication between the human maintenance and de novo DNA (cytosine-5) methyltransferases. EMBO J., 10.1093/emboj/cdf401.

41. Patnaik,D., Hang,G.C., Estève,P.O., Benner,J., Jacobsen,S.E. and Pradhan,S. (2004) Substrate specificity and kinetic mechanism of mammalian G9a histone H3 methyltransferase. J. Biol. Chem., 10.1074/jbc.M409604200.

42. Eng,J.K., McCormack,A.L. and Yates,J.R. (1994) An approach to correlate tandem mass spectral data of peptides with amino acid sequences in a protein database. J. Am. Soc. Mass Spectrom., 10.1016/1044-0305(94)80016-2.

43. Ponnaluri,V.K.C., Zhang,G., Estève,P.O., Spracklin,G., Sian,S., Xu,S. yong, Benoukraf,T. and Pradhan,S. (2017) NicE-seq: High resolution open chromatin profiling. Genome Biol., 10.1186/s13059-017-1247-6.

44. Chin,H.G., Sun,Z., Vishnu,U.S., Hao,P., Cejas,P., Spracklin,G., Estève,P.O., Xu,S.Y., Long,H.W. and Pradhan,S. (2020) Universal NicE-seq for high-resolution accessible chromatin profiling for formaldehyde-fixed and FFPE tissues. Clin. Epigenetics, 10.1186/s13148-020-00921-6.

45. Langmead,B. and Salzberg,S.L. (2012) Fast gapped-read alignment with Bowtie 2 - Supplemental Information. Nat. Methods.

46. Li,H., Handsaker,B., Wysoker,A., Fennell,T., Ruan,J., Homer,N., Marth,G., Abecasis,G. and Durbin,R. (2009) The Sequence Alignment/Map format and SAMtools. Bioinformatics, 10.1093/bioinformatics/btp352.

47. Tarasov,A., Vilella,A.J., Cuppen,E., Nijman,I.J. and Prins,P. (2015) Sambamba: Fast processing of NGS alignment formats. Bioinformatics, 10.1093/bioinformatics/btv098.

48. Zhang,Y., Liu,T., Meyer,C.A., Eeckhoute,J., Johnson,D.S., Bernstein,B.E., Nussbaum,C., Myers,R.M., Brown,M., Li,W., et al. (2008) Model-based analysis of ChIP-Seq (MACS). Genome Biol., 10.1186/gb-2008-9-9-r137.

49. Ramírez,F., Ryan,D.P., Grüning,B., Bhardwaj,V., Kilpert,F., Richter,A.S., Heyne,S., Dündar,F. and Manke,T. (2016) deepTools2: a next generation web server for deep-sequencing data analysis. Nucleic Acids Res., 10.1093/nar/gkw257.

50. Heinz,S., Benner,C., Spann,N., Bertolino,E., Lin,Y.C., Laslo,P., Cheng,J.X., Murre,C., Singh,H. and Glass,C.K. (2010) Simple Combinations of Lineage-Determining Transcription Factors Prime cis-Regulatory Elements Required for Macrophage and B Cell Identities. Mol. Cell, 10.1016/j.molcel.2010.05.004.

51. R Core Team. (2020) A Language and Environment for Statistical Computing. Available online: http://www.r-project.org. *R Found. Stat. Comput*.

52. Zhang,J., Lee,D., Dhiman,V., Jiang,P., Xu,J., McGillivray,P., Yang,H., Liu,J., Meyerson,W., Clarke,D., et al. (2020) An integrative ENCODE resource for cancer genomics. Nat. Commun., 10.1038/s41467-020-14743-w.

53. Robinson,J.T., Thorvaldsdóttir,H., Winckler,W., Guttman,M., Lander,E.S., Getz,G. and Mesirov,J.P. (2011) Integrative genomics viewer. Nat. Biotechnol., 10.1038/nbt.1754.

54. Estève,P.O., Vishnu,U.S., Chin,H.G. and Pradhan,S. (2020) Visualization and Sequencing of Accessible Chromatin Reveals Cell Cycle and Post-HDAC inhibitor Treatment Dynamics. J. Mol. Biol., 10.1016/j.jmb.2020.07.023.

55. Jørgensen,S., Elvers,I., Trelle,M.B., Menzel,T., Eskildsen,M., Jensen,O.N., Helleday,T., Helin,K. and Sørensen,C.S. (2007) The histone methyltransferase SET8 is required for S-phase progression. J. Cell Biol., 10.1083/jcb.200706150.

56. An,Z., Chen,Y., Koomen,J.M. and Merkler,D.J. (2012) A mass spectrometry-based method to screen for α-amidated peptides. Proteomics, 10.1002/pmic.201100327.

57. Hengel,S.M., Shaffer,S.A., Nunn,B.L. and Goodlett,D.R. (2009) Tandem Mass Spectrometry Investigation of ADP-ribosylated Kemptide. J. Am. Soc. Mass Spectrom., 10.1016/j.jasms.2008.10.025.

58. Huen,M.S.Y., Sy,S.M.H., Van Deursen,J.M. and Chen,J. (2008) Direct interaction between SET8 and proliferating cell nuclear antigen couples H4-K20 methylation with DNA replication. J. Biol. Chem., 10.1074/jbc.C700242200.

59. Girish,T.S., McGinty,R.K. and Tan,S. (2016) Multivalent interactions by the set8 histone methyltransferase with its nucleosome substrate. J. Mol. Biol., 10.1016/j.jmb.2016.02.025.

60. Wang,C., Qu,C., Alippe,Y., Bonar,S.L., Civitelli,R., Abu-Amer,Y., Hottiger,M.O. and Mbalaviele,G. (2016) Poly-ADP-ribosylation-mediated degradation of ARTD1 by the NLRP3 inflammasome is a prerequisite for osteoclast maturation. Cell Death Dis., 10.1038/cddis.2016.58.

61. Hu,K., Wu,W., Li,Y., Lin,L., Chen,D., Yan,H., Xiao,X., Chen,H., Chen,Z., Zhang,Y., et al. (2019) Poly (ADP -ribosyl)ation of BRD 7 by PARP 1 confers resistance to DNA -damaging chemotherapeutic agents . EMBO Rep., 10.15252/embr.201846166.

62. Wang,Y., Xu,Z.D., Mao,J.H., Hsieh,D., Au,A., Jablons,D.M., Li,H. and You,L. (2015) PR-Set7 is degraded in a conditional Cul4A transgenic mouse model of lung cancer. Chinese J. Lung Cancer, 10.3779/j.issn.1009-3419.2015.06.15.

63. Pannetier M, Julien E, Schotta G, Tardat M, Sardet C, Jenuwein T, Feil R. PR-SET7 and SUV4-20H regulate H4 lysine-20 methylation at imprinting control regions in the mouse. EMBO Rep. 2008 Oct;9(10):998–1005. doi: 10.1038/embor.2008.147. Epub 2008 Aug 22. PMID: 18724273; PMCID: PMC2525564.

64. Hori T, Shang WH, Toyoda A, Misu S, Monma N, Ikeo K, Molina O, Vargiu G, Fujiyama A, Kimura H, Earnshaw WC, Fukagawa T. Histone H4 Lys 20 monomethylation of the CENP-A nucleosome is essential for kinetochore assembly. Dev Cell. 2014 Jun 23;29(6):740–9. doi: 10.1016/j.devcel.2014.05.001. PMID: 24960696; PMCID: PMC4081567.

65. Ke,Y., Han,Y., Guo,X., Wen,J., Wang,K., Jiang,X., Tian,X., Ba,X., Boldogh,I. and Zeng,X. (2017) PARP1 promotes gene expression at the post-transcriptiona level by modulating the RNA-binding protein HuR. Nat. Commun., 10.1038/ncomms14632.

66. Wang,Y., Guo,Y., Tang,C., Han,X., Xu,M., Sun,J., Zhao,Y., Zhang,Y., Wang,M., Cao,X., et al. (2019) Developmental Cytoplasmic-to-Nuclear Translocation of RNA-Binding Protein HuR Is Required for Adult Neurogenesis. Cell Rep., 10.1016/j.celrep.2019.10.127.

67. Krishnakumar,R. and Kraus,W.L. (2010) PARP-1 Regulates Chromatin Structure and Transcription through a KDM5B-Dependent Pathway. Mol. Cell, 10.1016/j.molcel.2010.08.014.

68. Kassner,I., Barandun,M., Fey,M., Rosenthal,F. and Hottiger,M.O. (2013) Crosstalk between SET7/9-dependent methylation and ARTD1-mediated ADP-ribosylation of histone H1.4. Epigenetics and Chromatin, 10.1186/1756-8935-6-1.

69. Le May,N., Iltis,I., Amé,J.C., Zhovmer,A., Biard,D., Egly,J.M., Schreiber,V. and Coin,F. (2012) Poly (ADP-Ribose) Glycohydrolase Regulates Retinoic Acid Receptor-Mediated Gene Expression. Mol. Cell, 10.1016/j.molcel.2012.09.021.

70. Guetg,C., Scheifele,F., Rosenthal,F., Hottiger,M.O. and Santoro,R. (2012) Inheritance of Silent rDNA Chromatin Is Mediated by PARP1 via Noncoding RNA. Mol. Cell, 10.1016/j.molcel.2012.01.024.

71. De Vos,M., El Ramy,R., Quénet,D., Wolf,P., Spada,F., Magroun,N., Babbio,F., Schreiber,V., Leonhardt,H., Bonapace,I.M., et al. (2014) Poly(ADP-ribose) polymerase 1 (parp1) associates with E3 ubiquitin-protein ligase UHRF1 and modulates UHRF1 biological functions. J. Biol. Chem., 10.1074/jbc.M113.527424.

72. Caiafa,P., Guastafierro,T. and Zampieri,M. (2009) Epigenetics: poly(ADP-ribosyl)ation of PARP-1 regulates genomic methylation patterns. FASEB J., 10.1096/fj.08-123265.

73. Shoaib,M., Chen,Q., Shi,X. et al. Histone H4 lysine 20 mono-methylation directly facilitates chromatin openness and promotes transcription of housekeeping genes. Nat Commun 12, 4800 (2021). https://doi.org/10.1038/s41467-021-25051-2

74. Kapoor-Vazirani,P. and Vertino,P.M. (2014) A dual role for the histone methyltransferase PR-SET7/SETD8 and histone H4 lysine 20 monomethylation in the local regulation of RNA polymerase II pausing. J. Biol. Chem., 10.1074/jbc.M113.520783.

75. DaRosa, P.A., Wang, Z., Jiang, X., et al. (2015) Allosteric activation of the RNF146 ubiquitin ligase by a poly(ADP-ribosyl)ation signal. Nature. 517(7533):223–226. doi:10.1038/nature13826

76. Wu, J., Qiao, K., Du, Y. et al. (2020) Downregulation of histone methyltransferase SET8 inhibits progression of hepatocellular carcinoma. Sci Rep 10, 4490. https://doi.org/10.1038/s41598-020-61402-7

77. Yang, J., Yan, R., Roy, A., Xu, D., Poisson, J., & Zhang, Y. (2015). The I-TASSER Suite: protein structure and function prediction. Nature methods, 12(1), 7–8.

78. Prilusky, J., Felder, C., Zeev-Ben-Mordehai, T. et al. (2005) FoldIndex©: a simple tool to predict whether a given protein sequence is intrinsically unfolded, Bioinformatics, Volume 21, Issue 16, Pages 3435–3438. https://doi.org/10.1093/bioinformatics/bti537

79. Xue, B., Dunbrack, R. L., Williams, R. W., Dunker, A. K., & Uversky, V. N. (2010). PONDR-FIT: a meta-predictor of intrinsically disordered amino acids. Biochimica et Biophysica Acta (BBA)-Proteins and Proteomics, 1804(4), 996–1010.

80. Chowdhury, Sourav, Dwipanjan Sanyal, Sagnik Sen, Vladimir N. Uversky, Ujjwal Maulik, and Krishnananda Chattopadhyay. 2019. “Evolutionary Analyses of Sequence and Structure Space Unravel the Structural Facets of SOD1” Biomolecules 9, no. 12: 826. https://doi.org/10.3390/biom9120826

81. Kong, B., Zhou, L. and Liu, W., 2012, January. Improved modularity based on Girvan-Newman modularity. In 2012 Second International Conference on Intelligent System Design and Engineering Application (pp. 293–296). IEEE.

